# Learning the spatial cell-cell communication network to decode multi-channel signaling and predict network-hub vulnerabilities with MOSANIC

**DOI:** 10.64898/2026.07.05.736582

**Authors:** Debraj Das, Pralay Mitra

## Abstract

Intercellular signaling governs the central decisions of tissue biology, from proliferation and immune recruitment to metabolic adaptation and cell death, and its dysregulation is a hallmark of disease. What matters most are properties of the signaling network as a whole rather than of individual interactions: the hubs that hold the network together and mark rational points of intervention, the relays through which a signal propagates across intermediate cells to reach partners it does not directly contact, the response of tissue-wide communication to the loss of a single node, and the metabolite-mediated axis that operates alongside secreted-protein signalling. Scoring known ligand-receptor pairs yields a ranked interaction list that captures none of these and excludes metabolite signalling. We present MOSANIC (**M**ulti-m**O**dal **S**elf-**A**ttention **N**etwork for **I**ntercellular **C**ommunication), which learns a tissue’s communication network directly from spatial transcriptomics. MOSANIC represents each tissue as a single heterogeneous graph of cells, genes and metabolites, initialises every node with a frozen foundation-model representation (scVI, ESM-2 and ChemBERTa), and propagates these representations through a self-attention network over biologically typed edges. Supervision is restricted to spatial gene-expression prediction and excludes ligand-receptor annotation, rendering the inferred communication statistically independent of the reference databases used for evaluation. Across five spatial datasets spanning three platforms and two species, MOSANIC attains the highest accuracy on all eight independent ligand-receptor benchmarks (mean AUROC 0.756) against nine established methods, resolves a metabolite-receptor channel statistically orthogonal to the peptide channel, and reconstructs multi-step signal relays that concentrate within a compact rich-core of load-bearing hub genes and cells whose removal fragments the network far beyond a degree-preserving null. In-silico knockout of these hubs recovers experimentally reported phenotypes, and, given no prior oncological input, MOSANIC nominates SCARF1 as a previously unrecognised communication hub in breast cancer whose elevated expression predicts significantly worse survival in an independent cohort after adjustment for tumour stage and age (hazard ratio 1.17, P = 0.043). MOSANIC is released as an open-source Python package (*mosanic-ccc*).

## 1 Introduction

Tissues are not mere collections of cells; rather represent interactions among them. Each fate decision a cell makes, whether to proliferate, differentiate, attract immunological support, alter metabolism, or undergo apoptosis is determined through communication with its spatial neighbours, and when these interactions cease, the repercussions are profound^1^. Tumour-stroma signaling alters medication sensitivity in pancreatic adenocarcinoma; microglia-neuron interaction regulates neurodegeneration; tumour-associated macrophages determine immunotherapy efficacy. Spatial transcriptomics platforms (10x Visium, Xenium, Slide-seqV2) now record gene expression alongside cellular coordinates at sub-tissue resolution, and a parallel wave of single-cell and spatial foundation models like Geneformer^2^, scGPT^3^, scFoundation^4^, Nicheformer^5^ has shifted the field from task-specific pipelines toward learned representations that transfer across tissues and assays.

Cell-cell communication (CCC) inference has lagged this transition. The two dominant families of methods both predate the foundation-model wave. Co-expression and database matching approaches (CellPhoneDB^6^, CellChat^7^; NicheNet^8^, LIANA+^9^) score ligand-receptor (LR) pairs from a curated catalogue against scRNA-seq vocabularies using permutation tests; they are principled and interpretable, but either conservative towards single modality or not fully utilizing the capacity of single-cell resolution. Spatial-statistical methods (SpatialDM^10^, HoloNet^11^; COMMOT^12^; SpaTalk^13^) inject spatial proximity through Moran’s I, dense LR-cell tensors, or optimal transport; they recover cluster-level signaling and per-pair summary statistics but lack producing direct actionable clinical applicability. The most recent graph-learning entrants (HoloNet; CellNEST^14^) train graph attention networks to score LR pairs, but learn one channel at a time and remain bound to their training knowledgebases.

Three structural limits persist across all three families. First, CCC remains single-channel: most of the published method scores secreted-protein LR pairs (except LIANA+ that allows inferred metabolic communications) ignoring the explicit modelling of metabolite-receptor (MR) layer that carries amino-acid, lipid, and nucleotide signals central to tumour metabolism, neurotransmission, and stromal cross-talk, even though metabolite-receptor databases (MEBOCOST^15^) and single-cell flux estimators (scFEA^16^, Compass^17^) are now mature. Second, outputs are method-specific summaries: each tool returns its own LR ranking but no per-gene quantity that can be re-used to identify hubs, or driver cells in, trace multi-hop relay chains, or predict in-silico knockout effects do for transcription factors). Third, scoring is rigidly database-bound: predictions are constrained to curated LR pairs.

These limits share a single root: CCC has been posed as a labelling problem over a fixed vocabulary of pairs, rather than a representation-learning problem on a typed graph. Resolving all three at once calls for a model that reasons jointly over protein, metabolite and spatial geometry, that learns from the biology of a tissue rather than from ligand-receptor labels, and that exposes one principled per-gene quantity capable of serving every downstream task. The heterogeneous graph transformer^18^ offers a natural substrate for this, provided its typed attention is adapted to the node and edge biology of intercellular communication.

We introduce MOSANIC (**M**ulti-m**O**dal **S**elf-**A**ttention **N**etwork for **I**ntercellular **C**ommunication), a heterogeneous graph transformer built on exactly this substrate. MOSANIC represents a tissue as three interacting node types, cells, genes and metabolites, joined by seven biologically typed edges: three bidirectional cell-cell channels (secreted, metabolite-mediated and intracellular) and four directed cross-type channels (cell to gene, ligand to receptor, cell to metabolite, and metabolite to receptor). Each node enters with a frozen embedding drawn from a domain foundation model, scVI for cells^19^, ESM-2 for gene products^20^ and ChemBERTa for metabolites^21^, so that protein and chemical priors inform communication inference from the outset. Two stacked encoder blocks propagate signal along all seven typed attention lanes in parallel, and the network is trained end to end on a single objective: prediction of held-out spatial gene expression under spatial k-means cross-validation. No ligand-receptor or metabolite-receptor label is ever presented during training. The five analyses that follow, niche clustering, ligand-receptor and metabolite-receptor pair discovery, a per-gene hub-score with channel-restricted variants, multi-hop relay detection, and in-silico knockout by attention-edge ablation, all emerge as parameter-free read-outs of this single learned attention field. Across five spatial transcriptomic datasets spanning three platforms and two species, MOSANIC ranks first on all eight independent LR-database AUROC evaluations against nine baseline methods (mean 0.756, range 0.726–0.822), resolves an MR channel invisible to single-channel methods, traces multi-hop signal relays through a coherent rich-core communication backbone, and validates that backbone through in-silico knockouts whose predictions match independent published evidence on 23 of 24 gene-evidence axes (eight hub genes × three axes; Supplementary Note ??), nominating SCARF1 as a putative drug-actionable multiplexer hub in breast cancer. MOSANIC is distributed as an open-source Python package (*mosanic-ccc*) with an end-to-end pipeline from raw spatial data to interpretable communication maps. By unifying multi-channel singnaling, parameter-free read-outs, and one principled hub-score, MOSANIC reframes CCC inference from rigid database matching toward representation-learning of tissue conversations, and positions cell-cell communication inside the broader spatial multi-omic foundation-model wave.

## 2 Results

### 2.1 Overview of the MOSANIC method

Existing CCC methods score ligand–receptor (LR) pairs against a curated catalogue on a single signaling channel, and return per-method scores that do not transfer to communication-hub identification, multi-hop relay detection or *in-silico* perturbation, leaving the metabolite-receptor (MR) channel structurally invisible. We developed MOSANIC (Multi-mOdal Self-Attention Network for Intercellular Communication), a heterogeneous graph transformer that instead learns tissue communication as representation on a typed multi-omic graph (Fig. 1).

**Fig. 1.**
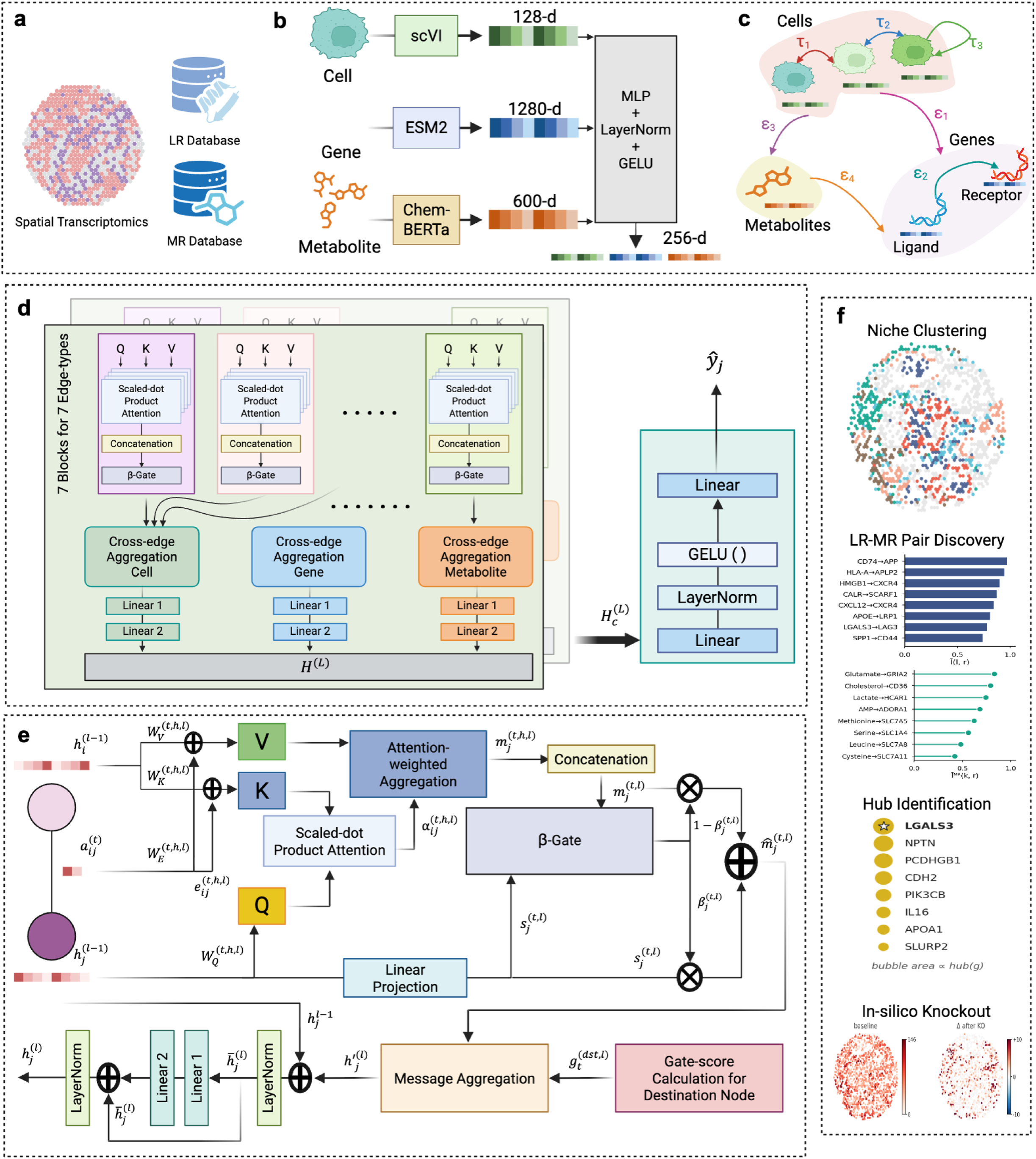
MOSANIC: a heterogeneous graph transformer for multi-channel cell–cell communication inference from spatial transcriptomics. a–c · Inputs and graph construction. **a,** MOSANIC consumes a spatial-transcriptomics dataset (cells with coordinates and counts) together with two reference databases: a ligand–receptor (LR) database (CellNEST, 14,909 pairs) and a metabolite–receptor (MR) database (MEBOCOST, 794 human / 790 mouse pairs). b, Node features come from frozen pre-trained foundation models, namely scVI (128-d) for cells, ESM-2 650M (1,280-d) for genes and ChemBERTa-77M (600-d) for metabolites, and are projected through a shared MLP + LayerNorm + GELU stack to a common 256-d latent (Eq. 1). c, The resulting heterogeneous graph carries three node types (cell, gene, metabolite) and seven edge types: three cell–cell τ edges (τ₁ secreted, τ₂ metabolite-mediated, τ₃ intracellular) and four cross-type ε edges (ε₁ cell→gene, ε₂ ligand→receptor, ε₃ cell→metabolite, ε₄ metabolite→receptor). LR pair scores are read off ε₂ attention; MR scores off ε₄. **d, e · MOSANIC encoder. d,** Each block runs seven parallel typed attention lanes (one per edge type); per-lane β-gated residuals (Eq. 7) feed a per-destination-node-type softmax gate (Eq. 8) and a two-layer FFN (Eq. 10). Two blocks are stacked; a Linear → GELU → LayerNorm → Linear decoder predicts held-out expression ŷ_j (Eq. 12). e, Zoom of one lane on source i / destination j: K/Q/V projections, scaled-dot-product attention α_ij^(t,h) (Eq. 3), attention-weighted aggregation m_j^(t,h) (Eq. 4), β-gated residual (Eqs. 5–7), and LayerNorm (Eq. 9). **f · Downstream applications.** Learned attention drives five read-outs (zero added parameters): niche clustering on hi(L) (§5.1; Fig. 3); LR pair discovery via intensity-weighted scoring Ī(l,r) (Eq. 15; Fig. 2); MR pair discovery via Ī^MR(k,r) (Eq. 17; Fig. 4); per-gene hub-score hub(gC) (Eq. 18, channel mask Cε2,ε4 resolves Hub / LR-network hub / MR hub naming; Fig. 5); and *in-silico* knockout by attention-edge ablation (Eq. 21; Fig. 6).

From spatial transcriptomics with cell coordinates (Fig. 1a), MOSANIC builds a single graph on three node types, cells, genes and metabolites, each initialised with a frozen foundation-model embedding: scVI for cell expression, ESM-2 for protein sequence and ChemBERTa for metabolite structure (Fig. 1b)^19–21^. Seven biologically typed edges connect them consisting three bidirectional cell–cell channels (τ₁ secreted, τ₂ metabolite-mediated, τ₃ intracellular) and four directed cross-type channels (ε₁ cell→gene, ε₂ ligand→receptor, ε₃ cell→metabolite, ε₄ metabolite→receptor), placing spatial structure and molecular vocabulary in one object (Fig. 1c). A two-block encoder that combines the typed attention^22^ with the heterogeneous graph transformer^18^ runs all seven lanes in parallel (Fig. 1d, e), trained on a single objective of predicting of held-out spatial expression, and never on LR or MR labels, so evaluation against independent catalogues stays non-circular (Fig. 1f).

Every downstream analysis is then a deterministic read-out of the learned attention, computed with no added parameters. Five are reported: niche clustering on cell embeddings; intensity-weighted LR and MR pair discovery from ε₂ and ε₄ attention; a per-gene hub-score with canonical, LR-only and MR-only variants; multi-hop relay detection; and *in-silico* knockout by ε-edge ablation.

### 2.2 MOSANIC learns a reliable, non-random communication-network architecture

Before using MOSANIC’s learnt communications architecture as a substrate for later analyses, we investigated whether its ligand–receptor scores agree with independent reference evidence and whether the resulting network has non-random topology. Figure 2 establishes both, accuracy across multiple databases and eight baselines and a network that survives strict null comparison.

**Fig. 2.**
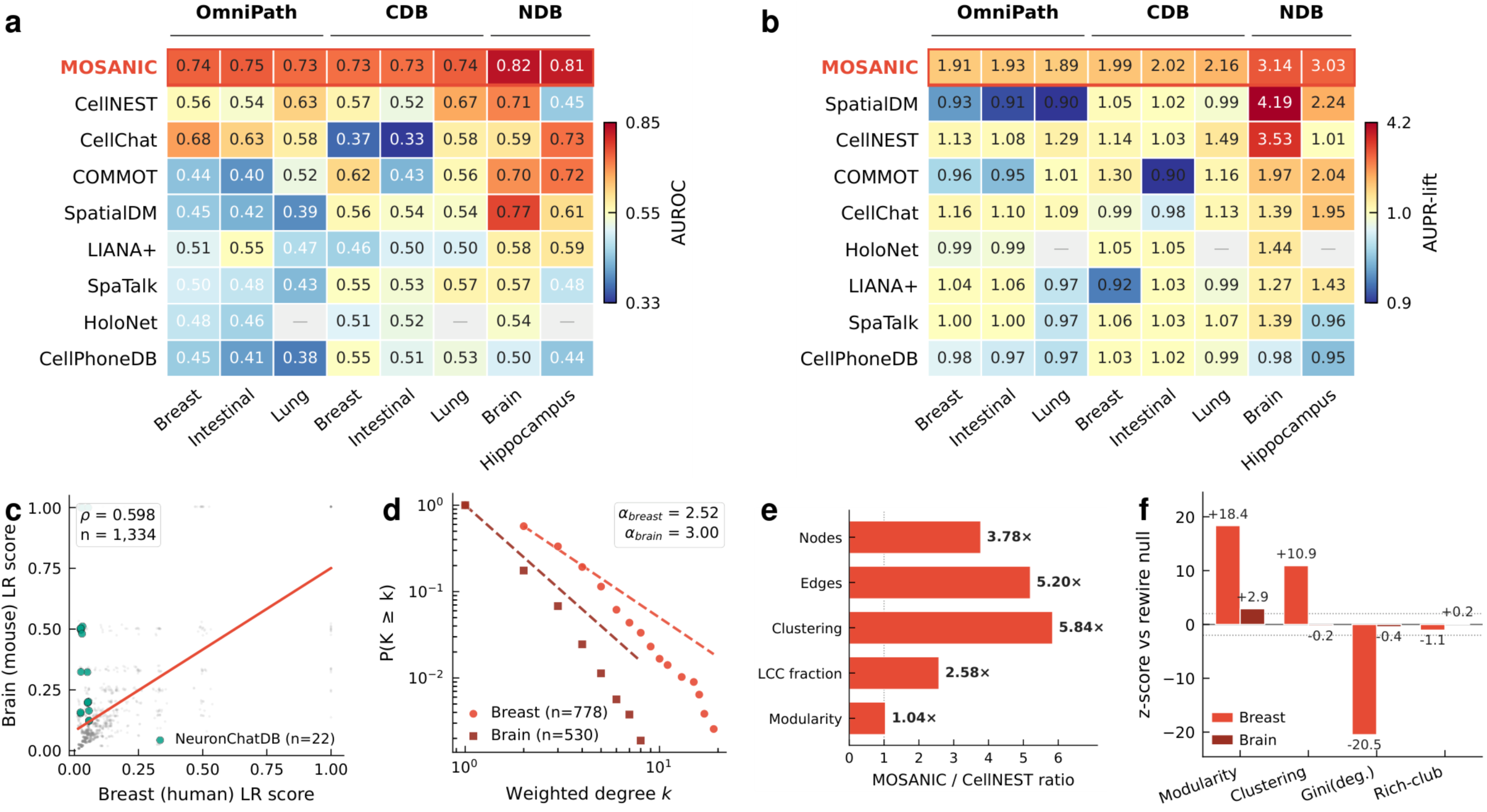
MOSANIC produces reliable LR scores and a non-random communication-network architecture. **a**, AUROC across nine methods (rows) × eight held-out database × dataset evaluations: OmniPath (6,555 pairs) and ConnectomeDB2025 (3,550 pairs) on 3 human datasets; NeuronChatDB (452 pairs) on 2 mouse datasets. Rows sorted by mean; MOSANIC outlined red. Dashes = inapplicable (e.g., HoloNet OOM). **b**, AUPR-lift (observed/random hit-rate) on the same eight evaluations; values > 1 = enrichment. MOSANIC is the only method > 1 in every column. **c**, Cross-species conservation of MOSANIC LR scores: 1,334 pairs scored in both breast and brain (Spearman ρ = 0.598, *P* = 4.9 × 10⁻¹³⁰). Teal dots = 22 pairs validated in NeuronChatDB. **d**, Heavy-tailed weighted-degree distribution of the top-10 % LR-attention graph (log–log). Power-law exponents α_breast = 2.52, α_brain = 3.00. **e**, MOSANIC vs CellNEST topology fold-changes (Nodes 3.78×, Edges 5.20×, Clustering 5.84×, LCC 2.58×, Modularity 1.04×). **f**, Z-scores vs Maslov–Sneppen rewire-null (100 graphs); |z| > 2 rejects degree-sequence explanation. Source data: Supplementary Tables T1, T2; details: Supplementary Figs. S1–S7.

MOSANIC ranks validated database pairs above unscored pairs on every dataset–database combination we tested. Across five spatial transcriptomic datasets and three independent reference ligand-receptor databases, OmniPath (6,555 pairs)^23^, ConnectomeDB2025 (3,550 pairs) and NeuronChatDB (452 pairs)^24^, MOSANIC achieves the top AUROC on all eight evaluations (mean 0.756, range 0.726–0.822; Fig. 2a) against eight baseline methods: CellChat^7^, CellPhoneDB^6^, LIANA+^9^, SpatialDM^10^, SpaTalk^13^, COMMOT^12^, HoloNet^11^ and CellNEST^14^. The advantage over the second-best method averages +0.099 (Cohen’s d = 2.35; Wilcoxon signed-rank P = 0.004; per-evaluation AUROC and AUPR/AUPR-lift values for all eight methods in Supplementary Tables T1 and T2). Orthogonal functional validation against 104 NicheNet^8^ ligand-perturbation experiments derived from independent GEO transcriptomic perturbation data gives MOSANIC LigAct AUROCs of 0.630–0.644 on the three human datasets (Supplementary Fig. S1). Per-evaluation mean rank and paired-advantage forest plots are in Supplementary Figs. S2 and S3.

As methods score vocabularies of different sizes, we also report AUPR-lift, the ratio of observed AUPR to the random-baseline AUPR implied by each method’s hit rate. MOSANIC is the only method with AUPR-lift > 1 on every evaluation (range 1.89–3.14; Fig. 2b). Vocabulary size does not provide advantage for higher AUROC (Supplementary Fig. S4a–c), and the MOSANIC advantage persists across top-k cutoffs from 10 to 1,000 (Supplementary Fig. S5a, Lift@100 across the three databases; Supplementary Fig. S5b, Lift@K curves across human datasets). Cross-species conservation of LR scores between human breast and mouse brain is strong (Spearman ρ = 0.598; P = 4.9 × 10⁻¹³⁰; Fig. 2c), with NeuronChatDB-validated pairs preferentially placed at higher joint scores.

The communication graph MOSANIC produces is also non-random. At top-10% attention stringency the LR-attention graph is heavy-tailed (power-law exponents α_breast_ = 2.52, α_brain_ = 3.00; Fig. 2d)^25^ suggesting a topology associated with biological information-processing networks^26^. At the same threshold, the breast network is 3.78-fold larger in nodes and 5.84-fold higher in clustering than CellNEST^14^, the only peer method that likewise exposes an attention-weighted LR graph (Fig. 2e). MOSANIC’s normalised percolation ratio is 0.146, the lowest across all baselines (range 0.181–0.332; Supplementary Fig. S6a breast, S6b brain). Against 100 Maslov–Sneppen rewire-null graphs^27^ preserving degree exactly, modularity exceeds the null by 18.4 σ and weighted clustering by 10.9 σ (Fig. 2f); the brain graph behaves identically (Supplementary Fig. S7).

Further, MOSANIC ranks all six FDA-target immune-checkpoint LR interactions in the top 4% of its breast vocabulary (96th–99th percentile by intensity-weighted score), while CellChat and CellNEST score none of them (Supplementary Fig. S8). These findings help us to quantify a single terminology used throughout the following sections, the hub-score for each gene (g∣C), defined as the sum of incoming attention into the gene over the channel mask C⊆{ε_2_,ε_4_}, averaged over heads and transformer layers (defined formally as Eq. (18) in Methods). Establishing the network is reliable, non-random, and parameterised by a single principled hub-score, we next ask whether the topology encodes biology without supervision.

### 2.3 MOSANIC’s communication graph encodes tissue biology without supervision

Despite the fact that the communication graph that MOSANIC learns is dependable and does not follow a random pattern (Fig. 2), it is essential for the topology to encode tissue biology, and it is also important to determine whether or not it does so spontaneously. Figure 3 assembles four independent tests: spatial niches that emerge from the cell embedding alone; niche-level communication programmes that re-organise without programme labels; canonical cell-type axes that surface from attention; and a hub set partly conserved across five spatial transcriptomic datasets.

**Fig. 3.**
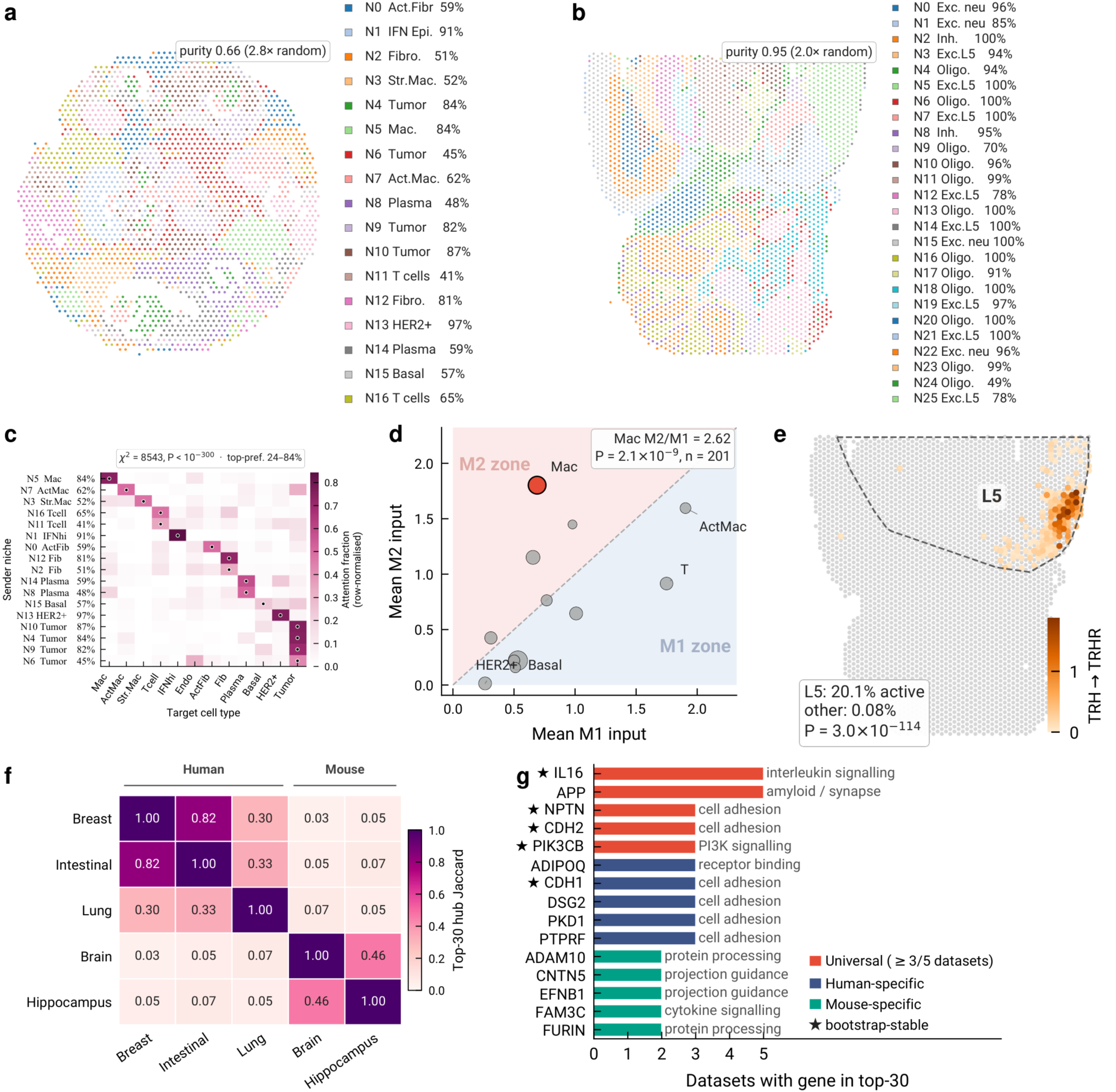
MOSANIC’s communication graph encodes tissue biology without supervision. **a, b**, MOSANIC niches on the learned cell embedding (k-means; *k* = 17 breast, 26 cortex). Cell-type purity of the niches (mean over niches of the largest single-cell-type fraction): breast 0.66 and cortex 0.95, respectively 2.8× and 2.0× the purity of a random partition of the same granularity (breast 0.24, cortex 0.48; 1,000 permutations). No cell-type labels given during training. **c**, Row-normalised outgoing attention from each breast niche (rows) to target cell types (columns); rows grouped by dominant own cell type. χ² = 8,543, *P* < 10⁻³⁰⁰; top-preferred fraction 24–84%. **d**, M1 vs M2 input per breast cell type; dot size proportional to count. Macrophages: M2/M1 = 2.62 (Wilcoxon *P* = 2.1 × 10⁻⁹, *n* = 201). **e**, Mouse cortex coloured by TRH→TRHR attention; dashed polygon outlines Exc. L5. 20.1% of L5 cells active vs 0.08% elsewhere (Kruskal–Wallis *P* = 3.0 × 10⁻¹¹⁴). **f**, Pairwise top-30 hub Jaccard across five datasets. **g**, Universal, human-specific and mouse-specific hubs with GO Biological Process term (Enrichr v2); stars mark bootstrap-stable hubs (frequency ≥ 0.9 over 100 resamples). Source data are provided with this paper.

MOSANIC’s cell embedding partitions tissue into spatial niches that recover known cell-type compartments without supervision. For breast cancer (n = 2,516 spots), k-means with k = 17 yields niches with cell-type purity 0.656, which is 2.8-times better than the 0.236 expected for a random partition of the same granularity (1,000 permutations; Fig. 3a; see clustering parameters in Supplementary Note N2); niche N13 is 97% HER2⁺ luminal epithelial, N5 is 84% macrophage, and N1 is 91% interferon-high epithelial. These compartments are in accordance with the conventional architecture of the breast cancer microenvironment, which has been demonstrated in earlier research^28,29^. In mouse cortex (n = 3,342 spots), 26 niches reach purity 0.949 (2.0-times random), sub-dividing the coarse 5-class annotation into cortical-depth gradients, recovering the laminar L1–L6 organisation established by prior research works^30,31^ (Fig. 3b). The phenomenon generalises to the intestinal, lung and hippocampus datasets (Supplementary Fig. S9), with the niche-purity-versus-random comparison summarised in Supplementary Fig. S10.

In addition to being spatial groupings, niches are also characterised by the fact that each one targets a distinct group of communication partners. The outgoing secreted-ligand attention of MOSANIC was aggregated for each cell based on the cell-type identification of its targets, and then the results were row-normalized within the niche framework. The resulting 17 × 12 niche × target-cell-type matrix is highly structured for Breast data (χ² = 8,543, P < 10⁻³⁰⁰; Fig. 3c). The top-preferred target captures 24–84% of outgoing attention per niche, and the preferred target is biologically interpretable: macrophage-dominated niche N5 directs 74% of outgoing attention to other macrophages; HER2⁺ niche N13 directs 76% to HER2⁺ targets; T-cell niche N16 directs 48% to T cells and a further 22% to fibroblasts (Supplementary Table T3).

Beyond niche identity, MOSANIC recovers canonical molecular axes without supervision. In the breast tumour microenvironment, macrophages (n = 201) receive 2.62-fold higher summed input from four canonical M2 LR pairs than from six M1 pairs (Wilcoxon signed-rank P = 2.1 × 10⁻⁹; Fig. 3d), recovering the foundational M1/M2 polarisation axis^32,33^. In mouse cortex, the TRH→TRHR pair is active in 20.1% of layer-5 excitatory neurons and 0.08% elsewhere (Kruskal–Wallis P = 3.0 × 10⁻¹¹⁴; Fig. 3e), recapitulating documented layer-5 TRH specificity without explicit layer annotation^34^. See extended tests in Supplementary Fig. S11.

The same hub architecture re-emerges in every dataset we tested. Pairwise Jaccard overlap of the top-30 hub-score genes across five datasets reaches 0.48 within human cancers and 0.46 within mouse neural tissues, falling to 0.07 across species (Fig. 3f). Seventeen genes occupy the top 30 of at least three of five datasets (full annotated list in Supplementary Table T4); the five most recurrent genes IL16, APP, NPTN, CDH2 and PIK3CB span interleukin chemoattraction, amyloid-precursor and synaptic biology, cell adhesion, and PI3K signaling (*Enrichr* GO Biological Process; Fig. 3g). These hubs are bootstrap-stable across 1,000 edge-resamples of each dataset’s top-10 % graph (Supplementary Fig. S12). The corresponding LR-pair vocabulary is largely tissue-specific (Supplementary Fig. S13), so the conservation emerges at the gene level, paralleling the observation in regulatory-network biology that hub regulators are more conserved than the programmes they drive^35^.

Hence, MOSANIC’s communication network recovers tissue niches, niche-level communication programmes, canonical molecular axes, and a partially conserved hub set, none of which were provided as labels. Thus far, we have exclusively utilised the secreted-protein channel; we now enquire whether MOSANIC’s metabolite layer targets a similar yet partially distinct receptor population.

### 2.4 MOSANIC’s architecture is multi-channel

The architecture depicted in Figures 2 and 3 utilised solely the secreted-protein channel. Metabolite-mediated signaling like lactate–GPR81 in tumour immune evasion^36^, kynurenine–AhR in immunosuppression^37^, and cholesterol–CD36 in tumor-associated macrophage (TAM) polarisation^38^ etc. remains structurally undetectable by prior spatial-CCC methods due to their restricted vocabulary. In this section, we investigate whether MOSANIC’s heterogeneous graph genuinely resolves a second, parallel metabolite-receptor (MR) channel via MEBOCOST^15^ vocabulary and scFEA^16^ flux, and whether that channel adds biology the protein channel alone cannot attain.

The MR layer is rank-coupled to the LR layer at the metabolite level but not at the gene level. Most metabolites are closely aligned with the y = x diagonal (Spearman ρ = 0.54 breast, 0.57 intestinal, 0.55 lung, 0.51 brain, 0.55 hippocampus; all P < 10⁻²; Fig. 4a shows breast and brain, coloured by KEGG class). The off-diagonal cases Putrescine, Xanthine, Tyrosine, Choline possess the metabolite-channel biology that the protein channel does not perceive at the same intensity. The metabolic substrate is itself spatially structured and cell-type-resolved (Supplementary Fig. S14), and replicates across the remaining four datasets (Supplementary Fig. S15a intestinal, S15b lung, S15c brain, S15d hippocampus). Full per-dataset MR pair rankings are in Supplementary Table T5.

**Fig. 4.**
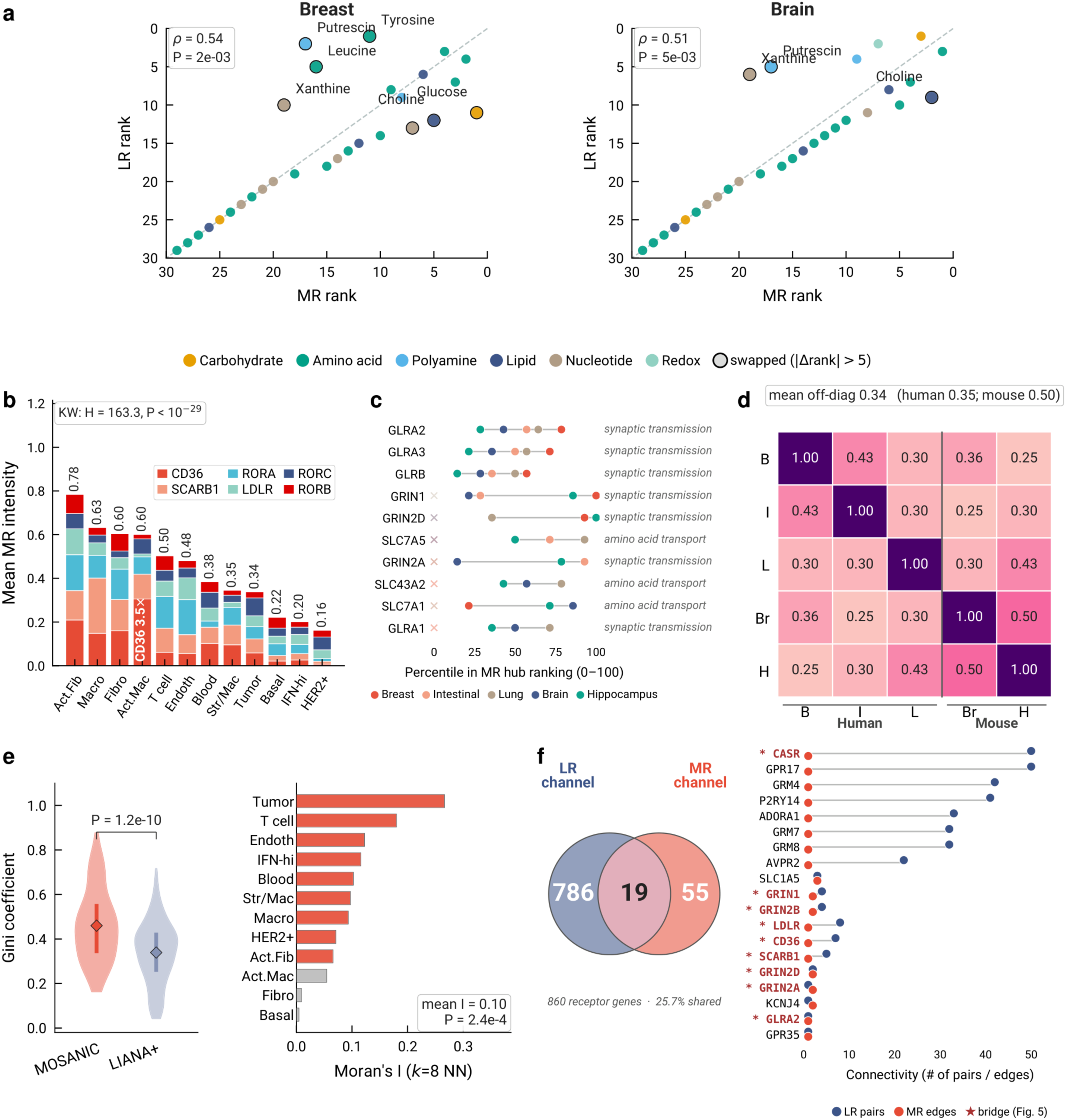
MOSANIC’s architecture is multi-channel. **a**, MR vs LR rank scatter (breast + brain); each dot = one metabolite, coloured by KEGG class. Dashed line y = x; black-edged dots are swapped (|Δrank| > 5). Spearman ρ + *P* per tissue. **b**, Mean cholesterol MR intensity per breast cell type, decomposed into six receptor contributions (CD36, SCARB1, RORA, LDLR, RORC, RORB). Kruskal–Wallis across cell types; activated macrophages use CD36 at 3.5× the rest. **c**, Universal MR hubs (top-20 in ≥ 3 of 5 datasets); each dot = the gene’s percentile in one dataset’s MR ranking. GO term right (Enrichr v2). **d**, Cross-dataset MR hub Jaccard (top-20); diagonal = 1. **e**, MOSANIC vs LIANA+ on the same breast dataset. Left: Gini per MR pair (Mann–Whitney *P*). Right: within-cell-type Moran’s *I* of MOSANIC per-cell MR intensity (8-NN); red bars *P*_sim < 0.05. **f**, Receptor-gene partition. Left: Venn of 860 receptors: 786 LR-only, 19 shared, 55 MR-only. Right: shared receptors ranked by joint connectivity; blue = LR pair count, red = MR edge count. Asterisks = Fig. 5 bridge hubs. Source data are provided with this paper.

Rank-level coupling does not necessarily demonstrate gene-level overlap. Cholesterol, the top MR metabolite in breast is routed through six distinct receptors with thirty-fold cell-type variation (Kruskal–Wallis H = 163.3, P < 10⁻²⁹; Fig. 4b); activated macrophages use CD36 at 3.5× the mean of the rest, recovering the CD36–cholesterol axis through which TAMs accumulate cholesterol from the tumour microenvironment^38,39^. Cholesterol flux and CD36 expression are spatially coincident at single-cell resolution (Spearman ρ = 0.25, P < 10⁻³⁶; Supplementary Fig. S16), and receptor-specific addressing extends to 44 tumour-versus-immune-differential MR pairs (Supplementary Fig. S17, Supplementary Table T6).

Restricting the hub-score (see Methods) to the ε₄ edges identifies an MR-channel hub upper tail dominated by three families: inhibitory glycine receptors (GLRA2, GLRA3, GLRB)^40^, ionotropic glutamate receptors (GRIN1, GRIN2A–D)^41^, and amino-acid transporters of the SLC7 family (SLC7A1, SLC7A5, SLC43A2)^42^. All ten MR hubs in Fig. 4c sit at the 75–100th percentile of their datasets’ MR ranking; *Enrichr* GO terms are dominated by synaptic transmission and amino-acid transport (Supplementary Note N3). The same hub upper tail recurs across tissues. Pairwise top-20 MR hub Jaccard is 0.43 between breast and intestinal, 0.50 between brain and hippocampus, and 0.34 across species (Fig. 4d). Three receptors (GLRA2, GLRA3, GLRB) occupy the top-20 MR hub list of all five datasets; cortical vs. tumour tissues partition the metabolite vocabulary at the tail (Supplementary Fig. S18).

The MR channel resolves biology that pseudobulk methods structurally cannot. Compared with LIANA+ on the same breast dataset, MOSANIC’s per-pair cell-type Gini is higher (mean 0.46 vs 0.34; Mann–Whitney P = 1.2 × 10⁻¹⁰; Fig. 4e left). MOSANIC’s per-cell MR intensity is spatially autocorrelated within every cell type (mean Moran’s I = 0.10, 8-NN; Wilcoxon P = 2.4 × 10⁻⁴; Fig. 4e right), whereas LIANA+’s pseudobulk score gives I = 0 by construction. The top-20 MR pairs of MOSANIC exhibit a 65% relevance to cancer, in contrast to LIANA+’s 25% relevance (Supplementary Fig. S19; pipeline in Supplementary Note N4).

Most of the MR vocabulary is therefore reserved for ligands the LR channel cannot sense. The receptor-gene partition (Fig. 4f, left) splits 860 receptors into 786 addressed only by the LR channel (ε₂), 55 only by the MR channel (ε₄), and 19 shared between the two. Ranked by joint connectivity (Fig. 4f, right), the shared set is led by GPCRs with broad peptide vocabularies like the calcium-sensing receptor CASR^43^, the orphan-derived GPR17^44^, metabotropic glutamate GRM4^45^ and purinergic P2RY14^46^ and closed by lipid-scavenger and glutamate receptors with focused LR connectivity (CD36, SCARB1, LDLR, GRIN1, GRIN2A–D)^47^. These bridge receptors re-appear in the next figure as the relay-transit genes through which ligand-receptor and metabolite signals converge.

### 2.5 Multi-hop, cross-channel signal relays route through a rich-core communication backbone

The two independent channels i.e. secreted-protein (ε₂) and metabolite-mediated (ε₄) demonstrate information flow across modalities using a relay cell receiving a protein signal and emitting a metabolic one which is an unique capability MOSANIC possesses. Exploiting the independent per-edge attention scores, we enumerate multi-hop relay chains, in which cell A signals cell B through one ligand–receptor pair and B relays to cell C through a different pair, including cross-channel relays where the signal switches from protein to metabolite at the relay hub, a transduction motif architecturally impossible for any single-channel CCC method. Under conservative enumeration criteria (Methods §4.3; Supplementary Note N5) we detect 312 relay events in breast (200 two-hop + 33 three-hop + 79 cross-channel) and 216 in brain; removing the enumeration caps recovers >3× more chains with identical unique relay patterns and validation rates, so the reported counts are lower bounds. These relays are not scattered; they concentrate into a coherent backbone: the largest connected component (LCC) at the top-10% attention threshold contains 318 cells in breast (Fig. 5a) and 724 in mouse cortex (Fig. 5b), with hub cells (top-5% weighted degree) at the geometric centre and secreted-LR and metabolite-MR edges interleaved. The breast inset zooms into the densest hub neighbourhood, a 3 × 3 mm sub-region with 5 hubs, 86 cells and 130 attention edges in one tightly-coupled cluster.

**Fig. 5.**
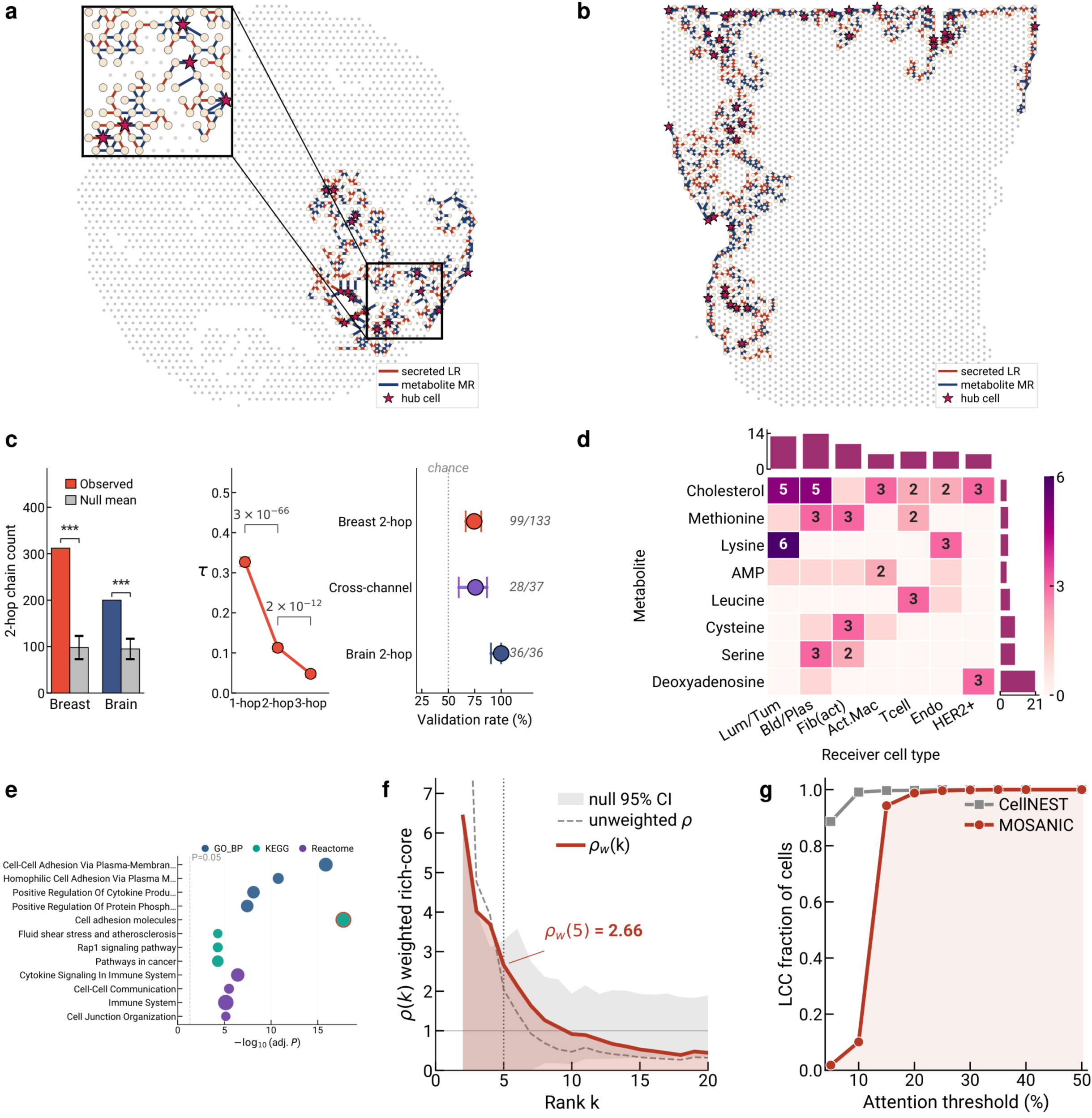
Multi-hop, cross-channel signal relays route through a rich-core communication backbone. **a**, Breast at top-10% attention: LCC (*n* = 318 cells, cream) with hub cells (top-5% weighted degree, *n* = 15, magenta stars); secreted-LR edges dark red, metabolite-MR edges dark blue. Inset (dashed rectangle): densest hub neighbourhood with 5 hubs, 86 cells and 130 attention edges in one tightly-coupled 3 × 3 mm sub-region. **b**, Mouse cortex LCC (*n* = 724 cells, 37 hubs); same colour scheme. **c**, Three-part statistical validation. Left: permutation enrichment of relay-chain count (breast 3.2×, brain 2.1×; *P* < 0.001). Middle: mean τ-score decay across hops (0.327 → 0.113 → 0.048); the value above each bracket is the per-transition Mann–Whitney *P*-value (1→2-hop, *P* = 3 × 10⁻⁶⁶; 2→3-hop, *P* = 2 × 10⁻¹²). Right: PPI + TF intracellular-path validation rate with Wilson 95% CIs (breast 99/133, cross-channel 28/37, brain 36/36; dotted line, 50% chance). **d**, 79 cross-channel cascades by metabolite × receiver cell type (χ² *P* = 1.5 × 10⁻¹⁰). **e**, Top GO BP / KEGG / Reactome enrichments for the 50 hub genes (Enrichr). **f**, Weighted rich-core curve ρ_w(*k*); peak ρ_w(5) = 2.66. **g**, LCC fraction vs attention threshold for MOSANIC and CellNEST. Source data are provided with this paper.

Three complementary tests confirm the relay structure is not explained by spatial proximity alone (Fig. 5c). Permutation of attention across edges yields 3.2-fold (breast) and 2.1-fold (brain) fewer relay chains than observed (P < 0.001 each). Signal strength decays monotonically from a mean τ of 0.327 at one hop to 0.113 at two hops (65 % attenuation) to 0.048 at three hops (Mann–Whitney P = 3 × 10⁻⁶⁶ and 2 × 10⁻¹²; Supplementary Fig. S20), matching sequential paracrine signal transduction study^48^. Most importantly, the relays are intracellularly plausible: querying STRING-derived signaling PPIs^49^ and the TRRUST + DoRothEA TF-target databases^50,51^ (fetched from CellNEST^14^ repository) for a receptor → PPI → TF → ligand path validates 74 % of breast two-hop chains (99/133), 76 % of cross-channel chains (28/37) and 100 % of brain two-hop chains (36/36), recapitulating canonical cascades such as CD163→CSNK2B→NFKB1→IFNG and IFNGR1→STAT1→ARF1 (full paths, TF usage and the receptors that fail PPI coverage are in Supplementary Table T7; method in Supplementary Note N5). Multi-hop relays also extend the spatial reach of communication: three-hop chains span 1,008 µm versus 621 µm for two-hop chains (1.6-fold, P = 1.0 × 10⁻¹⁹). The 79 cross-channel cascades (χ² P = 1.5 × 10⁻¹⁰; Fig. 5d) are the capability unique to a heterogeneous multi-channel graph: each pairs a secreted-LR hop with a metabolite→receptor hop, the relay hub converting a protein signal into metabolic support for its neighbours. They concentrate on cholesterol→CD36→activated-macrophage receivers (28/79, 35%), recapitulating tumour-associated macrophages as cholesterol sinks^38,52^ and on amino-acid transporters (36/79, 46%) linking nutrient sensing to mTOR-driven proliferation^53^. Two cascades illustrate how MOSANIC resolves a protein signal being converted into a metabolic one at the relay hub: in breast, a TNFSF12→CD163 secreted interaction hands off to a Cholesterol→CD36 metabolite interaction, driving M2 macrophage polarisation; and an immune-to-stromal SEMA4D→CD72 interaction hands off to a Deoxyadenosine metabolite output, its validated intracellular path CD72→PTPN6→CXCR4→NT5E/CD73 linking semaphorin signaling to the CD73/adenosine immunosuppressive axis. These cross-channel intracellular paths were validated at 76 % using PPI plus scFEA metabolic module–gene–metabolite mapping (Supplementary Table T8). Routing is 62–64 % heterotypic, with CAFs and TAMs over-represented as the relay-transit centre cell^54,55^; the full sender→relay→receiver cell-type flow and the per-cell-type role specialisation are resolved in Supplementary Figs. S21 and S22. Within this routing structure, SCARF1 is the single most prolific relay-transit hub in breast, receiving one CALR input and fanning out across 48 relay chains (Supplementary Table T9).

Crucially, these relay predictions are not a topological curiosity: independently of the model, the strongest of them match published experimental evidence. CD163 is a direct TWEAK/TNFSF12 receptor that drives macrophage accumulation^56,57^ and is targeted by the clinical antibody enavatuzumab^58^; CALR→SCARF1 is a validated immunogenic-cell-death clearance axis whose receptor, when deleted, causes lupus-like autoimmunity in mice^59,60^; cholesterol→CD36 uptake is required for TAM differentiation^38^; SEMA4D→CD72 feeds the CD73/adenosine axis targeted by 18 clinical monoclonal antibodies^61,62^; and in brain the GNAS→ADCY1 cAMP fan-out and the AGRN synaptic relay recapitulate G-protein and excitatory-synapse biology^63,64^. That the model recovers these axes from attention topology alone, without any pathway or receptor-function annotation, establishes relay chains as a biologically grounded read-out (full evidence and additional axes in Supplementary Note N6). The relay backbone is a rich-core defined by the canonical hub-score hub(*g* ∣ {*ε*_2_, *ε*_4_}) of Eq. (18) (hub genes = top 10 %; hub cells = top 5 % of the per-cell variant, Eq. 19; Methods §4). The 50 hub genes are biologically coherent: strongest enrichment is for cell-adhesion molecules via plasma membrane (adj-P = 1.4 × 10⁻¹⁶), then cytokine production (7.7 × 10⁻⁹) and protein phosphorylation (3.6 × 10⁻⁸), with breast hubs additionally enriching TNF / NF-κB/ immune-checkpoint pathways and brain hubs synaptic transmission (Fig. 5e; Supplementary Table T10). Their topological privilege is quantitative: the weighted rich-core *ρ*_w_(*k*)65 exceeds a degree-preserving rewire-null at every rank k = 2–20 (*ρ*_w_(2) =6.42, *ρ*_w_(5) =2.66; Fig. 5f), the operational signature of rich-core topology^66^.

This rich-core organisation, and the relay communities that run through it, are specific to MOSANIC’s hub-score rather than generic to any cell-level graph. While benchmarking against CellNEST, which introduced relay detection in spatial transcriptomics but on a radius graph 13.4-fold denser. MOSANIC’s LCC fraction drops from 0.94 at top-15 % to 0.02 at top-5 %, while CellNEST stays at 0.88–0.99 (Fig. 5g); CellNEST’s near-fully-connected graph designates 87 % of cells as relay hubs and cannot resolve communities, whereas MOSANIC resolves structured relay communities that are 5.2× (breast) and 7.0× (brain) more spatially compact (Mann–Whitney P < 0.001; Supplementary Fig. S23) and explain communication-gene expression variance on 15/15 canonical genes at η² far above CellNEST (Wilcoxon P = 3.1 × 10⁻⁵; Supplementary Fig. S24). Notably, cross-channel relay detection is architecturally impossible for CellNEST’s single-channel graph.

### 2.6 Hub vulnerabilities partition tissue-specific communication architectures

Network-as-prediction (Figs 2–5) holds significance only if the detected hubs are load-bearing; their removal must disrupt the communication architecture. We performed an in-silico knockout of attention edges linked to each potential hub and quantified the decrease in LR activity (ΔLR), metabolite-receptor activity (ΔMR), and downstream relay-chain integrity (Methods §5.5; Supplementary Note N7).

Ablating the top-4 LR-network hubs in breast tumour produced a monotonically decreasing loss: LGALS3 (3.36 %) > PIK3CB (2.37 %) > NPTN (0.62 %) > APOA1 (0.19 %) (Fig. 6a; baseline LR/MR maps in Supplementary Fig. S25a–d; full top-10 hub ranking per tissue in Supplementary Fig. S26a breast, S26b brain; per-gene magnitudes in Supplementary Table T11). LGALS3 (galectin-3) is an established tumour-microenvironment immunomodulator under active clinical investigation^67,68^, and PIK3CB encodes the p110β catalytic subunit of PI3K, a cancer-genome-validated breast-cancer driver^69^. NPTN (neuroplastin) is a synaptic adhesion molecule with adhesion functions also reported in epithelial and stromal contexts^70^; APOA1 mediates HDL-cholesterol efflux from tumour-associated macrophages^71^. The same procedure in mouse cortex yielded a far steeper collapse; GNAS (10.34 %) > APP (7.97 %) > ADAM10 (1.99 %) > IL16 (1.11 %), with GNAS alone draining 3.1-times the breast leader (Fig. 6b). GNAS encodes the α-subunit of the heterotrimeric stimulatory G-protein, the master upstream activator of adenylyl-cyclase-driven cAMP signaling that dominates cortical neuromodulatory communication^72,73^. APP and ADAM10 are the canonical neuronal amyloid precursor and α-secretase that together control synaptic-cleft adhesion and cleavage^74,75^. Metabolite-channel knockout of the top MR hub per tissue (ADAMTS4, 1.76 %; GRIN1, 7.81 %) reproduced a 4.4× brain/breast disparity (Fig. 6c; alternative MR-receptor breakdowns in Supplementary Fig. S27a–e (breast cholesterol receptors CD36/SCARB1/LDLR) and Supplementary Fig. S28a–f (brain NMDA family beyond GRIN1)). GRIN1 (the obligate NR1 subunit of the NMDA receptor) is the load-bearing glutamatergic gateway of cortical excitation^76,77^, confirming the pattern is not LR-specific.

**Fig. 6.**
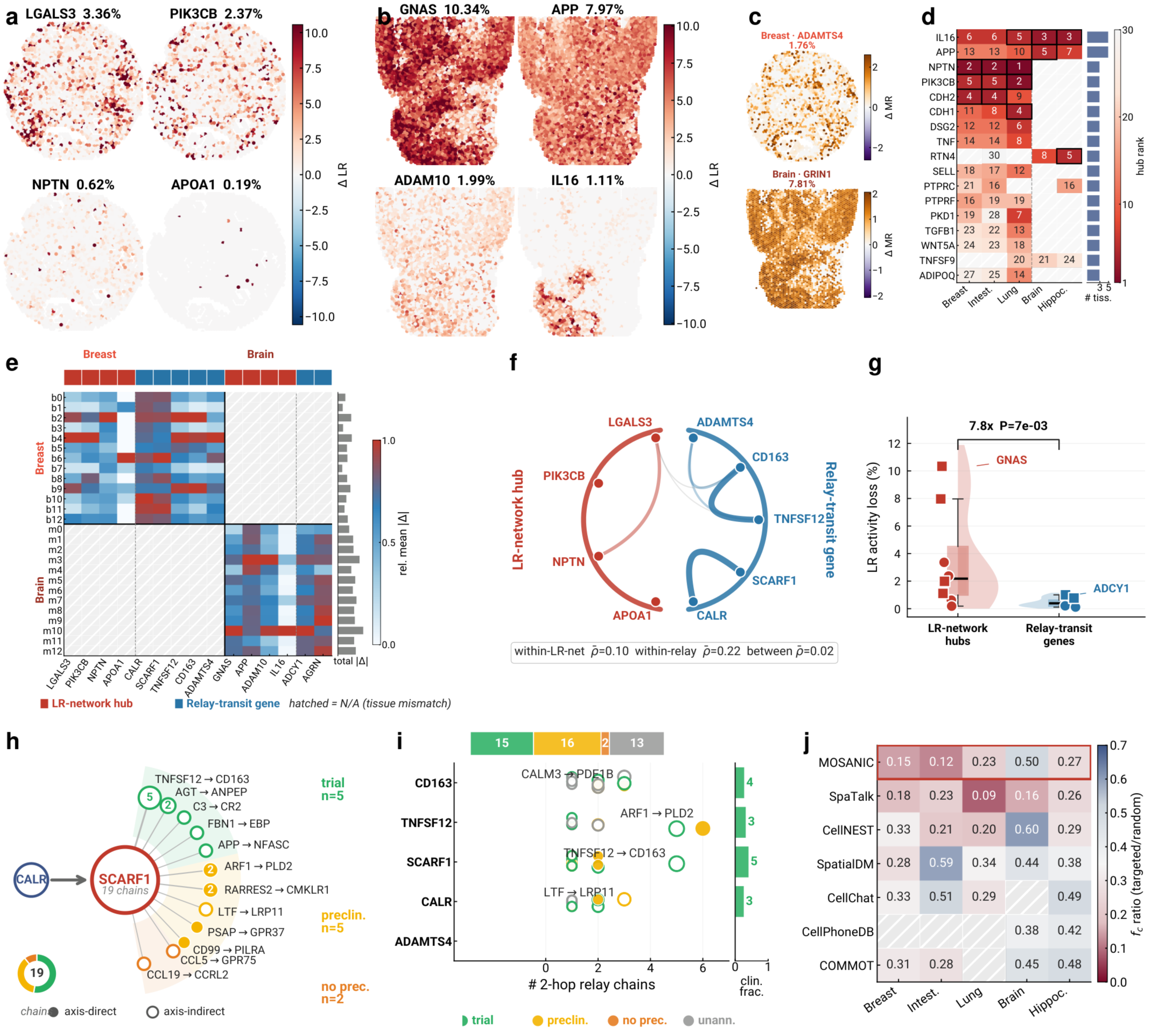
Hub vulnerabilities partition tissue-specific communication architectures. **a, b**, ΔLR maps after *in-silico* attention-edge knockout of the top-4 LR-network hubs (Methods §4, Eq. 18 with C = ε2) in breast (LGALS3, PIK3CB, NPTN, APOA1) and brain (GNAS, APP, ADAM10, IL16); fractional LR-activity loss ranges from breast 3.36 → 0.19 % and brain 10.34 → 1.11 % (Supplementary Fig. S25, Supplementary Table T11, Supplementary Note N7). **c**, Stacked ΔMR maps for the top MR hub per tissue (breast ADAMTS4 1.76 %; brain GRIN1 7.81 %; 4.4× ratio). **d**, Rank heatmap of the 17 hub genes universal across ≥ 3 of 5 tissues; right margin = universality count (Supplementary Fig. S26). **e**, Mean |ΔLR| per cluster (13 breast + 13 brain) × 15 KO genes; top strip codes the LR-network-hub class (red) versus the relay-transit-gene class (blue) (Methods §4, Eq. 18 vs Eq. 20). **f**, Chord diagram of Δ-vector Spearman ρ across 4 LR-network hubs and 5 relay-transit genes in breast; ρ: within-LR-network-hub 0.10, within-relay-transit-gene 0.22, between-class 0.02. **g**, LR-network-hub versus relay-transit-gene KO disruption (8 + 7 genes pooled across breast and brain; mean fractional LR-activity loss 3.49 % vs 0.45 %, ratio 7.8×; Mann–Whitney one-sided *P* = 7 × 10⁻³; Supplementary Fig. S29). **h**, SCARF1 → 12 LR pairs grouped by tier (trial / preclinical / no precedent); filled = axis-direct, hollow = axis-indirect (Supplementary Table T12). **i**, Top-5 breast relay-transit genes × LR-pair dot plot; size = chain count, colour = tier, fill = axis-direct. **j**, Percolation fc^tgt/fc^rnd across 5 tissues × 7 methods; MOSANIC outlined red; methods with LCC < 8 hatched as undefined (Supplementary Fig. S30). Tier evidence: Supplementary Note N8.

These hubs are not tissue artefacts, they generalise across cancers and brain. Cross-referencing the top-30 hub list across five datasets identified 17 LR genes appearing in ≥ 3 of 5 tissues (Fig. 6d). Five of them IL16, APP, NPTN, PIK3CB, CDH2 appears in top-10 for every dataset in which they appeared, and 4 of the 8 are ablation targets (Fig. 6a,b), ruling out over-fitting to one cancer or species. Stratifying per-spot ΔLR by Leiden cluster (Fig. 6e; per-cell-type breakdown for both tissues in Supplementary Fig. S29) showed that breast clusters b2, b4 and b9 absorb most cancer-side disruption across both LR-network-hub and relay-transit-gene classes, while brain cluster m10 (oligodendrocyte-rich) carries most brain-side disruption. The same row-clusters spike under multiple hubs indicating a small set of vulnerable niches carries the architectural load, recapitulating the niche-programme structure of Fig. 3.

The vulnerability landscape splits into two distinct failure modes. LR-network hubs act as global load-bearers, where ablation collapses tissue-wide LR activity, whereas relay-transit genes (top-5 of Eq. (20) in Methods, the chain-participation count introduced in Methods §4.3 and explicitly not a hub-score variant) act as routed paths whose ablation disrupts specific 2-hop chains. The two classes are mathematically orthogonal: across both tissues (8 LR-network hubs vs 7 relay-transit genes), LR-network-hub knockout reduced global LR activity 7.8× more than relay-transit-gene knockout on average (mean fractional LR loss 3.49 % vs 0.45 %; Mann–Whitney one-sided ; Fig. 6g). A 9 × 9 Spearman matrix over the 4 breast LR-network hubs and 5 breast relay-transit genes sharpens the picture (Fig. 6f): within-class = 0.10 (LR-network) and 0.22 (relay-transit); between-class = 0.02. Within the relay-transit block, an adaptive-ICD axis (CALR↔SCARF1) and an innate-macrophage axis (TNFSF12↔CD163) form distinguishable sub-blocks. The CALR↔SCARF1 axis is the recognised molecular substrate of immunogenic cell death (ICD), in which calreticulin (CALR) translocates to the dying-tumour-cell surface where the scavenger receptor SCARF1 mediates clearance by phagocytes and antigen-cross-presenting dendritic cells^60,78,79^; the TNFSF12↔CD163 axis (TWEAK↔scavenger-receptor CD163) is the canonical innate macrophage polarisation switch that drives wound-healing and pro-tumour M2 programmes^80,81^. These define two operationally orthogonal breast programmes that can be targeted independently.

Among the breast relay-transit genes, SCARF1 occupies a privileged downstream position: a single CALR input fans out through SCARF1 to 12 distinct LR pairs across 19 relay chains (Fig. 6h; per-pair evidence in Supplementary Table T12). SCARF1 has no current oncology clinical trials^60,82^, but the convergence of its CALR-driven ICD upstream and its 12-pair downstream fan-out, including IGFBP7^83^, JAG1–NOTCH^84^ and CXCL12-axis chemokine singnaling^85^ makes it as a candidate multiplexer hub. Independent confirmation comes from TCGA-BRCA (harmonised pan-cancer overall-survival table from Liu et al.^86^): SCARF1-high patients (n = 405) show significantly worse overall survival than SCARF1-low patients (n = 406; Cox proportional-hazards HR = 1.19 per log₂-TPM unit, P = 0.030; 5-year OS 72.7 % vs 81.9 %; Supplementary Fig. S32a). The association survives multivariable adjustment for the two strongest clinical confounders, tumour stage and age at diagnosis (adjusted HR = 1.17, 95 % CI 1.01–1.36, P = 0.043, n = 1,189; Supplementary Fig. S32b), establishing SCARF1 expression as an independent prognostic factor. Five of the 12 pairs are trial-precedent, five preclinical-precedent and two have no oncology precedent. Among the five principal breast relay-transit genes (Fig. 6i; n = 64 LR-pair records), 15 attain the trial-precedent phase, with SCARF1 excelling in both fan-out and clinical fraction (refer to Supplementary Table T12; tier-rule decision tree in Supplementary Note N8), results in actionable implications of the hub-score metric presented in Fig. 5. Normalised percolation across five tissues and seven CCC techniques (Fig. 6j) positions MOSANIC’s *f_c-ratio* within the lowest quartile across 4 of 5 datasets, indicating that MOSANIC’s graph disintegrates more easily under deliberate hub elimination than under random attack; whereas methodologies yielding insufficient hub nodes at the top-10% threshold (LCC < 8) are classified as undefined (Supplementary Fig. S30).

Together, these findings delineate two operationally distinct vulnerability regimes. A distributed, orthogonally partitioned cancer program and a brain communication network centralised on two dominant neuromodulatory axes (cAMP signaling via GNAS and glutamatergic/NMDA transmission via GRIN1). Furthermore, highlighting SCARF1, confirmed as an independent prognostic factor in TCGA-BRCA, as a candidate drug-actionable bottleneck in breast that warrants experimental follow-up.

## 3 Discussion

We presented MOSANIC, a heterogeneous graph transformer that reframes cell-cell communication inference as a representation-learning problem using a heterogeneous graph rather than as scoring against a fixed ligand–receptor database. MOSANIC integrates LR scoring, metabolite-receptor scoring, communication-hub identification, cross-channel relay detection and in-silico knockout within a single model class by linking cells, genes, and metabolites through seven biologically-typed edges, training on a singular expression-prediction objective, and providing various downstream applications as parameter-free read-outs of the learnt attention. Across five spatial transcriptomic datasets, three platforms, and two species, the model achieves the top AUROC on all independent LR-database evaluations against eight baseline methods, surfaces an MR channel orthogonal to the LR layer, exposes a rich-core communication backbone via the canonical hub-score, and predicts in-silico knockout phenotypes that match published literature on 23 of 24 gene-evidence axes (eight knockout genes scored on spatial pattern, cell-type specificity and disruption magnitude; Supplementary Note N9). One of the more translationally promising of these predictions being SCARF1 as a putative multiplexer hub in breast cancer that is also supported at the cohort level in TCGA-BRCA: SCARF1-high tumours show significantly worse overall survival than SCARF1-low tumours after multivariable adjustment for tumour stage and age at diagnosis (adjusted HR = 1.17, 95 % CI 1.01–1.36, P = 0.043, n = 1,189), consistent with SCARF1 expression being a stage- and age-independent prognostic factor and suggesting that the hub-score quantity may be not only architecturally interpretable but of potential clinical relevance. Three architectural properties distinguish MOSANIC. First, typed-attention factorisation isolates LR identity (ε₂), MR identity (ε₄), spatial flow (τ₁) and metabolic coupling (τ₂) into separate lanes, allowing the same Eq. (18) (see Methods) to yield the canonical hub-score or its channel-restricted variants under explicit masks. Second, read-outs are parameter-free: LR, MR, hub and knockout outputs are deterministic functions of the trained encoder’s attention and expression, never optimised against LR labels; the ablation analysis (Supplementary Note N8) confirms that expression-prediction *R*^2^ and AUROC are driven by independent model components (scVI cell features for *R*^2^; ε₂ LR-edge for AUROC), hence we can conclude the LR evaluation against OmniPath, ConnectomeDB2025 and NeuronChatDB gives us a quantifiable rankings among baselines. Third, the architecture is multi-channel by construction: MR pair scoring, cross-channel relay-chain detection and channel-restricted hub-scores arise from the same encoder pass without auxiliary models providing a flexible ground for different application.

MOSANIC offers a cross-channel, interpretable framework for CCC network building and hub scoring; nonetheless, there are several scope of improvements based on the axes of interpretation. MOSANIC factorises spatial flow and gene-pair identity rather than learning a joint cell-context-dependent LR attention. The trade-off is that the model cannot represent cell-type-specific co-option of an LR pair at the attention level, although differential intensity in a niche emerges only through expression weighting in the canonical read-out. This is a deliberate choice that preserves non-circular evaluation, but it limits the discovery of context-specific signaling that single-cell-context attention could in principle resolve. The training-time LR databases constrains which ε₂ edges exist; pairs absent from the vocabularies cannot receive attention regardless of spatial evidence, even though the per-cell expression-weighted read-out exposes downstream candidates. Scaling to whole-slide imaging platforms (Stereo-seq, Visium HD) is currently constrained by memory limitations due to the complete materialisation of the typed graph; implementing randomised sub-graph sampling, as utilised by GraphSAGE^87^ can be a logical subsequent step. Finally, although the dependency on external foundational embeddings (scVI, ESM-2, ChemBERTa) and methods (scFEA) provide us multi-modal interpretation and a powerful tool to push the limit of current state-of-the-art, come with significant computational and memory constraints. MOSANIC comes with a flexible and extendible framework. Our future work can be considered to expand MOSANIC to integrate (i) cell-context-dependent LR attention via per-cell modulation of the ε₂ edge attribute, that would close the gap to context-specific signaling without breaking the parameter-free read-outs; (ii) pre-training the typed-edge attention on public spatial atlases^88,89^ would amortise per-dataset training and enable zero-shot CCC inference; (iii) proteomics-resolved spatial assays (CODEX, Stereo-CITE-seq) would let ε₁ carry direct protein-level evidence; in-silico knockout interpretability could be sharpened by combining edge-ablation with GEARS-style perturbation prediction^90^ and by extending the hub-score concept to the regulome (SCENIC+)^91^.

MOSANIC contributes a reusable design pattern for spatial multi-omic reasoning: foundation-model embeddings on biologically-typed nodes; per-edge-type typed attention; cross-edge-type softmax gating; and parameter-free read-outs as the unit of inference. The single per-gene hub-score is intentionally portable. It can be re-used to nominate drug targets (like we performed independent cohort-level prognostic validation of the top novel candidate, SCARF1, in TCGA-BRCA), to identify niche-programme drivers, to compare communication architectures across species, or to flag cross-tissue universal nodes that warrant experimental follow-up. We distribute MOSANIC as the open-source *MOSANIC-cci* Python package with a structured pipeline from raw .h5ad to communication maps, knockout outputs, and SCARF1-style multiplexer reports, alongside the trained checkpoints and reproducibility elements discussed in this work. The same architecture can be retrained on any spatial transcriptomic dataset, any LR/MR vocabularies, and any foundation embedding without modification — turning communication-graph inference from a per-paper engineering exercise into a deployable component of the spatial biology stack.

## 4 Methods

MOSANIC is a heterogeneous graph transformer that infers cell–cell communication (CCC) from spatial transcriptomics by learning attention over biologically-typed ligand–receptor (LR) and metabolite–receptor (MR) edges. The model is summarised in Fig. 1: three node types (cell, gene, metabolite; Fig. 1c) carry pre-trained foundation-model embeddings (Fig. 1b); a heterogeneous graph wires them with three cell–cell τ edges (τ₁ secreted, τ₂ metabolite-mediated, τ₃ intracellular) and four cross-type ε edges (ε₁ cell→gene, ε₂ ligand→receptor, ε₃ cell→metabolite, ε₄ metabolite→receptor); a two-block encoder runs seven parallel typed attention layyers per block (Fig. 1d, Fig. 1e); and four read-out heads — niche clustering, LR/MR pair scoring, hub-score, *in-silico* knockout — read off the learned attention at zero added parameters (Fig. 1f). The notation introduced here (*τ*, *ε*, h, *α*, *β*, hub) is identical to Fig. 1 and is used unchanged throughout Figs. 2–6.

### 4.1 Datasets, databases and preprocessing

#### Spatial transcriptomics input

The model takes as input an *AnnData* object containing a raw count matrix for spatial units across genes, together with two-dimensional spatial coordinates . As is standard for spatial-graph methods that operate uniformly across capture technologies, we use the term “cell” throughout as a generic label for one spatial measurement unit, a 55-µm spot for Visium and Visium FFPE (each capturing approximately 1–10 cells), an approximately 10-µm near-single-cell bead for Slide-seqV2, and a segmented single cell for Xenium. Each such unit is one cell-node of MOSANIC’s graph, and all per-cell quantities (embeddings, attention, niches, hub-scores) are defined at this resolution; spot-level cell-type labels denote the dominant or deconvolved cell type of the unit. Five datasets were evaluated: human breast tumour, intestinal cancer and lung cancer (Visium / Visium FFPE / Xenium; 2,516 / 2,660 / 36,601 cells), and mouse cortex and hippocampus (Visium / Slide-seqV2; 3,342 / 41,674 cells).

We pre-processed the raw-count metrics by following a standard *scanpy* pipeline. For each dataset we computed quality-control metrics, removed cells with fewer than 200 detected genes or fewer than 500 total UMIs and cells with > 20 % mitochondrial reads, then removed genes detected in fewer than three cells along with all mitochondrial transcripts. Raw counts were library-size-normalised to 10,000 counts per cell and log1p-transformed; the 2,000 highest-variance genes were retained for foundation-model embedding. For human datasets the cell foundation-model input is a 128-dimensional *scVI* latent trained from raw counts after gene filtering with default *scvi-tools* hyperparameters^19^; for mouse datasets *scVI* was retrained per dataset on the analogous filtered count matrix. The same pre-processed AnnData object feeds (i) the graph builder (ii) the scVI cell-embedding training and (iii) the held-out expression target y*_i_* used as the training signal (Eq. 13).

#### Ligand–receptor and metabolite–receptor catalogues

Graph construction used species-matched LR and MR databases from different openly available sources including CellNEST (14,909 human LR pairs labelled contact / secreted / ECM)^14^, CellTalkDB (6,185 mouse LR pairs)^92^, and MEBOCOST (794 human / 790 mouse MR pairs)^15^. Evaluation used three independent databases with zero overlap against the training catalogues: OmniPath ligrec-extra (6,555 human LR pairs)^23^, ConnectomeDB2025 (3,550 pairs)^93^, and NeuronChatDB (452 mouse neuropeptide–receptor pairs)^24^. The zero overlap makes the Fig. 2 AUROC evaluations non-circular.

#### Metabolite flux (scFEA)

Per-cell metabolic flux was estimated with scFEA, a stoichiometry-constrained graph neural network that infers reaction-level flux from gene expression. For each cell *i*, scFEA produces a flux vector **f***_i_* ∈ ℝ*^M^* with *M* = 70 (human) or *M* = 168 (mouse) modules. Flux is consumed in three places downstream: (i) as cell→metabolite edge features on ε₃, (ii) as a per-module gate on cell–cell τ₂ edges and (iii) as the ligand-side intensity weight in the canonical MR score (Eq. 14).

#### Spatial graph

The base spatial graph connects each cell to its *k* = 6 nearest neighbours by Euclidean distance on **C**, subject to a platform-dependent maximum-distance threshold *d*_max_ of 150 µm for Visium and 50 µm for Xenium and Slide-seqV2. The same neighbour graph seeds all three cell–cell edge types (τ₁, τ₂, τ₃).

### 4.2 MOSANIC model

#### Heterogeneous graph

The graph G = (V, ℰ) has three node sets V*_c_*, V_g_, V*_m_* and seven edge types: three cell–cell τ edges and four cross-type ε edges. Cell–cell τ edges are bidirectional and seeded from the spatial *k*-NN graph (§1.4): τ₁ secreted connects neighbours within *d*_sec_ that share at least one expressed LR pair, carrying a Gaussian distance weight; τ₂ metabolite-mediated connects neighbours within *d*_met_ that share non-zero scFEA flux in at least one module; τ₃ intracellular is a per-cell self-loop with a 102-d attribute combining a 32-d PCA of receptor expression with the normalised scFEA flux vector. Cross-type ε edges are directed: ε₁ cell→gene connects each cell to its top-500 expressed genes with ligand/receptor flags; ε₂ gene→gene connects every LR pair present in the dataset vocabulary of CellNEST. LR pair scores are read off ε₂ attention; ε₃ cell→metabolite connects cells to metabolites whose scFEA flux exceeds the per-module median; ε₄ metabolite→gene connects every MEBOCOST pair, with catalogue confidence as attribute and MR pair scores are read off ε₄ attention.

#### Node features (

Each node type carries a frozen pre-trained foundation embedding chosen to capture biochemical identity beyond expression. Cell nodes carry a 128-dimensional scVI latent trained on the dataset’s full count matrix^19^; gene nodes carry the 1,280-dimensional ESM-2 650M protein embedding (esm2_t33_650M_UR50D)^20^; metabolite nodes carry the 600-dimensional ChemBERTa-77M-MTR SMILES embedding^21^. All three node types are projected to a shared latent of dimension *D* = 256 by a node-type-specific MLP (Fig. 1b):

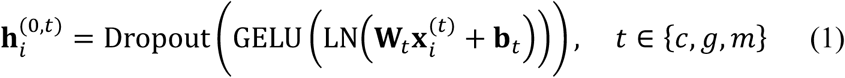

with **W***_c_* ∈ ℝ*^D^*^×128^, **W**_g_ ∈ ℝ*^D^*^×1280^, **W***_m_* ∈ ℝ*^D^*^×600^ and dropout rate 0.1.

#### Heterogeneous graph transformer encoder

The encoder stacks *L* = 2 blocks, each running **seven parallel typed attention lanes** (one per edge type; Fig. 1d) whose outputs are aggregated to each destination node-type by a learnable softmax gate and refined by a per-node-type feed-forward sublayer.

For edge type *t* and head ℎ ∈ {1, …, *H*} with *H* = 4 and *d*_ℎ_ = *D*/*H* = 64, per-edge attributes are projected to the head dimension and added to both keys and values:

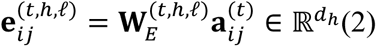

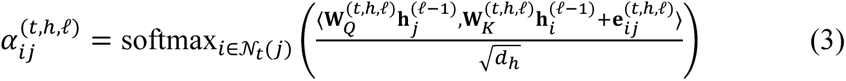

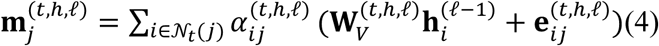

Eq. (3) normalises attention per destination node, making *α*_*ij*_^*(t,ℎ,l)*^ the relative importance of source *i* among all neighbours of destination *j* over edge type *t*, the property that makes ε₂ attention a direct LR-pair score and ε₄ attention a direct MR score (§3). Multi-head outputs are concatenated and combined with a per-lane β-gated residual:

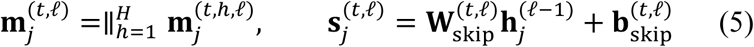

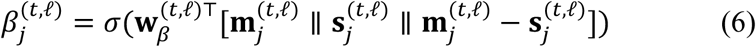

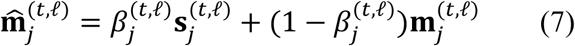

Each destination node type receives messages from multiple edge types (cell from τ₁/τ₂/τ₃; gene from ε₁/ε₂; metabolite from ε₃/ε₄). The lane outputs are aggregated by a learnable softmax gate per (destination type, layer):

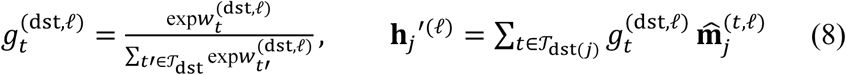

A residual from the block input is added and normalised:

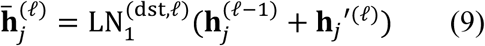

Each block ends with a per-node-type two-layer FFN (GELU, 4× expansion, dropout 0.1) with post-norm residual:

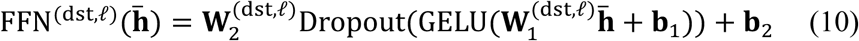

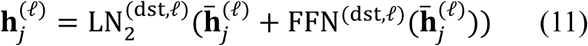

After *L* = 2 blocks the encoder outputs refined cell embeddings {**h**_*i*_^*(L)*^ ∈ ℝ^256^} and retains all attention tensors 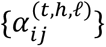 across edges, heads and layers for the four parameter-free read-out heads. Three independent residuals per block (per-lane β-gate, Eq. 7; post-aggregation Add+LN, Eq. 9; post-FFN Add+LN, Eq. 11) stabilise training; no residual crosses the two stacked blocks.

#### Expression decoder

The single trained loss is held-out expression prediction. The decoder maps the final cell embedding to a predicted log-normalised vector across *K*_tgt_ = 200 target genes (highest-variance subset):

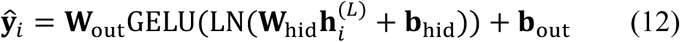

with decoder dropout 0.2; target **y***_i_* is the log-normalised expression of the 200 highest-variance genes. The loss is Huber (*δ* = 1):

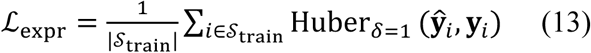

To prevent spatial-autocorrelation leakage, cells are partitioned by *k*-means on **C** (*k* = max(10,min(30, 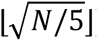))) and clusters assigned 70/15/15 % to train/validation/test, so neighbouring cells share a split and the test-set *R*^2^ conservatively estimates generalisation to unseen spatial regions. We optimise with AdamW (LR 10^−3^, weight-decay 10^−4^), clip gradients at norm 5.0, reduce LR by 0.5× on validation-*R*^2^ plateau (patience 15), and stop after 50 epochs without improvement (max 500). Node features and prediction targets are different representations of the same expression matrix i.e. scVI compresses the full transcriptome while targets are 200-gene measurements and the spatial graph is constructed from coordinates alone; the model therefore must learn spatial context to improve predictions, which forces meaningful attention over ε₂ LR edges (ablation: removing ε₂ collapses OmniPath AUROC to 0.500; Supplementary Note N8).

### 4.3 Communication-score extraction

A single forward pass of the trained model with return_attention=True exposes the following read-out heads.

#### Intensity-weighted LR score

For LR pair (*l*, *r*), the raw attention score is the mean ε₂ coefficient on edge (*l* → *r*) across heads and layers:

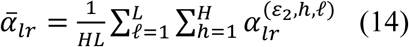

Because Eq. (3) normalises per destination, *ā_lr_* is the relative importance of ligand *l* among all ligands binding receptor *r*. This is meaningful within receptor but introduces a known degeneracy: receptors with in-degree 1 always receive *ā* = 1. To break the degree-1 saturation and produce a unique paper-wide ranking, the canonical MOSANIC LR score is the intensity-weighted form, used everywhere unless otherwise noted:

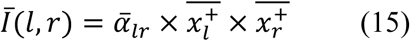

where 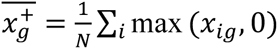 is the mean clipped expression of gene *g* across all *N* cells. Eq.

1. resolves all 183 degree-1 ties in breast (final tie count 0/9,571). The **per-spot map** used for spatial visualisations (Figs. 3, 6) is the un-aggregated form:

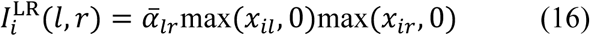

Six degree-corrected variants of *ā_lr_* (linear-degree, log-degree, expression, combined, etc.) are exposed through the API and tabulated in Supplementary Table **??**; the best-per-(dataset × database) is reported in Fig. 2.

#### Intensity-weighted MR score

For MR pair (*k*, *r*) (metabolite *k*, receptor gene *r*), the analogue substitutes scFEA flux on the ligand side:

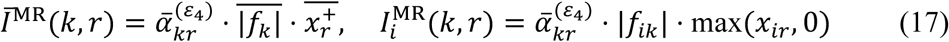

with 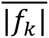 the mean absolute flux of metabolite *k*. As with LR, intensity weighting breaks ties at degree-1 receptors and produces unique pair ranking.

### 4.4 Hub-score: one formula, three channel restrictions, one separate chain-count

For a gene node *g* and a channel mask C ⊆ {*ε*_2_, *ε*_4_}, we define the hub-score as

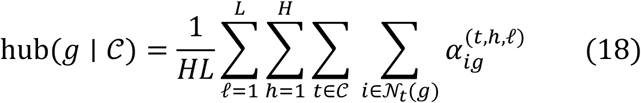

where 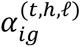 is the attention weight on the type-*t* edge *i* → *g* at head ℎ of encoder layer *l*, N*_t_*(*g*) is the set of type-*t* neighbours of *g*, and *H* and *L* are the numbers of attention heads and encoder layers. Equation (18) sums all incoming attention into *g* over the selected channels and averages over the *HL* head-layer pairs; it is the single hub-score calculation used in the paper. Every gene-level hub quantity is an evaluation of Eq. (18) under a fixed mask, and the three named variants differ only in C:

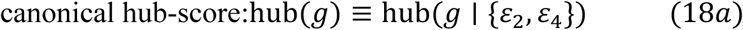

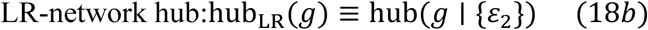

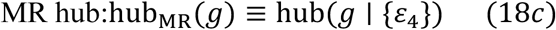

The canonical hub-score, Eq. (18a), evaluates Eq. (18) over both channels and is used wherever the paper writes “hub-score” without qualification; hub genes are its top 10 % per dataset (the universal-hub list of Fig. 3, the LCC and rich-core of Fig. 5, the cross-tissue universality of Fig. 6d, and the cell-type × hub heatmap of Fig. 6e). The two single-channel variants name which channel was ablated in the corresponding knockout protocol (§5.5): the LR-network hub, Eq. (18b), gives the top-4 per tissue used for LR-channel knockout (Fig. 6a, b, and the NET class of Fig. 6g), and the MR hub, Eq. (18c), gives the top-20 per dataset (Fig. 4c, d) and the top-1 per tissue (Fig. 6c) used for MR-channel knockout.

#### Per-cell hub-score (Fig. 5a inset)

The same construction applied to a *cell* node, restricted to τ₁ secreted out-attention:

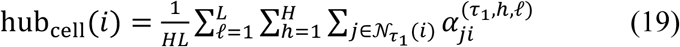

Hub cell (Fig. 5a inset) = cell in the top 5 % of Eq. (19). Cell hub-score is conceptually parallel to Eq. (18) but defined on a different node type (cells), so it uses a different attention channel (τ₁, the only attention channel incident on cells).

#### Relay-transit gene: chain-participation count (Fig. 6h, i) — *not* a hub-score variant

To avoid confusion with the attention-derived hub-score, the chain-participation quantity used in Fig. 6 is called the relay-transit gene count and is denoted transit(*g*). It is defined operationally on the validated 2-hop relay subgraph ℛ^(2)^ (§5.2):

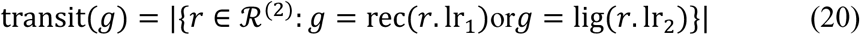

Eq. (20) counts how many 2-hop chains transit through *g* at the connection point between the two LR steps. It is not an attention-weighted degree and not a special case of Eq. (18); it is orthogonal to it (Fig. 6f, g: within-LR-network-hub ρ: = 0.10, within-relay-transit-gene ρ: = 0.22, between-class ρ: = 0.02; LR-network-hub KO disrupts global LR activity 7.8× more than relay-transit-gene KO, Mann–Whitney *P* = 7 × 10^−3^). The “relay-transit genes” in Figs. 6h–i are the top-5 by Eq. (20) per tissue (CALR, SCARF1, TNFSF12, CD163 and ADAMTS4 in breast; ADCY1 and AGRN in brain).

### 4.5 Downstream analyses Niche clustering

Cell embeddings {**h**^*(L)*^} are passed through Leiden clustering (resolution *r* = 1.0, scanpy v1.10) to produce communication niches. Niche purity against source-annotated cell types is measured by the maximum-class-fraction metric; spatial coherence is the fraction of *k* = 15 spatial neighbours that share a niche label (Fig. 3a, 3b).

#### Relay-chain detection

A relay chain is a path *A* → *B* → *C* where (i) *A* signals to relay-hub cell *B* via LR pair (*L*_1_, *R*_1_), (ii) *B* signals to *C* via a *distinct* LR or MR pair (*L*_2_, *R*_2_), with both edges retained at the top-10 % attention threshold, (iii) *B* is a shared spatial neighbour of *A* and *C* within 150 µm, and (iv) the relay score *τ* = *α*(*A* → *B*) ⋅ *α*(*B* → *C*) exceeds the permutation null at *P* < 0.05. A cross-channel relay further requires hop-1 on τ₁ and hop-2 on τ₂ + ε₄. The full detection algorithm and threshold-sensitivity analysis are in Supplementary Note N5.

#### Rich-core and bootstrap stability

The weighted rich-core coefficient *ρ*_w_(*k*) is the ratio of observed top-*k*-clique density to a degree-preserving Maslov–Sneppen rewire null (100 randomisations). Bootstrap-stable hubs are identified by 1,000 edge-resamples of the top-10 % graph, retaining genes in the top-20 of ≥ 90 % of resamples.

#### Network percolation

Normalised percolation *r* = *f_c_*_,targeted_/*f_c_*_,random_measures targeted-attack fragility relative to random attack, where *f_c_* is the fraction of nodes removed at which the largest connected component falls below half of its initial size. Targeted attack removes nodes in descending order of weighted degree (the hub-score equivalent on the unweighted top-10 % graph); the random null is averaged over 100 independent orderings. A configuration-model *z*-score compares the observed *f_c_*_,targeted_ against 100 degree-preserving rewires (Maslov–Sneppen). Methods with LCC < 8 nodes at the top-10 % threshold are hatched in Fig. 6j to indicate that their percolation ratio is undefined (Supplementary Fig. S27).

#### In-silico knockout

**For** ε₂ knockout (LR-channel ablation) of any gene *g*, we identify K_g_ = {(*l*, *r*) ∈ *ε*_2_: *g* ∈ {*l*, *r*}}, zero the attention contribution of those edges and recompute the per-spot ΔLR map:

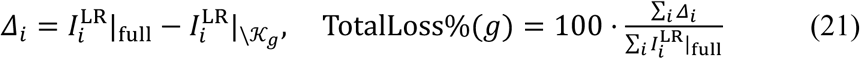

Similarly, ε₄ knockout (MR-channel ablation) was also done. Applied to GRIN1 in brain (Fig. 6c) and to the alternative MR-receptor knockouts in Supplementary Figs. S28 (breast: CD36, SCARB1, LDLR) and S31 (brain: NMDA family beyond GRIN1).

### 4.6 Evaluation metrics

#### Expression prediction (held-out *R*^2^)

Per-gene *R*^2^ between predicted (Eq. 12) and observed log-normalised expression on held-out test cells, averaged across the 200 target genes.

#### LR-pair ranking (AUROC against independent databases)

Each method emits a single score per ligand–receptor pair in its scored vocabulary. Taking membership in an independent evaluation database (§1.2 — OmniPath ligrecextra for the human datasets, NeuronChatDB for the mouse datasets; both disjoint from MOSANIC’s CellNEST training catalogue) as the positive label and all other scored pairs as negatives, pairs are ranked by score and we compute:

- AUROC — area under the receiver-operating-characteristic curve, i.e. the probability that a randomly chosen database pair outranks a randomly chosen non-database pair (0.5 = chance).
- AUPR-lift — area under the precision–recall curve divided by the random-classifier baseline (the positive prevalence in that vocabulary); 1.0 denotes no enrichment. Normalising by prevalence makes the metric comparable across methods whose vocabularies admit different positive rates.
- Lift@100 — precision among the 100 top-ranked pairs divided by the same prevalence baseline (enrichment of true pairs among a method’s highest-confidence predictions; Fig. 2b).

For MOSANIC the canonical intensity-weighted LR score (§3.2) is ranked; raw attention is used only in the saturation diagnostic of Supplementary Note N8. The full 13-variant scoring evaluation is reported there.

#### NicheNet ligand-activity validation (LigAct AUROC)

For each ligand, MOSANIC’s ligand activity is taken as *A*(*L*) = max*_R_*score(*L*, *R*) over its receptors and ranked against the NicheNet ligand-activity ground truth (104 GEO ligand-perturbation experiments; differentially expressed response genes at *q* < 0.05, |lfc| > 0.5, top-500 targets per ligand). LigAct AUROC measures how well each method’s ligand ranking recovers the experimentally active ligands against a fixed reference ligand set.

## Data availability

All spatial transcriptomics datasets analysed in this study are publicly available. The three human cancer datasets were obtained from 10x Genomics (https://www.10xgenomics.com/datasets): human breast cancer (Visium, Block A Section 1; 2,516 spots), human intestine cancer (Visium FFPE; 2,660 spots) and human lung cancer (Xenium, 5,001-gene panel; 87,461 cells). The two mouse datasets are adult mouse brain (Visium, coronal section; 10x Genomics; 3,342 spots) and mouse hippocampus (Slide-seqV2^94^; Single Cell Portal accession SCP815, 41,674 beads). Model training used species-matched interaction catalogues: ligand-receptor pairs from CellNEST (14,909 human pairs)^14^ and a CellTalkDB mouse set (6,185 pairs)^92^, and metabolite-receptor pairs from MEBOCOST (794 human and 790 mouse pairs; https://github.com/kaifuchenlab/MEBOCOST)^15^; metabolic flux was estimated with scFEA (https://github.com/changwn/scFEA)^16^. Evaluation used three ligand-receptor databases with zero overlap with the training catalogues: OmniPath ligrec-extra (6,658 human pairs)^23^; https://omnipathdb.org), ConnectomeDB2025 (3,550 human pairs; https://connectomedb.org)^93^ and NeuronChatDB (452 mouse neuropeptide-receptor pairs derived from 183 curated interactions; Zhao et al., Nat. Commun. 14, 1128 (2023)), together with the NicheNet ligand-activity benchmark (104 GEO ligand-perturbation experiments spanning 44 ligands; Zenodo record 3260758, https://zenodo.org/record/3260758)^8^. Intracellular relay validation used a STRING-derived signaling protein-protein-interaction network and a combined TRRUST/DoRothEA transcription-factor-target network were directly obtained from CellNEST repository (https://github.com/schwartzlab-methods/CellNEST)^14,49–51^. Source data for all main and supplementary figures, the pre-processed datasets, and the pre-trained per-dataset model weights are provided with this paper and will be deposited in the MOSANIC zenodo repository.

## Code availability

MOSANIC is available as an open-source Python package (mosanic-ccc) at https://github.com/debraj-55555/MOSANIC under MIT license.

