## Supplementary Information for "Learning the spatial cell-cell communication network to decode multi-channel signaling and predict network-hub vulnerabilities with MOSANIC"

*This file contains Supplementary Figures S1–S32, Supplementary Tables T1–T14, and Supplementary Notes N1–N10.*

**Supplementary Figures**

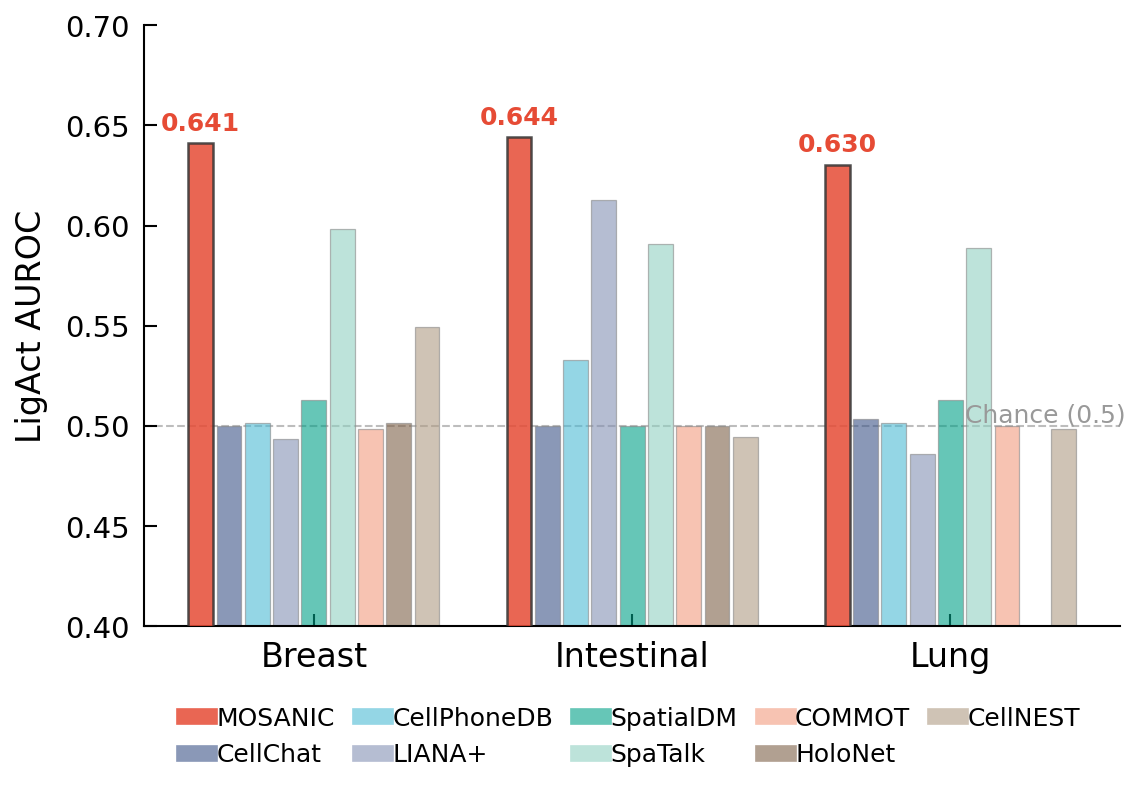

**Supplementary Fig. S1 | Functional validation against ligand-perturbation experiments..** LigAct AUROC for nine methods on three human datasets, computed against the NicheNet ligand–perturbation library (Browaeys et al. 2020; 104 GEO ligand-addition / ligand-blockade experiments). For each dataset and method, NicheNet assigns a "regulatory potential" score to each ligand–target gene pair based on observed downstream expression changes; the LigAct AUROC asks how well the method's per-LR scores rank these verified regulatory pairs above random. MOSANIC achieves LigAct AUROC 0.630–0.644 across Breast, Intestinal, and Lung, the only method consistently above chance on all three datasets.

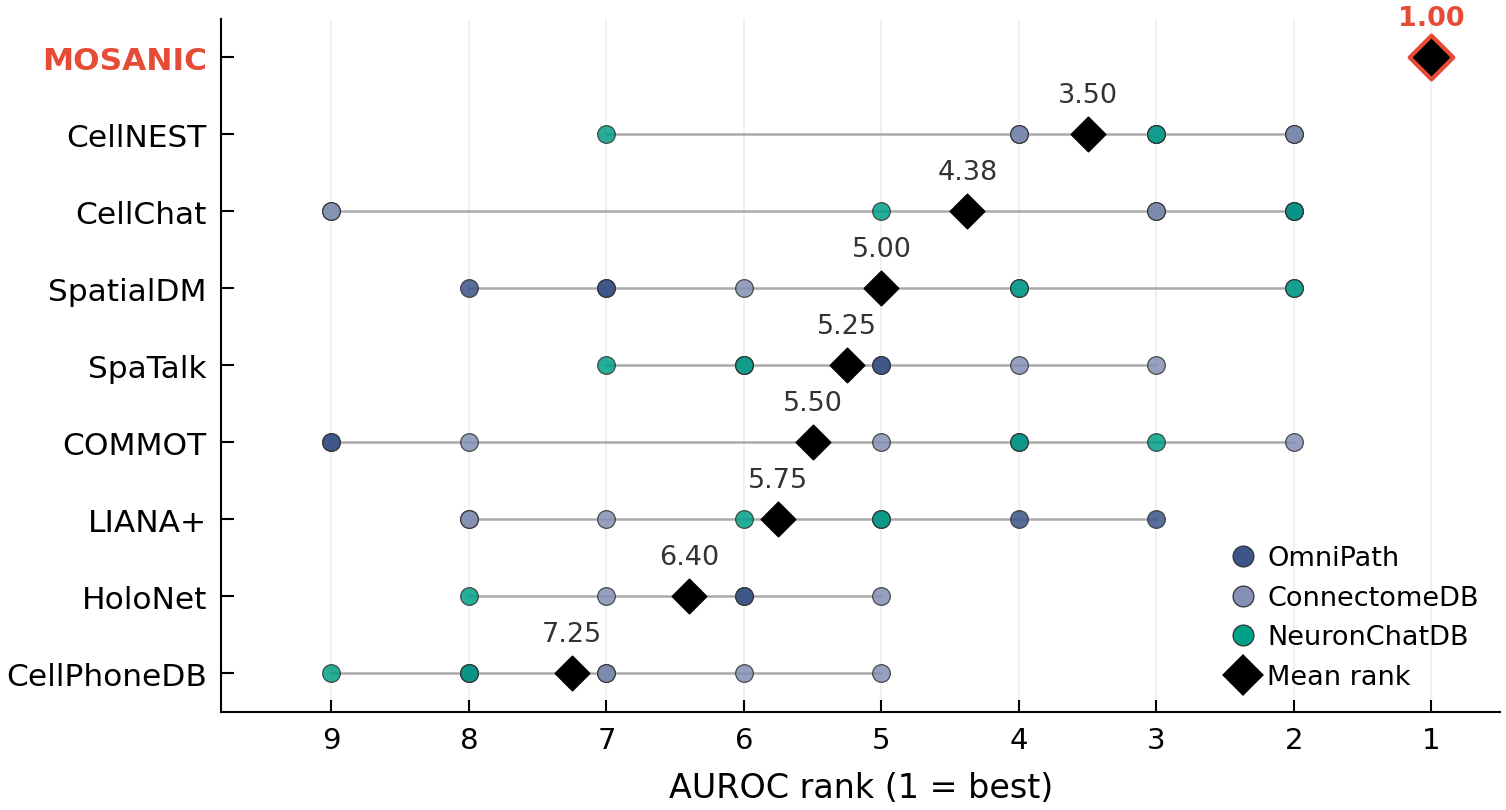

**Supplementary Fig. S2 | Mean AUROC rank across all eight held-out evaluations..** For each of the nine methods, the mean rank (1 = best) across the eight database × dataset AUROC evaluations of Fig. 2a. Individual ranks coloured by database (OmniPath blue, ConnectomeDB mid-blue, NeuronChatDB teal). MOSANIC achieves rank 1 in all 8 evaluations (mean rank = 1.00); the next-best method (CellNEST) mean rank is 3.5.

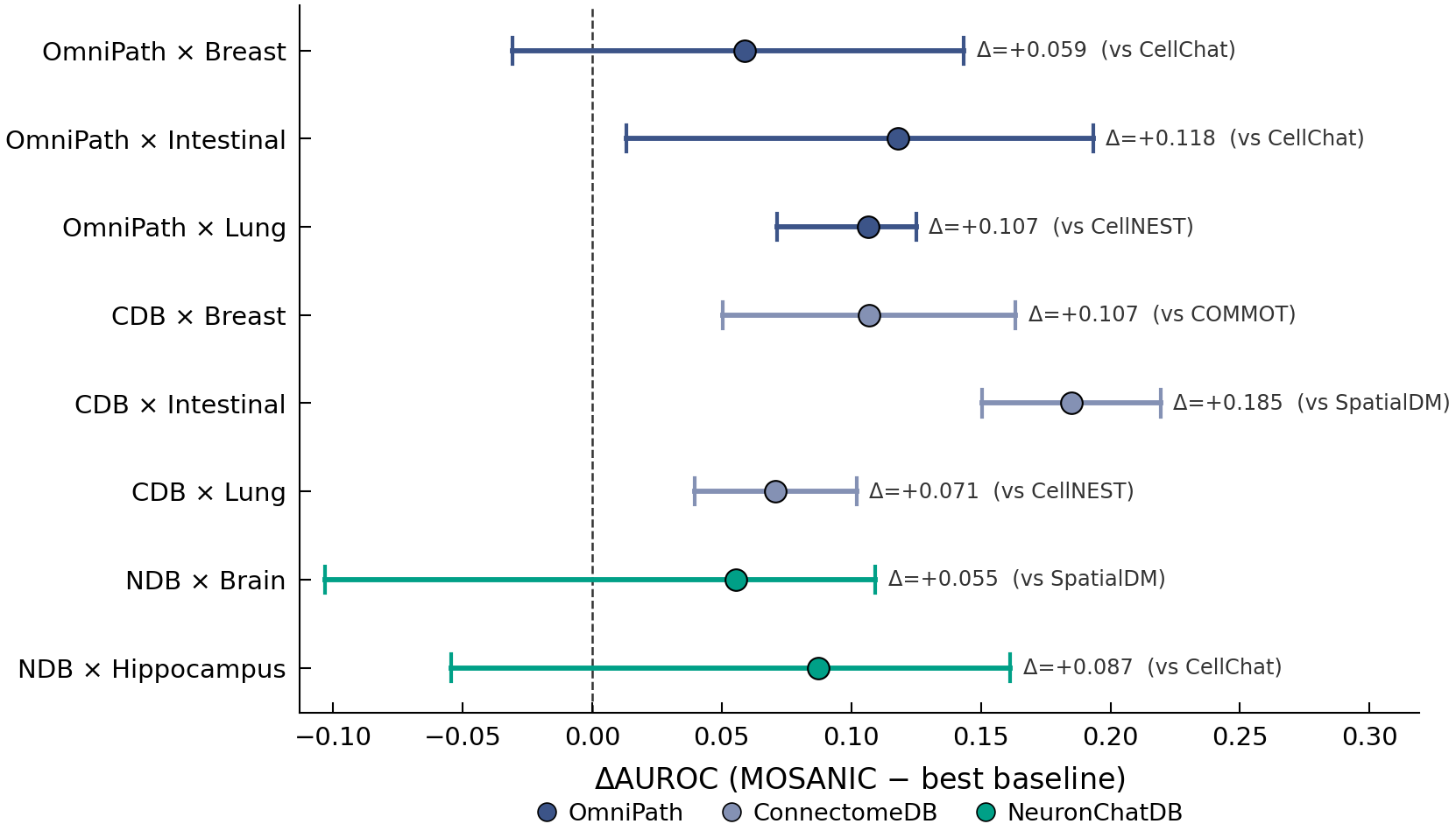

**Supplementary Fig. S3 | AUROC advantage forest plot (MOSANIC minus next-best)..** Per-evaluation AUROC difference between MOSANIC and the second-best method for that evaluation, with bootstrap 95 % confidence intervals (2,000 resamples per pair). All eight differences are positive (range +0.05 to +0.14, mean +0.099; Cohen's d = 2.35). The Wilcoxon signed-rank test comparing MOSANIC to the next-best at each evaluation yields P = 0.004 (n = 8).

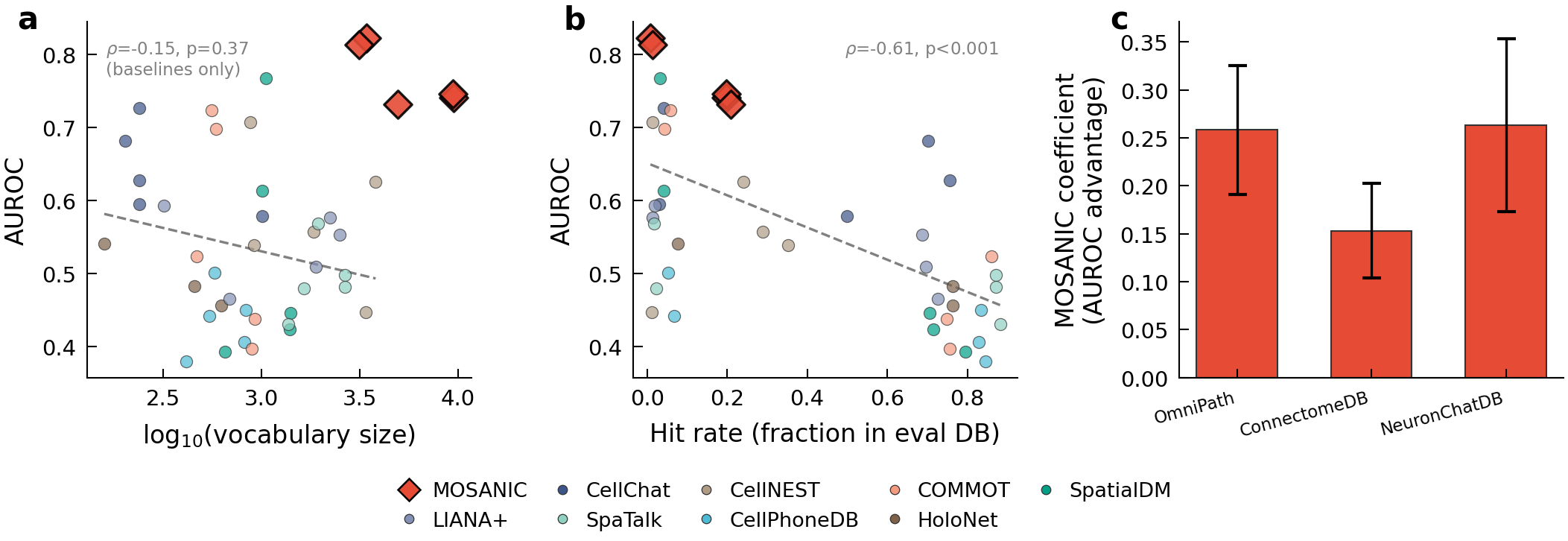

**Supplementary Fig. S4 | Vocabulary-size bias analysis..** Three sub-panels address reviewer concerns about methods scoring different vocabulary sizes: (a) scatter of vocabulary size vs AUROC across all 72 (method × evaluation) points, no systematic advantage for smaller vocabularies. (b) scatter of per-method hit rate (fraction of database pairs in method's vocabulary) vs AUROC, MOSANIC's advantage is not explained by hit-rate variation. (c) AUPR-lift (vocabulary-corrected) gain of MOSANIC over each baseline at matched top-k cutoffs, showing the gap is invariant to cutoff.

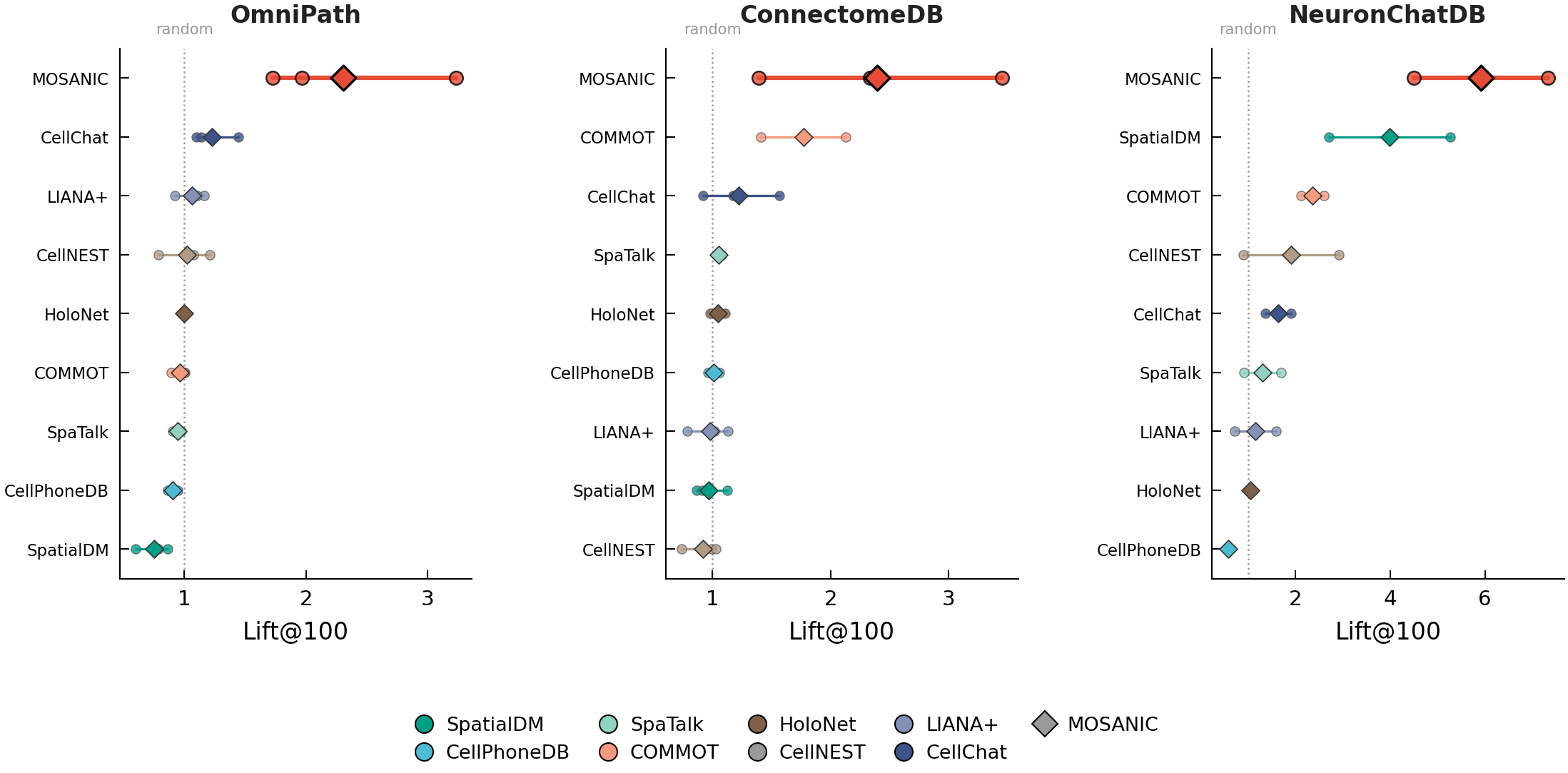

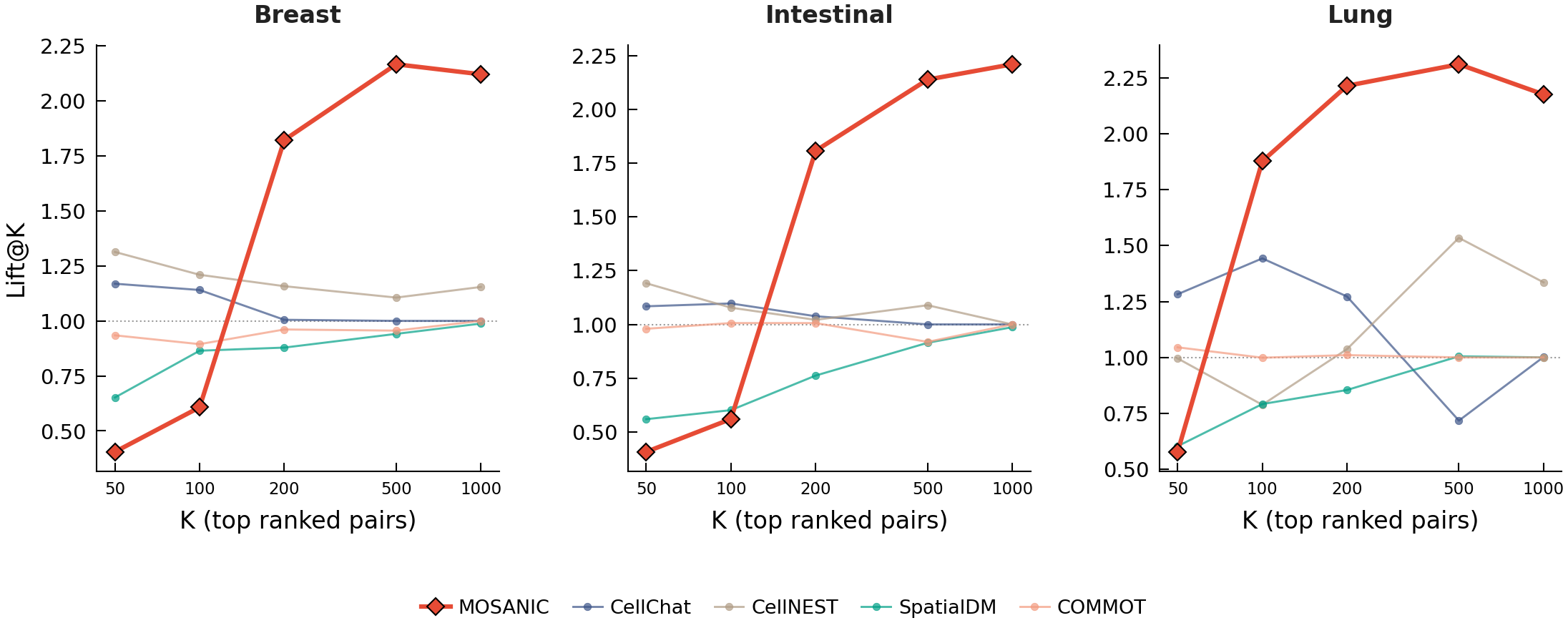

**Supplementary Fig. S5 | Lift@K enrichment curves..** For every method × evaluation, the fraction of true database pairs captured in the method's top-k ranked pairs as k increases from 10 to 1,000, normalised by random expectation ("lift"). MOSANIC's curves lie above every baseline on every evaluation; at k = 100, MOSANIC's enrichment ranges from 1.4× to 7.3× across the eight evaluations (median ≈ 2.3×), exceeding every baseline in each. Panel (a) shows Lift@100 as a dot-whisker summary across evaluations; panel (b) shows the full Lift@K curves.

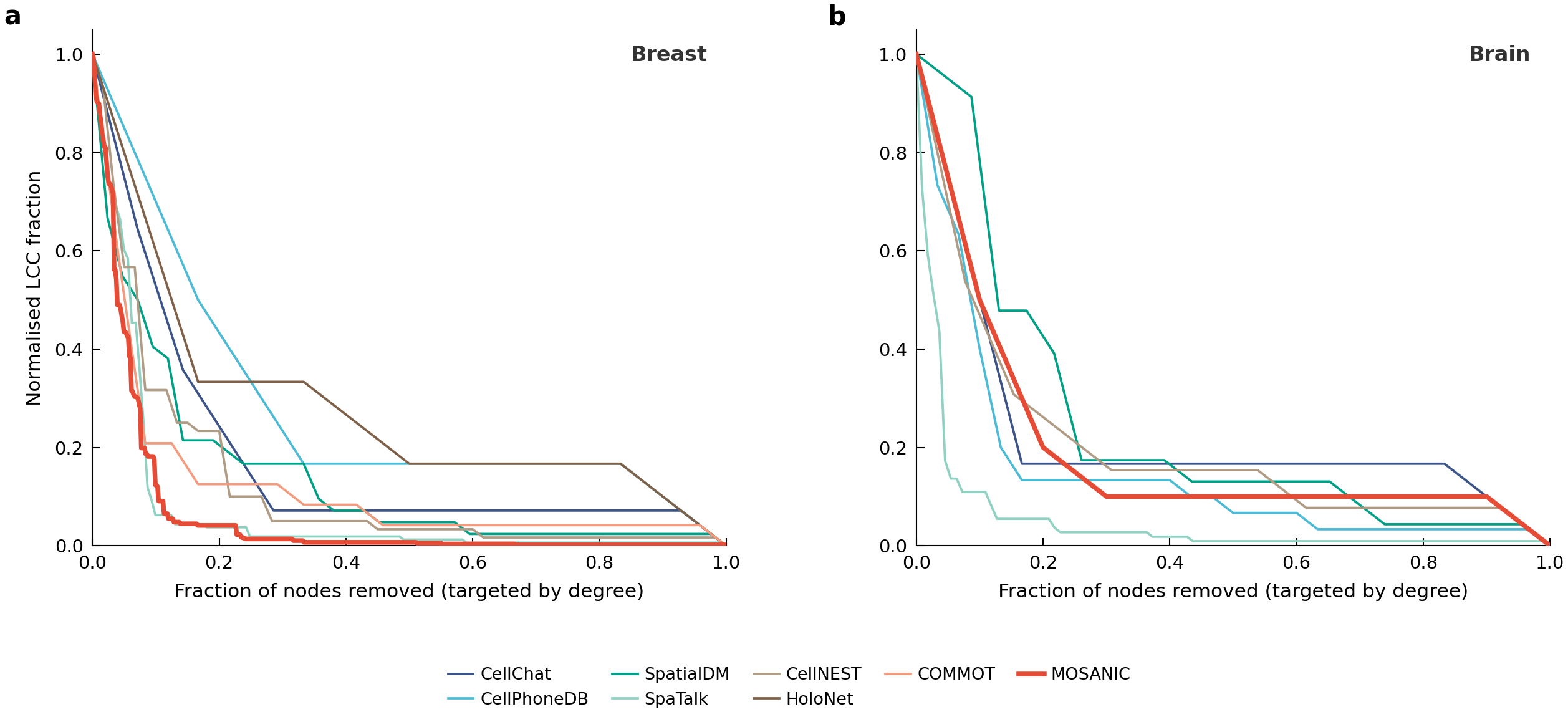

**Supplementary Fig. S6 | Cross-method percolation, architectural vulnerability across all eight baselines..** Targeted-attack percolation curves (fraction of nodes removed, in descending hub-score order, vs largest-connected-component fraction) on the breast gene graph at matched top-10 % stringency for all nine methods. To control for graph-size differences (MOSANIC's graph has 778 gene nodes; CellChat's has 201), we report the normalised ratio r = f_c_targeted / f_c_random rather than raw f_c. MOSANIC r = 0.146 (lower than all eight baselines, which span 0.181–0.332) with z-score = –15.1 against a degree-preserving configuration-model null. Brain percolation is under-powered at top-10 % (n = 10 LCC nodes across most methods; shown for completeness).

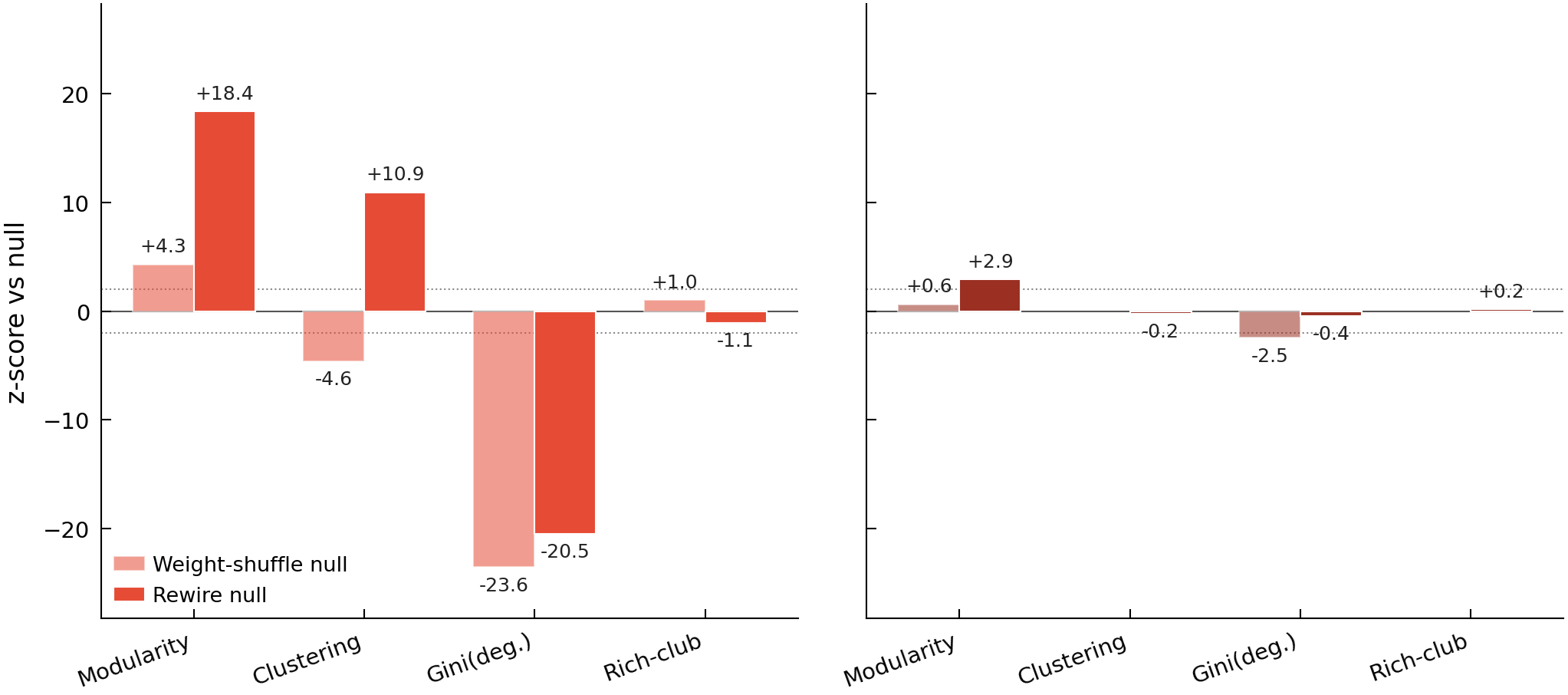

**Supplementary Fig. S7 | Maslov–Sneppen z-scores for both breast and brain graphs (extension of Fig. 2f)..** Same four topology metrics shown in the main panel f (Modularity, weighted Clustering, Gini of weighted degree, Rich-club coefficient), compared against both weight-shuffle and rewire nulls for both Breast and Brain. Weighted clustering is undefined for brain (tree-like graph, zero triangles); rich-club z-scores do not reach significance in either dataset, confirming that while hubs exist, they do not preferentially connect to each other beyond what degree heterogeneity predicts.

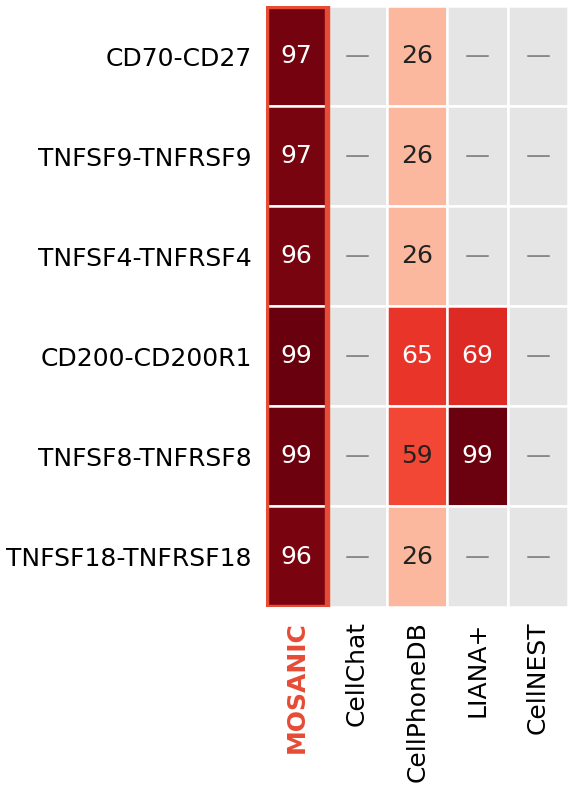

**Supplementary Fig. S8 | Vocabulary coverage of FDA-target immune-checkpoint LR pairs..** Six clinically-validated immune-checkpoint ligand-receptor pairs (CD70→CD27, TNFSF9→TNFRSF9, TNFSF4→TNFRSF4, CD200→CD200R1, TNFSF8→TNFRSF8, TNFSF18→TNFRSF18) are scored on the breast dataset by five spatial CCC methods (rows = pairs; columns = MOSANIC, CellChat, CellPhoneDB, LIANA+, CellNEST). Each cell shows the percentile rank that the method assigns the pair within its own scored vocabulary (100 = top-scored); missing pairs are absent from that method's curated vocabulary and shown as a dash. For MOSANIC, pairs are ranked by the canonical intensity-weighted LR score (Ī = ᾱ · x̄_ligand · x̄_receptor; Supplementary Note N9) rather than raw attention, which otherwise saturates these in-degree-1 receptor pairs to an identical value. MOSANIC ranks all six pairs in the top 4% of its vocabulary (96th–99th percentile); CellChat and CellNEST score none of the six (curated databases exclude them); CellPhoneDB scores all six but ranks most near the 26th percentile; LIANA+ covers only two. Because the comparison concerns which LR pairs each method's vocabulary admits, a property complementary to AUROC and AUPR-lift (Fig. 2a, b), this analysis lives in the Fig. 2 supplementary set rather than under emergent biology in Fig. 3. ---

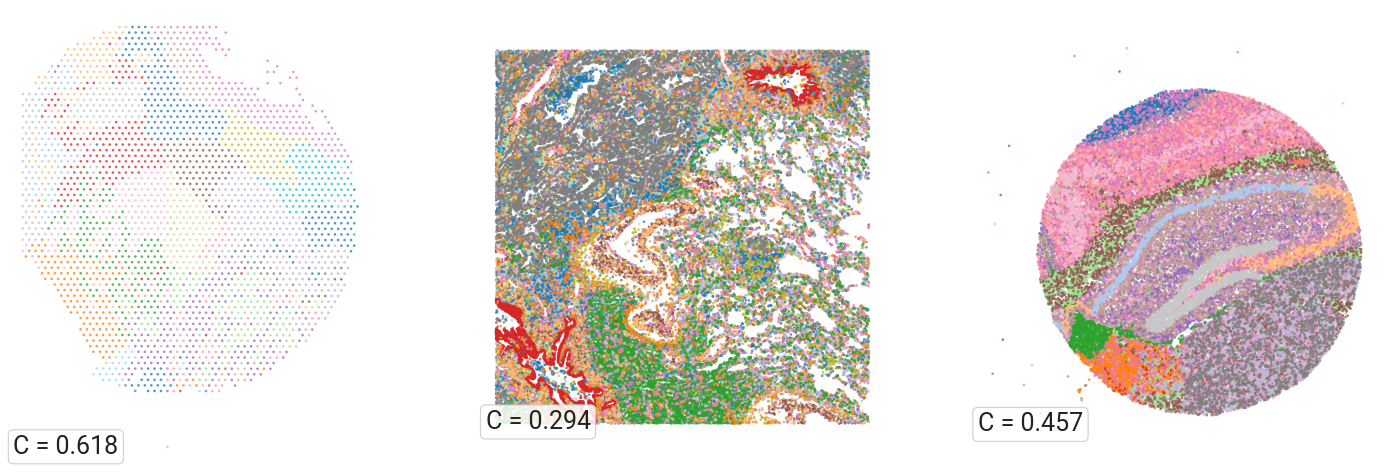

**Supplementary Fig. S9 | Per-dataset niche spatial maps (Intestinal, Lung, Hippocampus)..** Three-panel spatial scatter showing MOSANIC Leiden niches on the learned cell embedding for the three datasets not shown in Fig. 3a, b. Each scatter coloured by niche id (tab20 palette, wrapped for k > 20); inline box shows spatial coherence C (fraction of k = 15 nearest spatial neighbours sharing the niche label). Per-dataset niches: Intestinal k = 21, C = 0.618; Lung k = 25, C = 0.294 (lower C reflects the 87 k single-cell Xenium density); Hippocampus k = 16, C = 0.457. The emergent-niche phenomenon documented in Fig. 3a, b therefore generalises across all five spatial transcriptomics platforms tested.

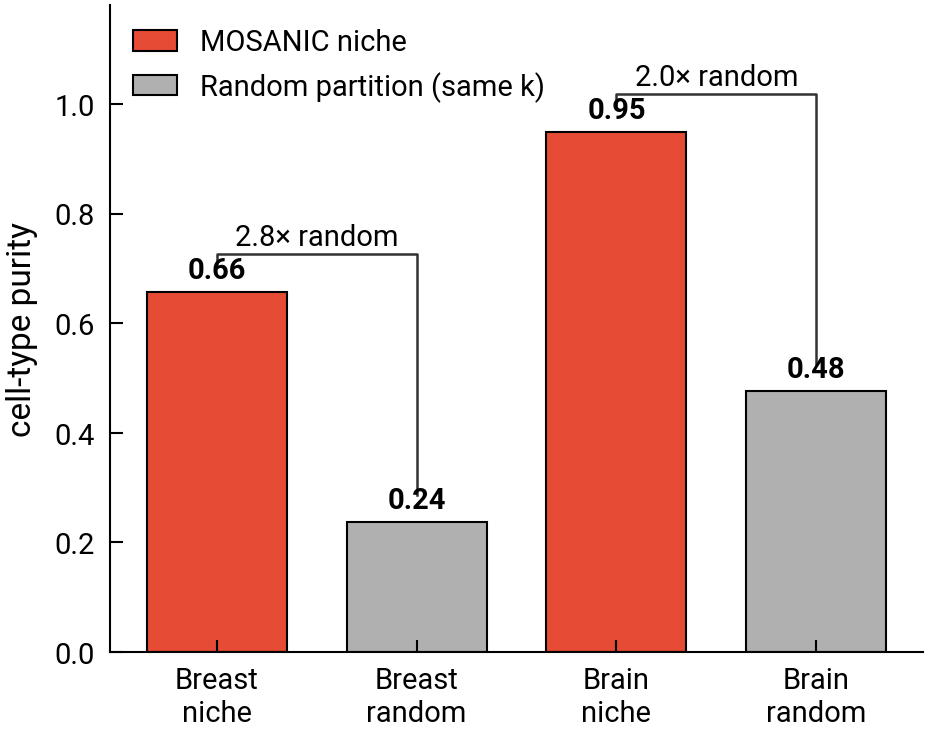

**Supplementary Fig. S10 | Niche-coherence comparison (Breast and Brain)..** Four-bar chart comparing the cell-type purity of MOSANIC's niches against a random partition of the same granularity, for Breast and Brain separately. Bar height = cluster purity (mean over niches of the largest single-cell-type fraction). MOSANIC niche bars red; random-partition baseline grey (mean purity over 1,000 label permutations of the same k). MOSANIC niches are 2.8× (breast: 0.66 vs 0.24) and 2.0× (brain: 0.95 vs 0.48) more cell-type-pure than chance, confirming that the unsupervised embedding recovers cell-type compartments without any cell-type labels during training.

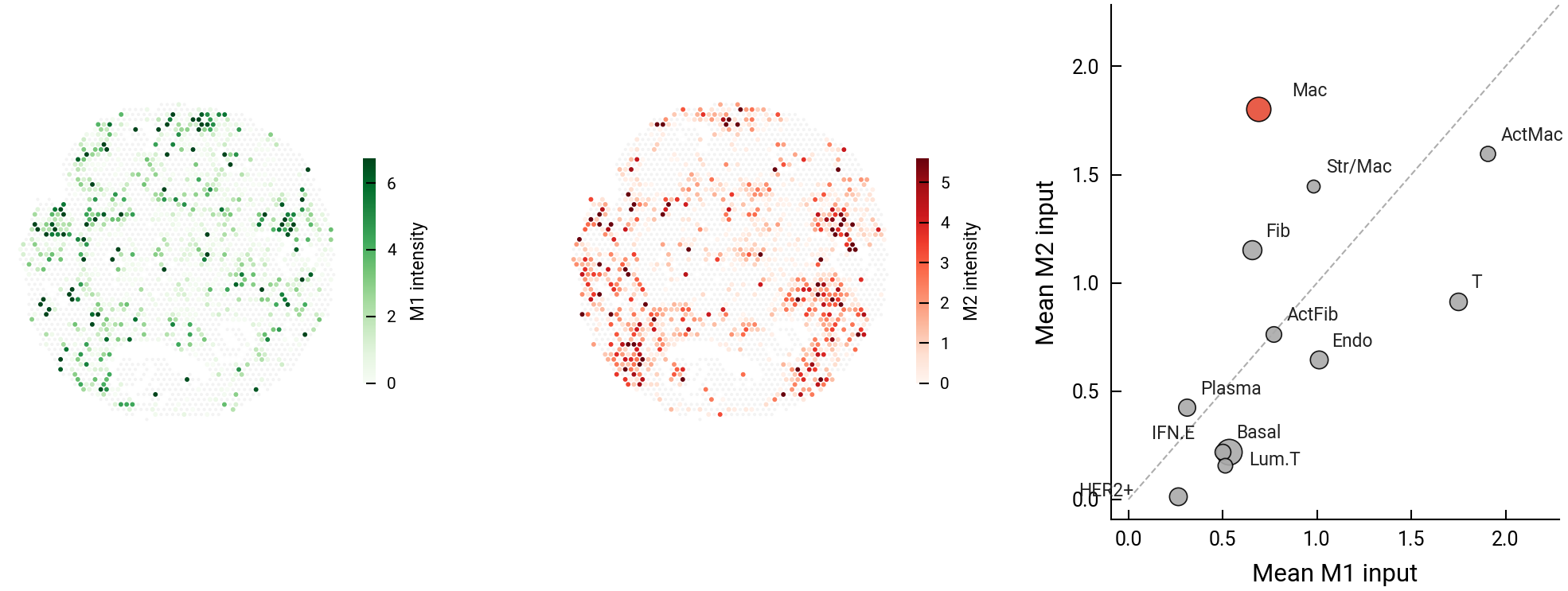

**Supplementary Fig. S11 | Extended M1 / M2 macrophage programme analysis..** Three-panel extension of Fig. 3d. Left: per-cell M1 intensity (sum of six canonical M1 pairs: IFNG→IFNGR1/2, SEMA4D→CD72, HMGB1→TLR2, CD86→CD80, SECTM1→CD7) spatially mapped on 2,516 breast spots; green gradient. Centre: per-cell M2 intensity (sum of four canonical M2 pairs: TNFSF12→CD163, LGALS3→FCGR2B, CD200→CD200R1, LEP→CD33) on the same spots; red gradient. Right: all 12 breast cell types on M1 vs M2 axes (dot size proportional to number of cells) with Macrophage highlighted; per-cell-type Wilcoxon signed-rank tests yield Macrophage P = 2.1 × 10⁻⁹ (n = 201, ratio 2.62); T cells P = 3.6 × 10⁻⁴ (n = 210); HER2+ P = 3.6 × 10⁻¹³ (n = 221).

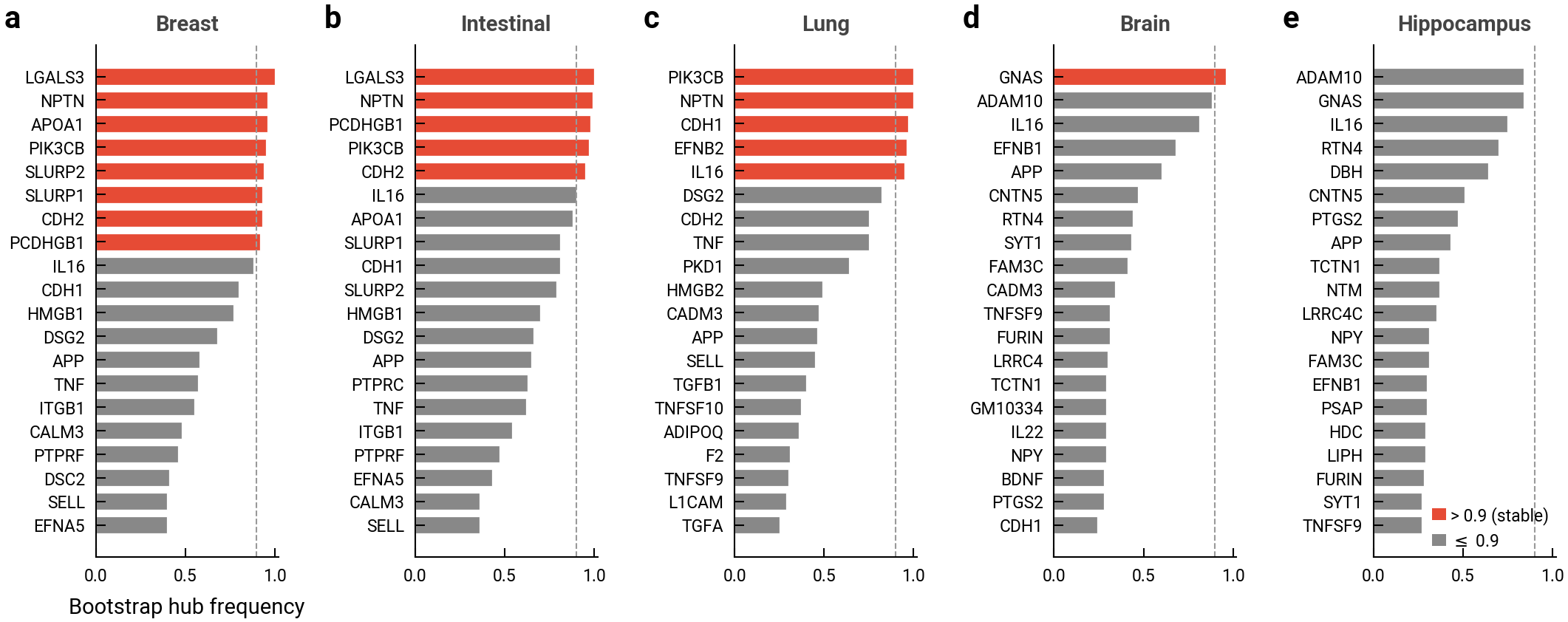

**Supplementary Fig. S12 | Bootstrap stability of hub membership per dataset..** Five horizontal bar charts (one per dataset) showing the top-20 genes ranked by bootstrap hub-membership frequency from A25. Each gene's bar is the fraction of B = 100 bootstrap resamples (drawn with replacement from the scored LR distribution) in which the gene enters the dataset's top-10 % hub list. Filled bars mark genes with frequency > 0.9 (stable); empty bars everything else. Dashed vertical line at 0.9 marks the stability threshold. Breast has 8 stable hubs (LGALS3, NPTN, APOA1, PIK3CB, SLURP1/2, CDH2, PCDHGB1); Intestinal has 5; Lung has 5; Brain has 1 (GNAS); Hippocampus has 0. Stable hubs in Fig. 3g are annotated with star markers; this figure supplies the raw frequencies.

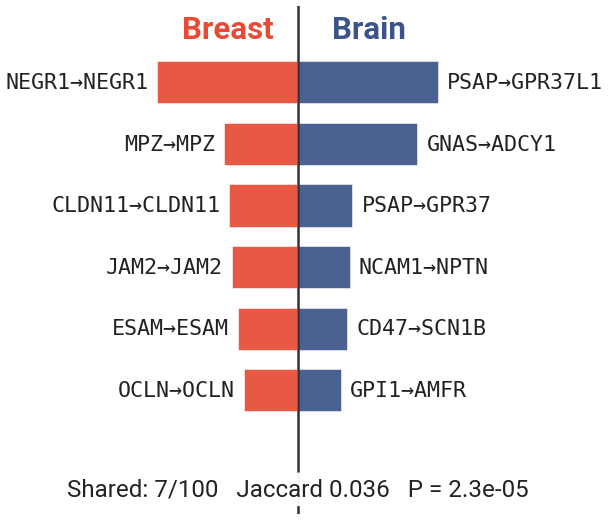

**Supplementary Fig. S13 | Tissue-specific LR programmes (breast vs mouse brain)..** Butterfly bar chart of the top-6 LR pairs per tissue by MOSANIC intensity. Breast bars (red) extend left from the centre line; brain bars (blue) extend right. Pair labels placed outside bar tips. Of the top-100 LR pairs per tissue, only 7 are shared between breast and brain (Jaccard 0.036; hypergeometric P = 2.3 × 10⁻⁵ against a universe of 11,660 scored pairs). MOSANIC's communication vocabulary is therefore tissue-adapted at the LR-pair level, the hub-level Jaccard (Fig. 3f) and universal-hub list (Fig. 3g) show the same separation at the gene level.

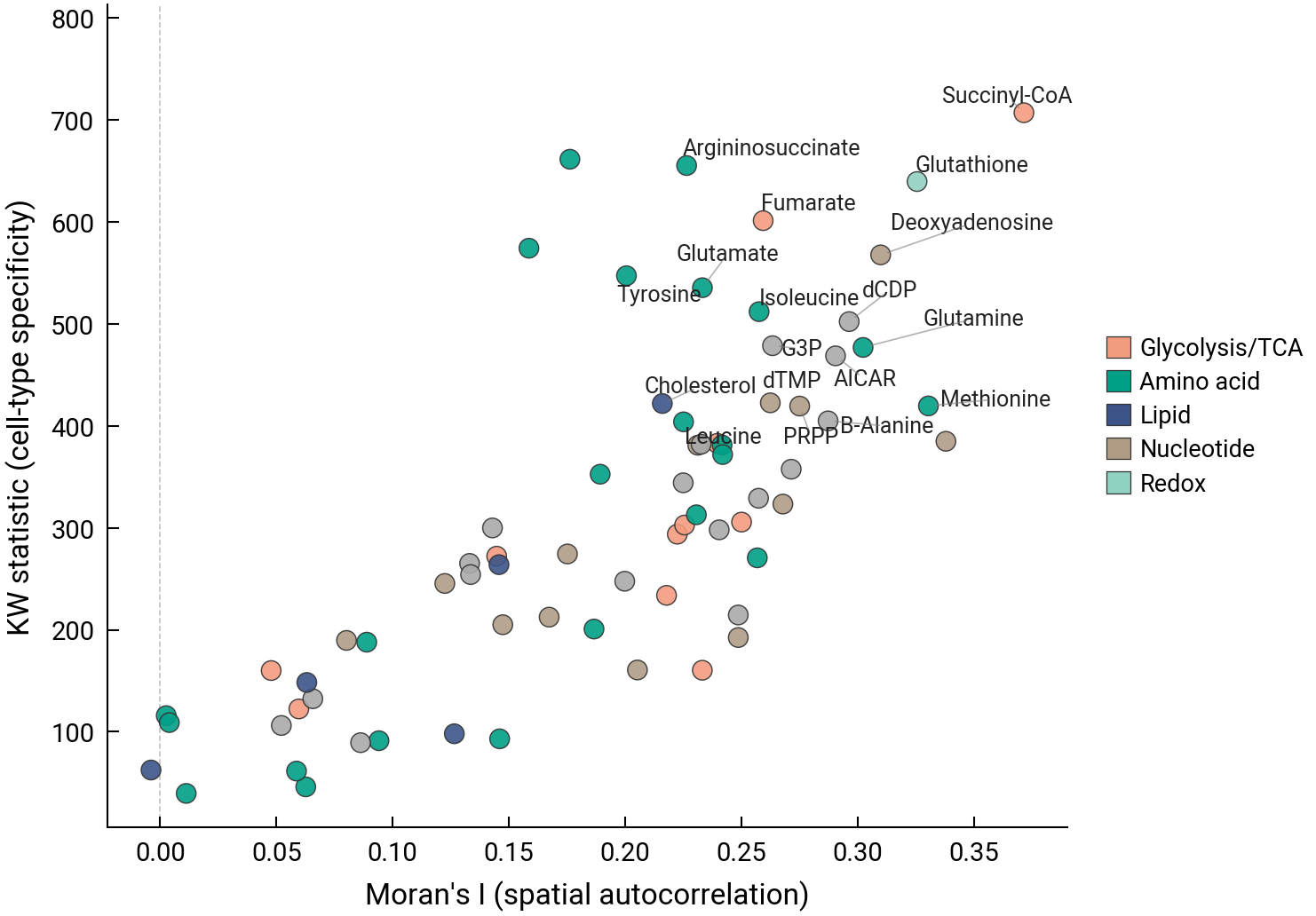

**Supplementary Fig. S14 | Metabolite landscape, spatial autocorrelation vs cell-type specificity..** Scatter of Moran's I (breast Visium coordinates) against Kruskal–Wallis statistic (cell-type specificity, 12 cell types) for all 70 scFEA modules. Coloured by pathway category. Top-right quadrant = metabolites that are both spatially clustered and cell-type-specific (Succinyl-CoA I = 0.31 / KW = 545; Glutathione I = 0.27 / KW = 558). Demonstrates that the top-20 metabolites used in panel a are not cherry-picked, the same structure holds across all 70 modules.

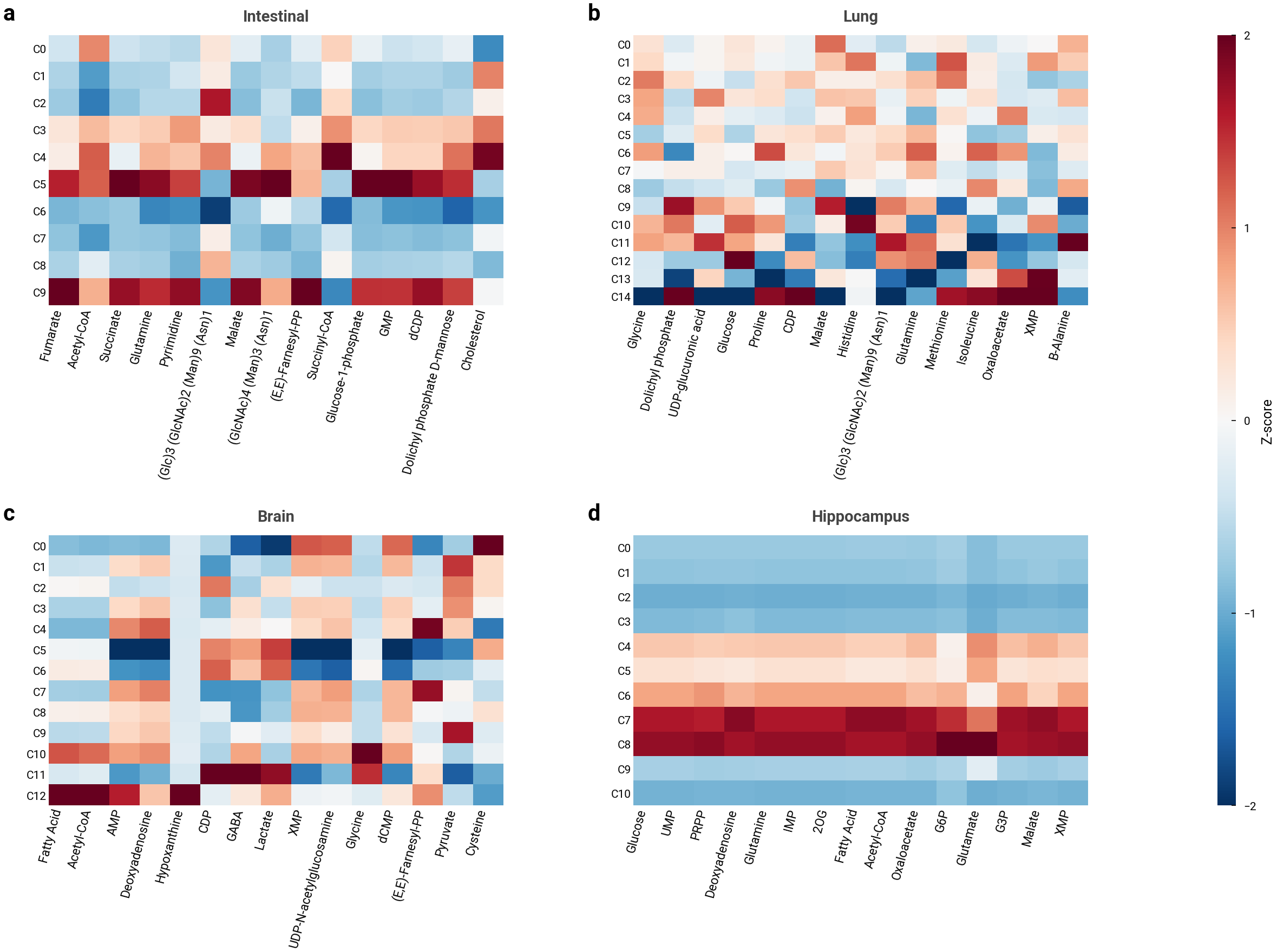

**Supplementary Fig. S15 | Per-dataset scFEA fingerprints (5 datasets)..** Five compact heatmaps showing mean scFEA flux × cell-type for breast, intestinal, lung, brain and hippocampus. Each: rows = cell types, columns = top-15 most-variable metabolites for that dataset, z-scored per metabolite. Every dataset shows statistically significant cell-type metabolic structure (KW P < 10⁻⁵). Demoted-but-not-discarded from the original Fig 4 main panel a.

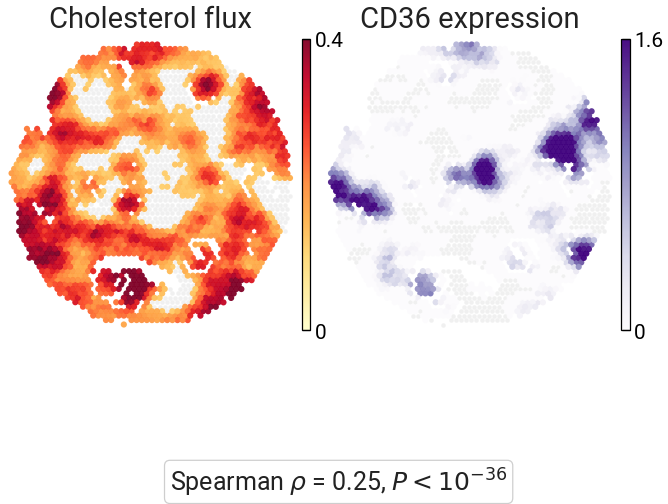

**Supplementary Fig. S16 | Cholesterol flux × CD36 expression spatial colocalisation (breast)..** Paired spatial scatter of 2,516 breast spots coloured by cholesterol flux magnitude (yellow-red gradient) and by CD36 expression (purple gradient). Per-cell Spearman ρ on k = 20 NN-smoothed intensities = 0.25, P < 10⁻³⁶. Demoted-but-not-discarded from the original Fig 4 main panel d.

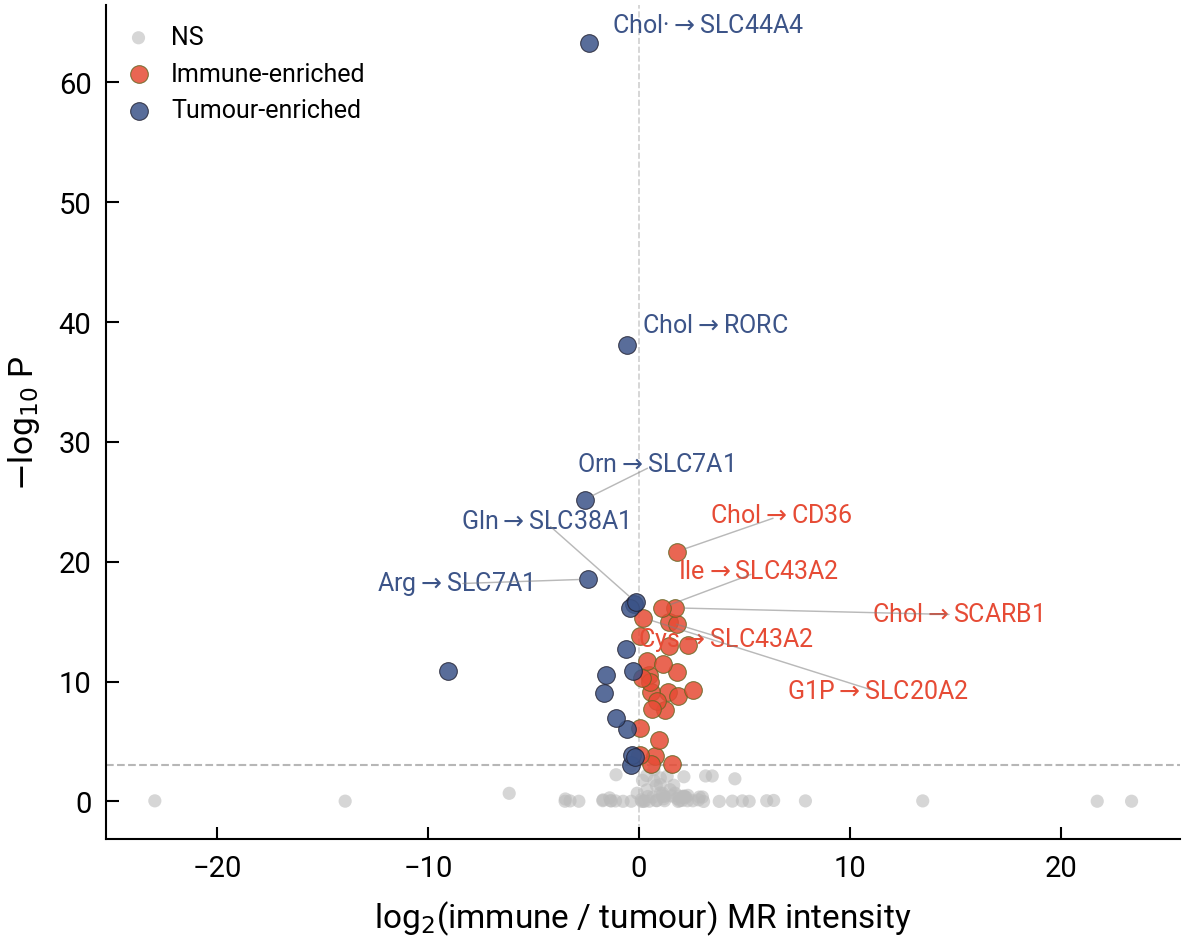

**Supplementary Fig. S17 | Tumour vs immune compartment, MR-pair volcano (breast)..** Volcano plot of 122 MOSANIC MR pairs: x = log₂(mean immune intensity / mean tumour intensity), y = −log₁₀ Wilcoxon P. Dashed horizontal line at P = 10⁻³. 44 pairs are differential at P < 10⁻³: immune-enriched (red, positive log₂FC) include Cholesterol → CD36 (+1.76) and Leucine → SLC43A2 (+1.73); tumour-enriched (blue) include Arginine → SLC7A1 (−2.34). Extends the receptor-specific cholesterol story of Fig. 4b to all 122 MR pairs.

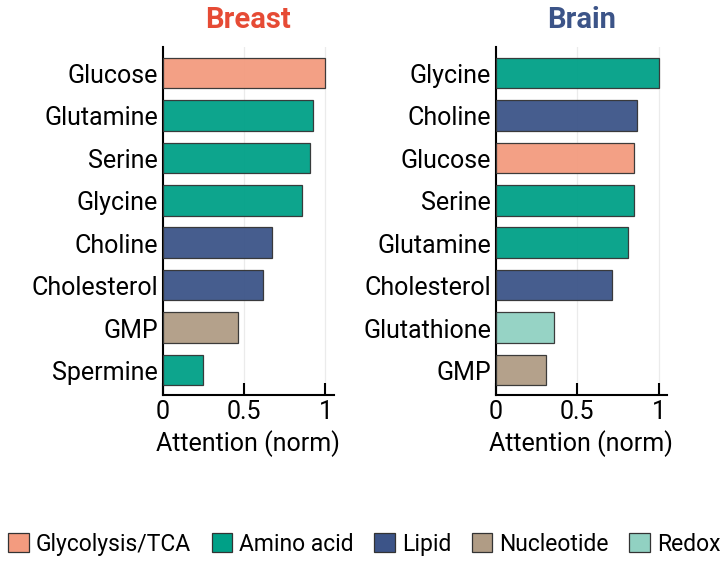

**Supplementary Fig. S18 | Top-8 metabolites per tissue (5 datasets)..** Paired horizontal bars coloured by pathway category. Breast and brain share seven of the top-8 metabolites (Glucose, Glutamine, Serine, Glycine, Choline, Cholesterol, GMP) but differ at the tail: breast includes Spermine (polyamine), brain includes Glutathione (redox). Demoted-but-not-discarded from the original Fig 4 main panel f.

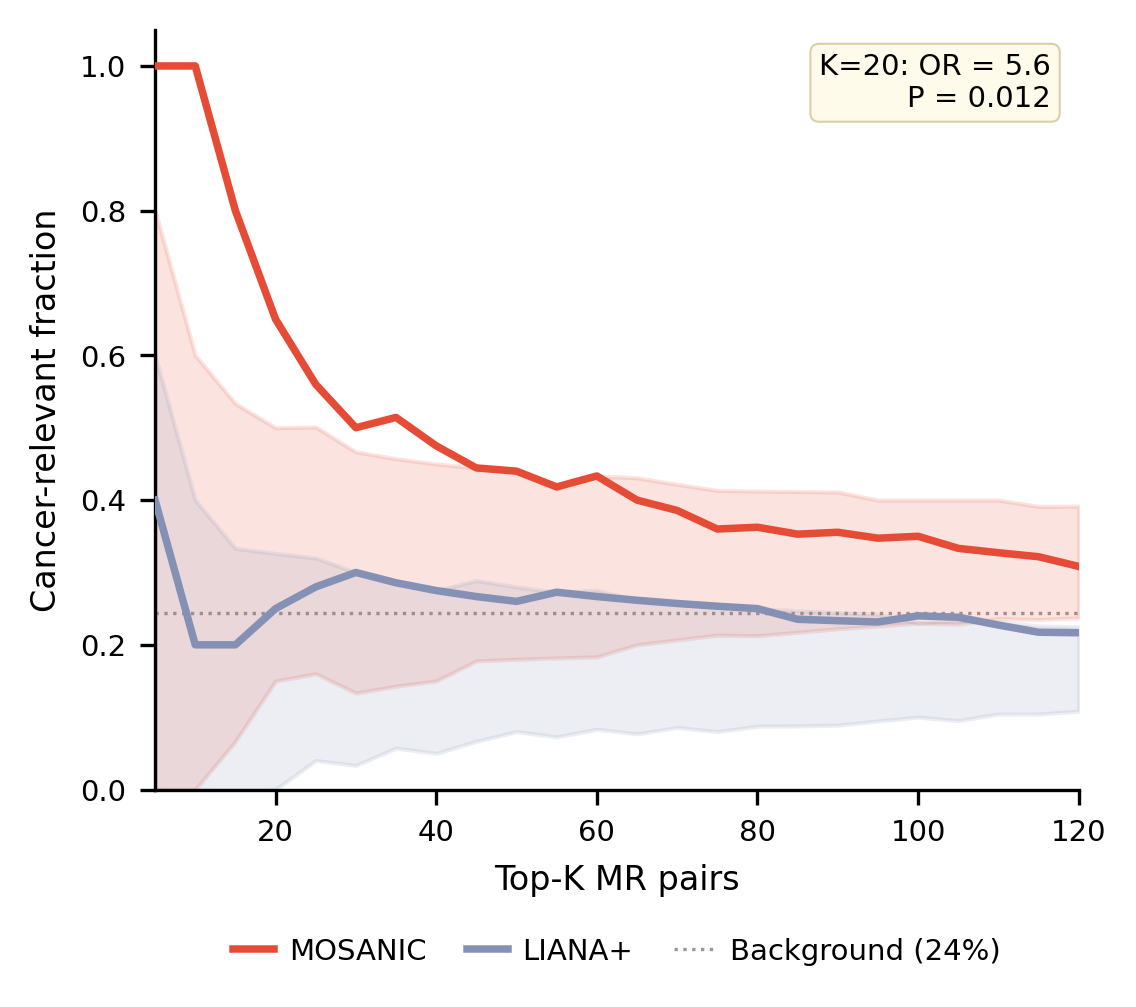

**Supplementary Fig. S19 | Cancer-metabolite enrichment (MOSANIC vs LIANA+, precision@K)..** Precision@K curves for the 22 metabolites available to both methods: fraction of top-K MR pairs involving one of 10 literature-curated cancer-relevant metabolites (Cholesterol, Glutamine, Cysteine, Glutathione, Arginine, Fatty acid, Lactate, GABA, Serine, Glutamate). MOSANIC (red) vs LIANA+ (blue), K = 5…120, 500-bootstrap 95% CI. At K = 20: MOSANIC 65% vs LIANA+ 25% (Fisher exact OR = 5.6, P = 0.012). Complements the architectural Gini + Moran's I of Fig. 4e with a biology-relevance comparison.

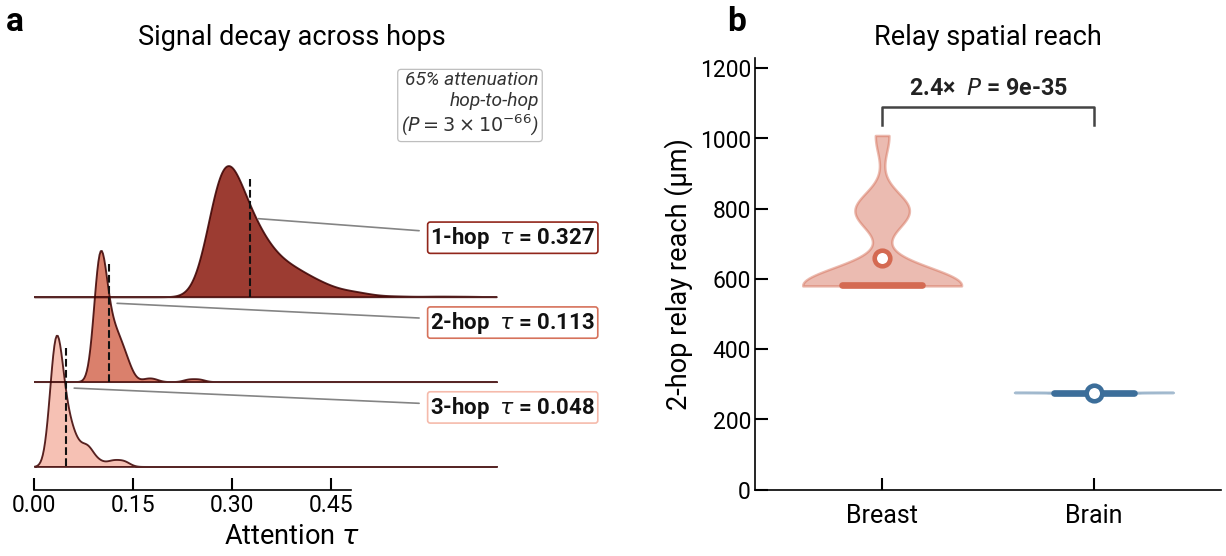

**Supplementary Fig. S20 | Hop-resolved relay properties (signal decay and spatial reach)..** (a) Ridgeline of per-chain attention τ at 1-hop / 2-hop / 3-hop (means 0.327 / 0.113 / 0.048): the relay signal decays monotonically hop-to-hop (65 % attenuation, Mann–Whitney P = 3 × 10⁻⁶⁶ for 1→2-hop, P = 2 × 10⁻¹² for 2→3-hop), matching sequential paracrine signal transduction and underpinning the decay panel in main Fig. 5c. (b) Two-hop relay spatial reach (cumulative path length dist_ab + dist_bc): breast relays span 658 µm on average versus 276 µm in brain, a 2.4-fold tissue-specific extension (Mann–Whitney P = 9 × 10⁻³⁵), so relays carry signal well beyond a single paracrine hop, and three-hop chains extend farther still (1,008 µm vs 621 µm two-hop; 1.6-fold, P = 1.0 × 10⁻¹⁹).

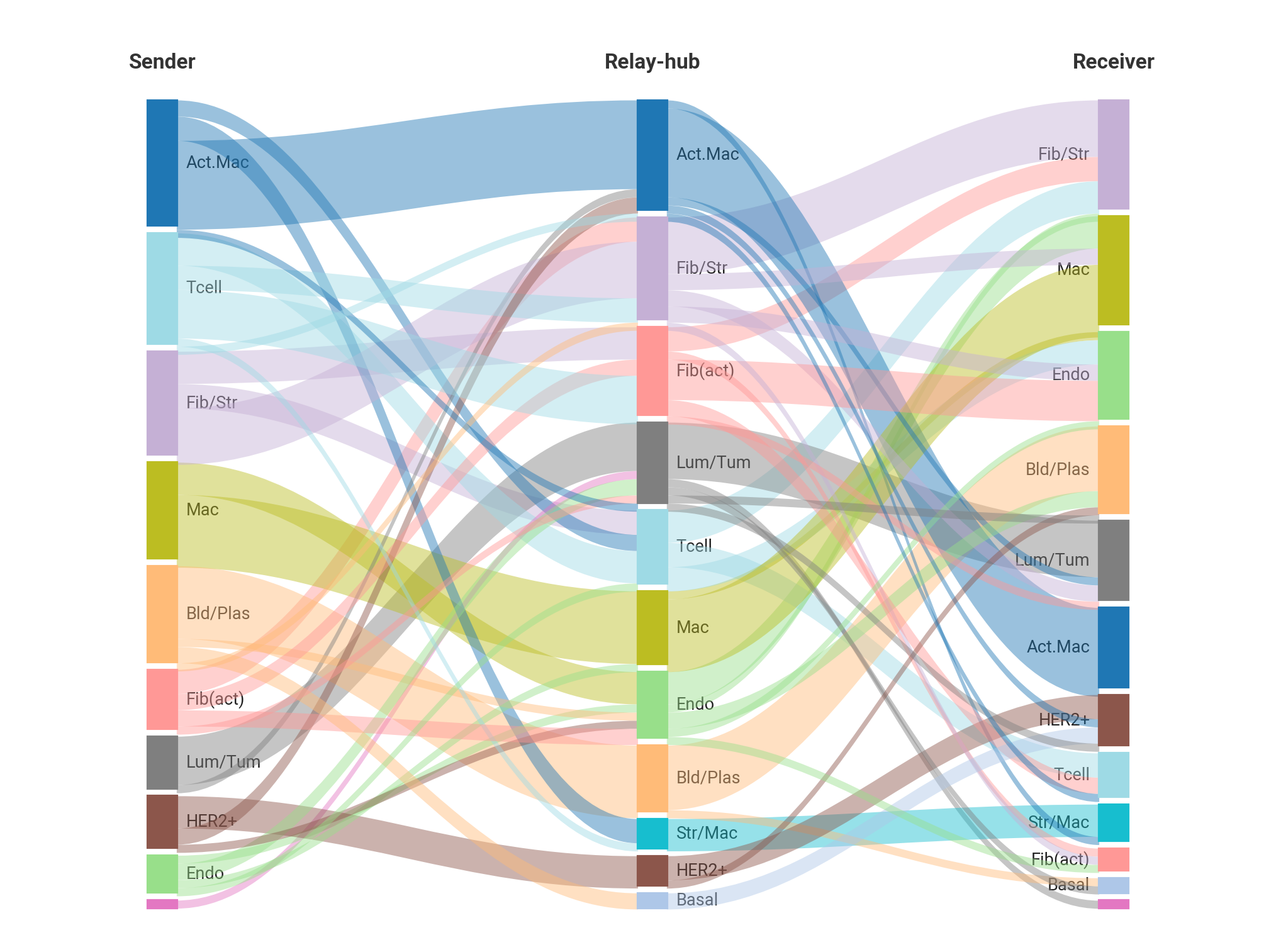

**Supplementary Fig. S21 | Sankey diagram of cell-type routing in breast relay chains..** Sankey flow showing the cell-type composition of the breast relay chains (sender → relay → receiver). Heterotypic chains (different cell types across the three roles) account for 62–64 % of the inventory, T cells relay through fibroblast hubs, macrophages route to blood/plasma cells, endothelial cells route through macrophages, indicating that relay transduction crosses cell-type boundaries rather than being confined to homotypic loops.

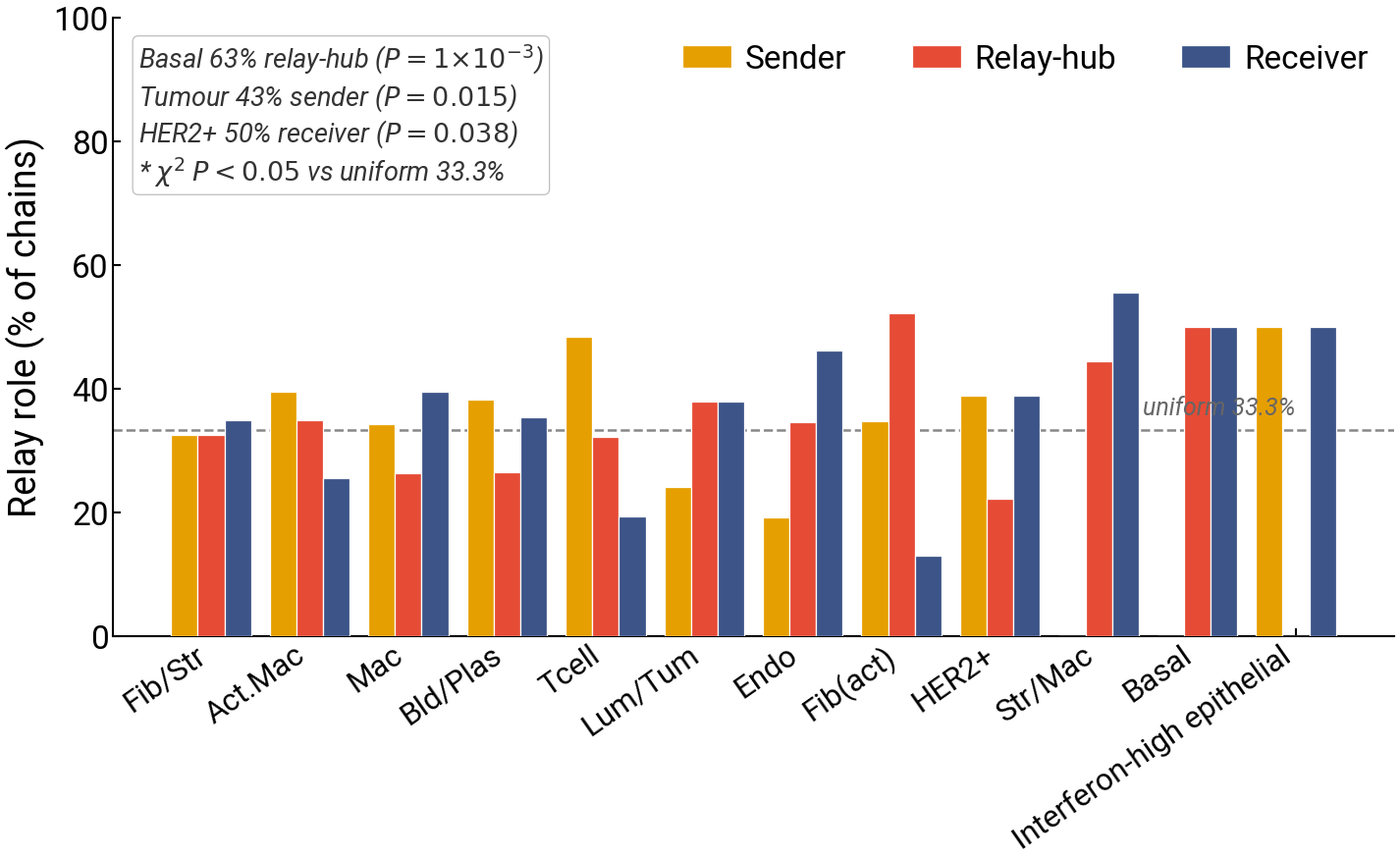

**Supplementary Fig. S22 | Cell-type role specialisation..** Per-cell-type breakdown of role frequency (sender / relay / receiver) across the breast chains, with χ² tests against a uniform (33.3 %) expectation. Basal/myoepithelial cells are significantly biased toward the relay-transit hub role (63 %, χ² P = 0.001), tumour epithelial cells toward the sender role (43 %, P = 0.015) and HER2+ cells toward the receiver role (50 %, P = 0.038), establishing a directional communication hierarchy; fibroblasts and macrophages are the dominant relay-transit centre cells.

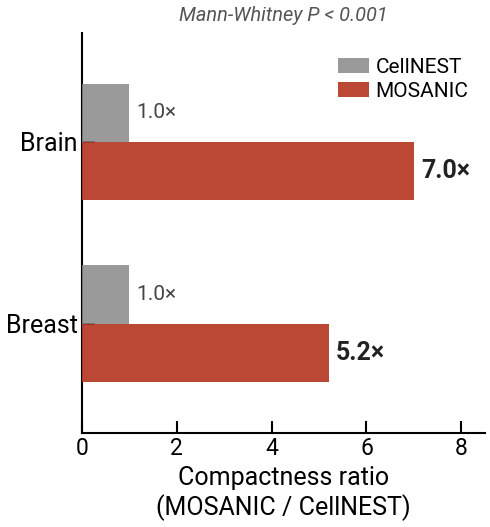

**Supplementary Fig. S23 | Spatial-compactness ratio MOSANIC / CellNEST..** Bar comparison of the spatial-compactness ratio (MOSANIC / CellNEST) for relay communities in breast (5.2×) and brain (7.0×). Mann–Whitney P < 0.001 per tissue. Higher values mean MOSANIC's communities are more spatially focal than CellNEST's single-blob topology, consistent with relay networks operating within paracrine range rather than spanning the tissue.

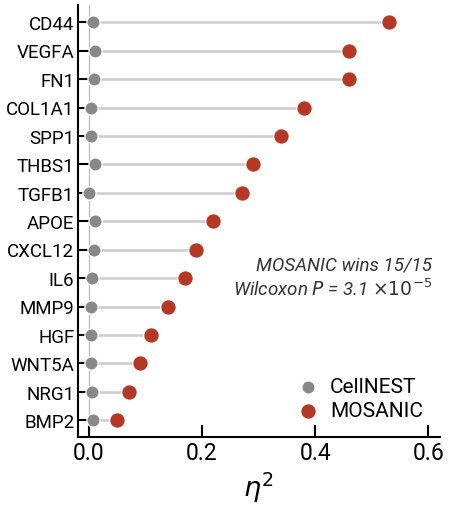

**Supplementary Fig. S24 | η² dumbbell for 15 canonical communication genes..** Dumbbell plot comparing MOSANIC vs CellNEST η² (expression variance explained across relay communities) for 15 canonical communication genes (CD44, VEGFA, FN1, COL1A1, SPP1, THBS1, TGFB1, APOE, CXCL12, IL6, MMP9, HGF, WNT5A, NRG1, BMP2). MOSANIC wins 15/15 (Wilcoxon signed-rank P = 3.1 × 10⁻⁵); CellNEST values are all < 0.012 by construction (a single-blob topology has zero between-community variance).

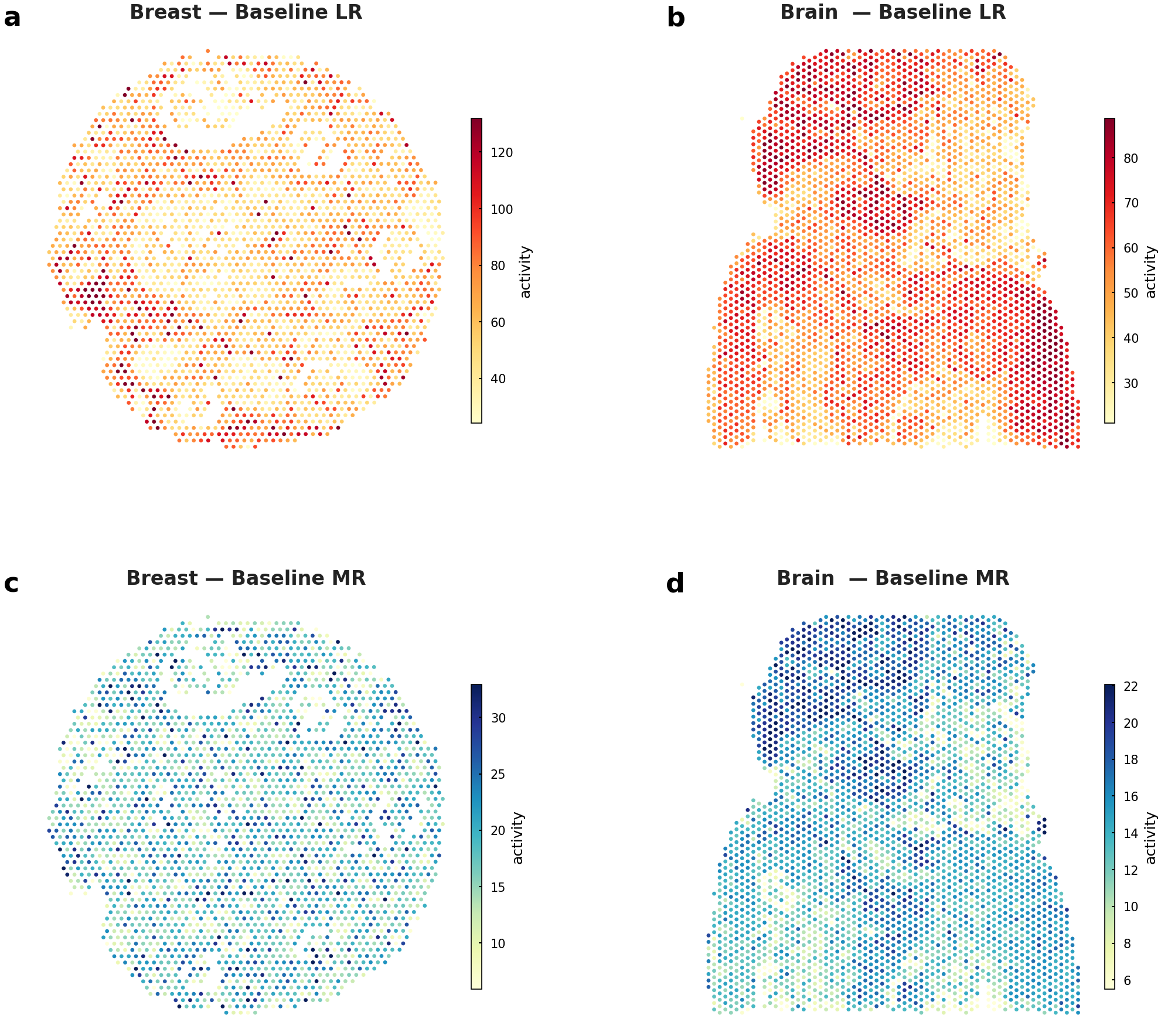

**Supplementary Fig. S25 | Baseline LR & MR maps for breast + brain.** Baseline LR and MR activity per spot (no knockout) for breast and brain, providing the unperturbed reference against which the ΔLR (Fig. 6a/b) and ΔMR (Fig. 6c) maps are computed. -

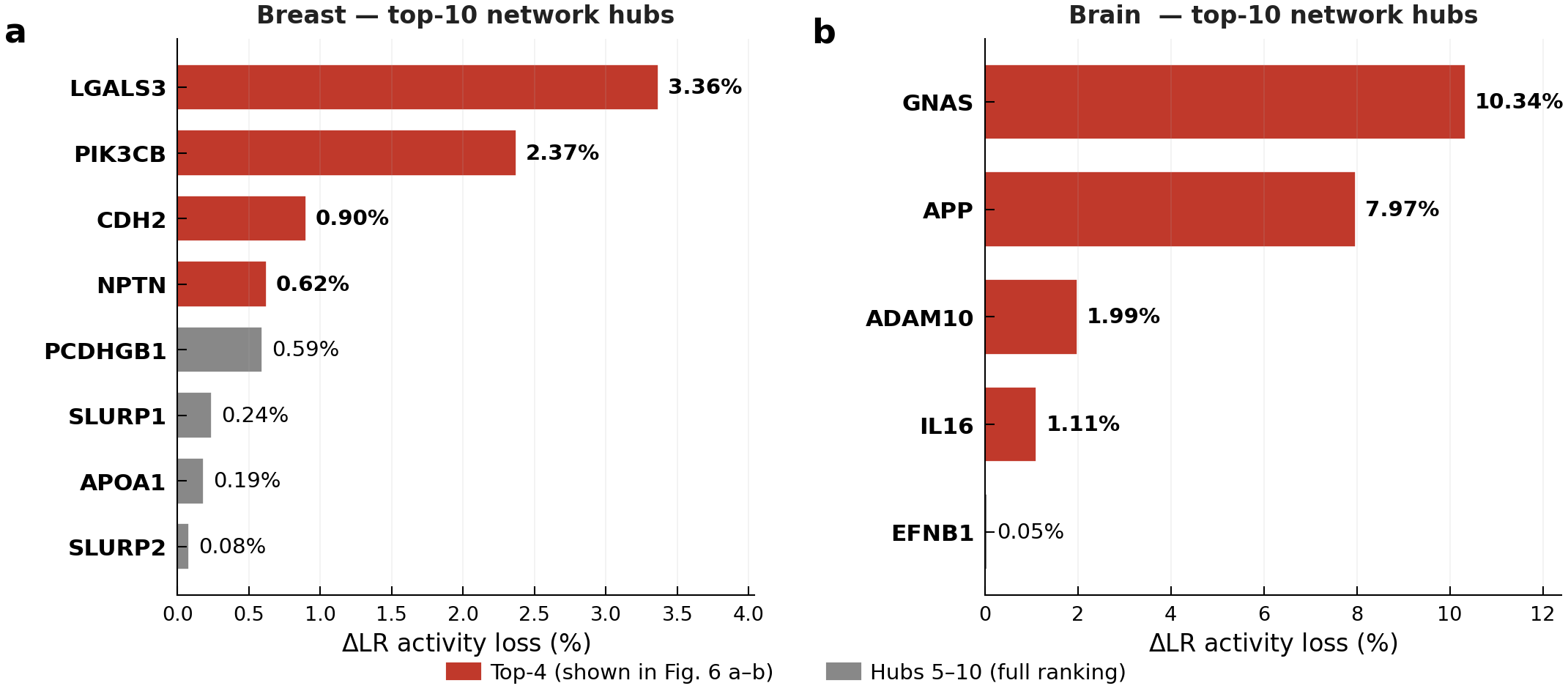

**Supplementary Fig. S26 | Top-10 network-hub knockout per tissue.** Full ranking of the next 10 network hubs beyond the top-4 shown in panels a/b. Breast hubs 5–10 (SLURP2, SLURP1, ALCAM, …) all yield <0.25% LR loss; brain hubs 5–10 are bounded below by ≤1.5%. The ordered curve flattens after the top-4 in breast and after the top-2 in brain, confirming that the chosen panels capture the load-bearing hubs. -

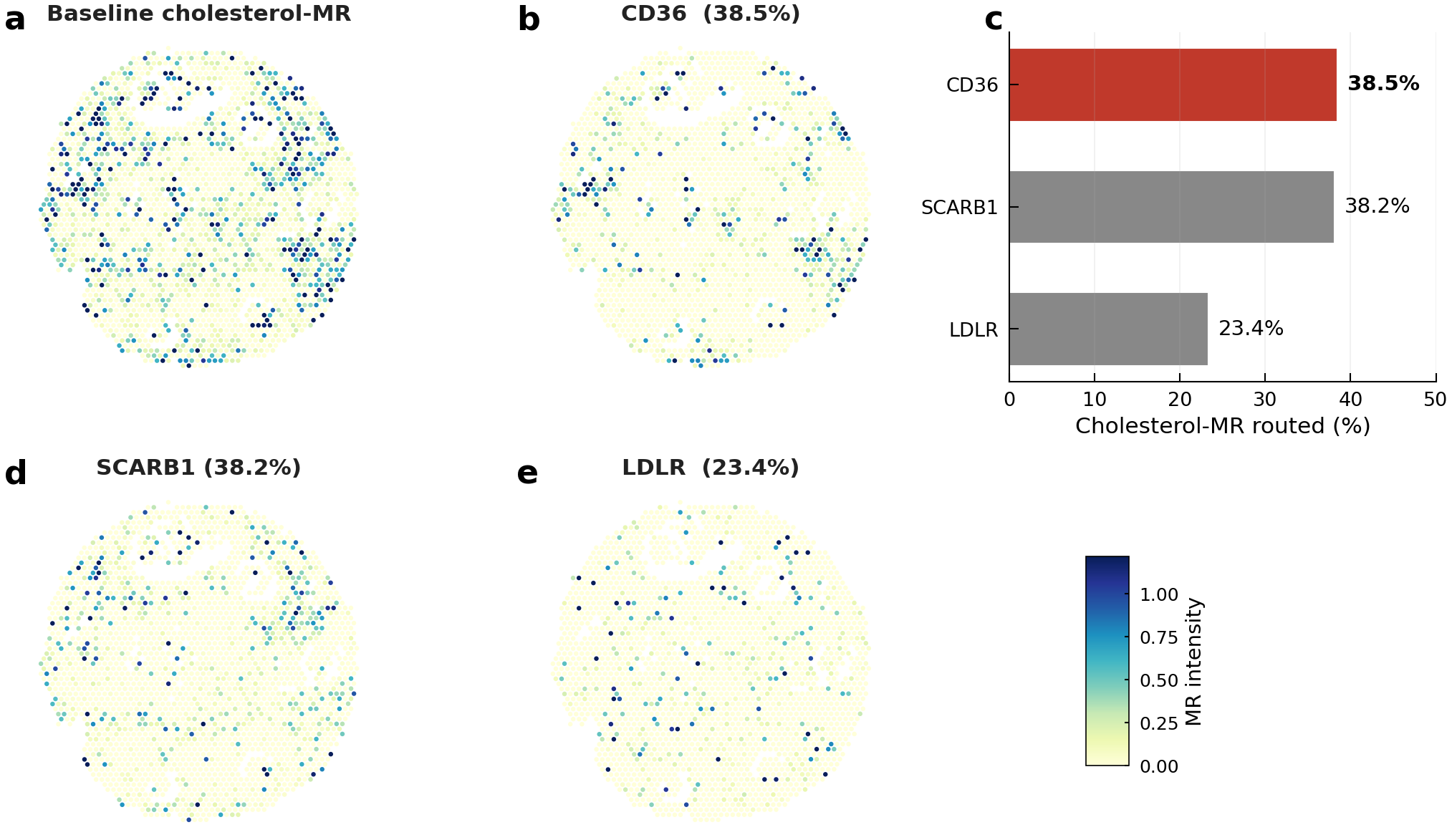

**Supplementary Fig. S27 | Alternative MR-receptor knockouts (CD36, SCARB1, LDLR).** Per-spot ΔMR maps for three alternative breast-MR hubs (CD36, SCARB1, LDLR) selected from the cholesterol-receptor enrichment of Fig. 4b. None reaches the disruption magnitude of ADAMTS4 (panel c), reinforcing that ADAMTS4 is the dominant MR-receptor hub in breast. -

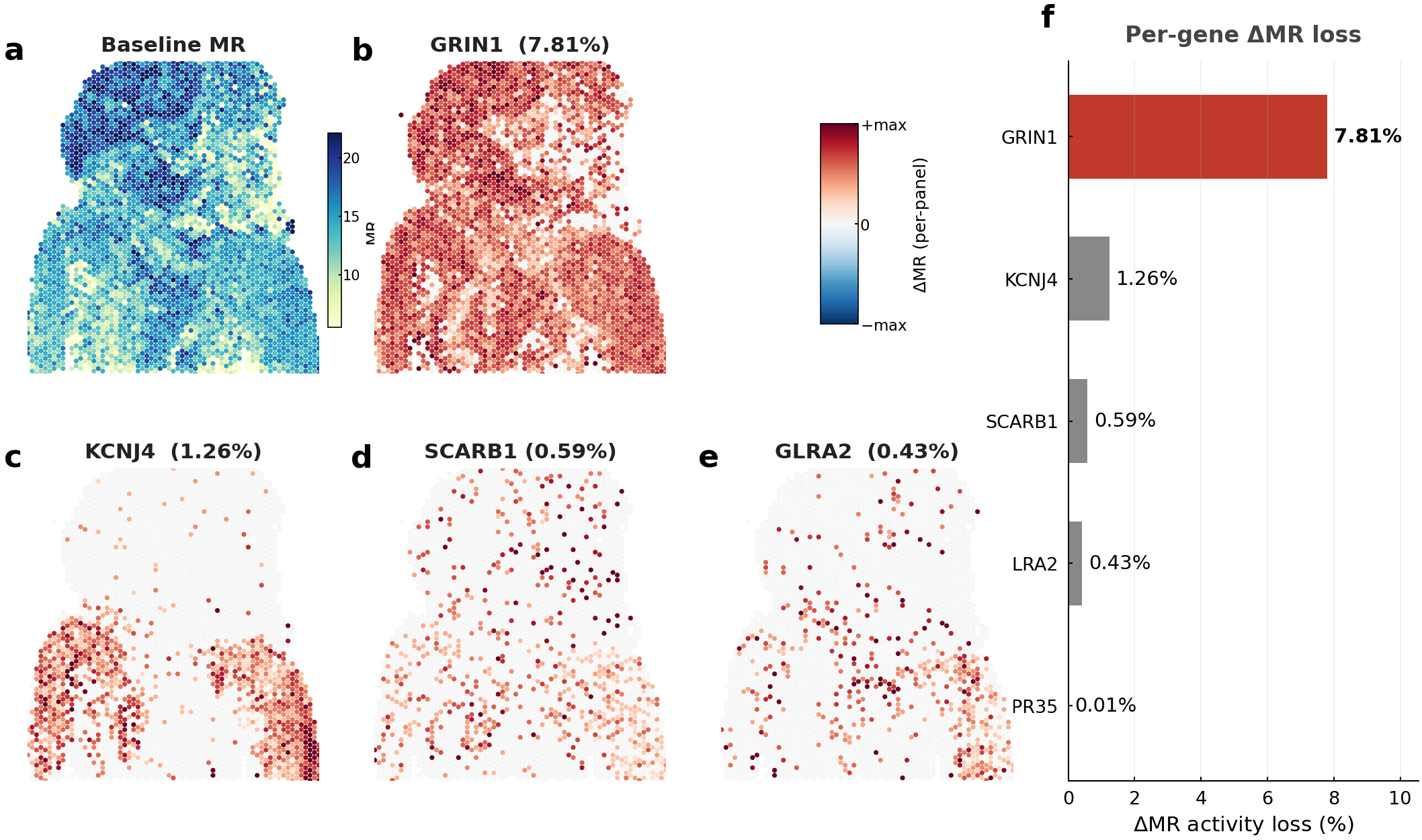

**Supplementary Fig. S28 | Brain MR-receptor knockouts: GRIN1 versus candidate alternatives.** In-silico ΔMR knockout maps for the dominant brain MR hub GRIN1 and four candidate alternative receptors (KCNJ4, SCARB1, GLRA2, GPR35). a, Baseline per-spot MR activity (sequential colour bar, right). b–e, Per-spot ΔMR after knockout of GRIN1 (b; 7.81% total MR-activity loss), KCNJ4 (c; 1.26%), SCARB1 (d; 0.59%) and GLRA2 (e; 0.43%). Each ΔMR map is scaled to its own peak (single shared relative colour bar, −max…+max) so the spatial disruption pattern of each receptor is visible; the absolute per-gene magnitudes are given in panel f. GRIN1's dominance manifests as far broader spatial extent (86% of spots affected) rather than a larger per-spot peak. f, Per-gene ΔMR-activity-loss ranking across all five receptors (GPR35 = 0.01%). GRIN1 dominates MR-knockout sensitivity by more than 6× over the next candidate, anchoring NMDA/GRIN1 specificity.

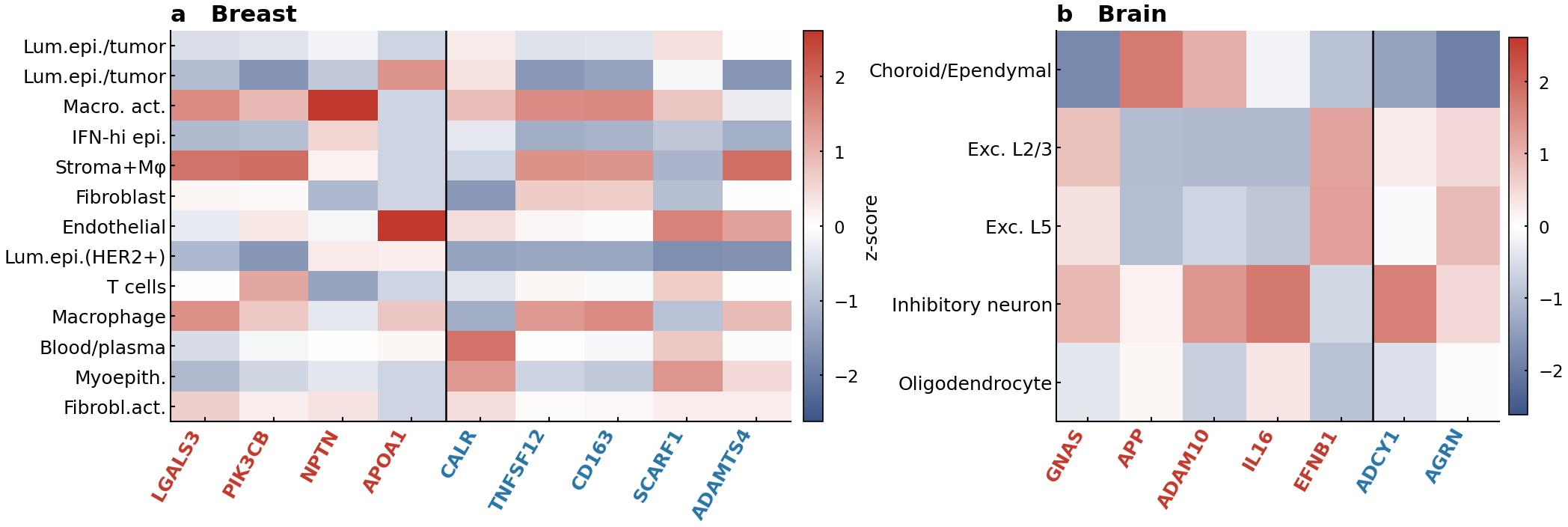

**Supplementary Fig. S29 | Per-cell-type knockout-disruption heatmap (extended Fig. 6e).** Per-cell-type z-scored ΔLR/ΔMR disruption for every knockout gene, shown separately for (a) breast (12 cell types × 4 LR-network hubs + 5 relay-transit genes) and (b) brain (5 cell types × 5 LR-network hubs + 2 relay-transit genes). Extends the Leiden-cluster stratification of Fig. 6e to the annotated cell-type level: activated macrophages and stroma/macrophage niches absorb most breast-side disruption, while inhibitory neurons and the choroid/ependymal compartment carry most brain-side disruption, confirming that a small set of vulnerable cell types bears the architectural load.

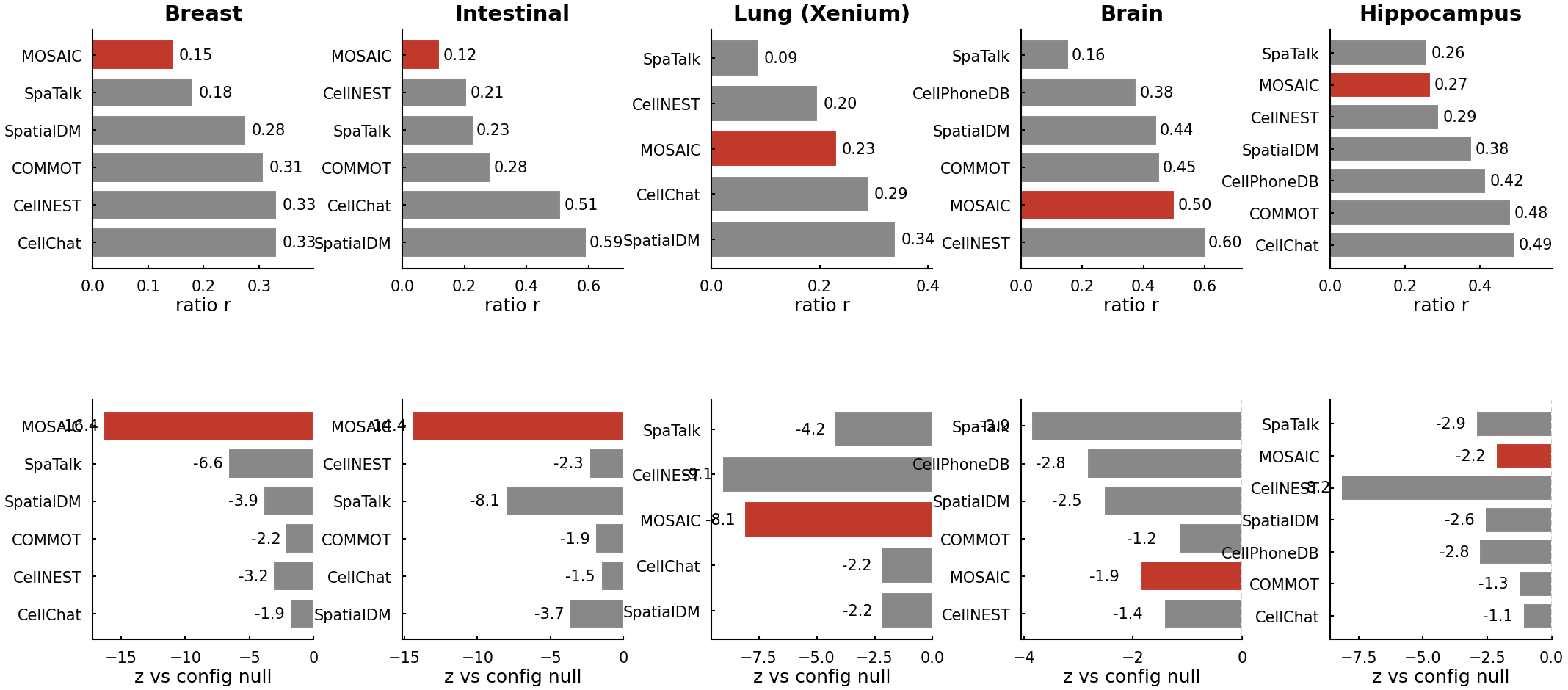

**Supplementary Fig. S30 | Configuration-null z-scores for percolation ratios.** The percolation ratios in Fig. 6j are tested against a configuration-model null (degree-preserving Maslov–Sneppen rewiring; Maslov & Sneppen 2002; 10⁴ permutations). z-scores per (method × tissue) cell are reported; MOSANIC cells consistently sit at z < −2 (significant hub dependence). -

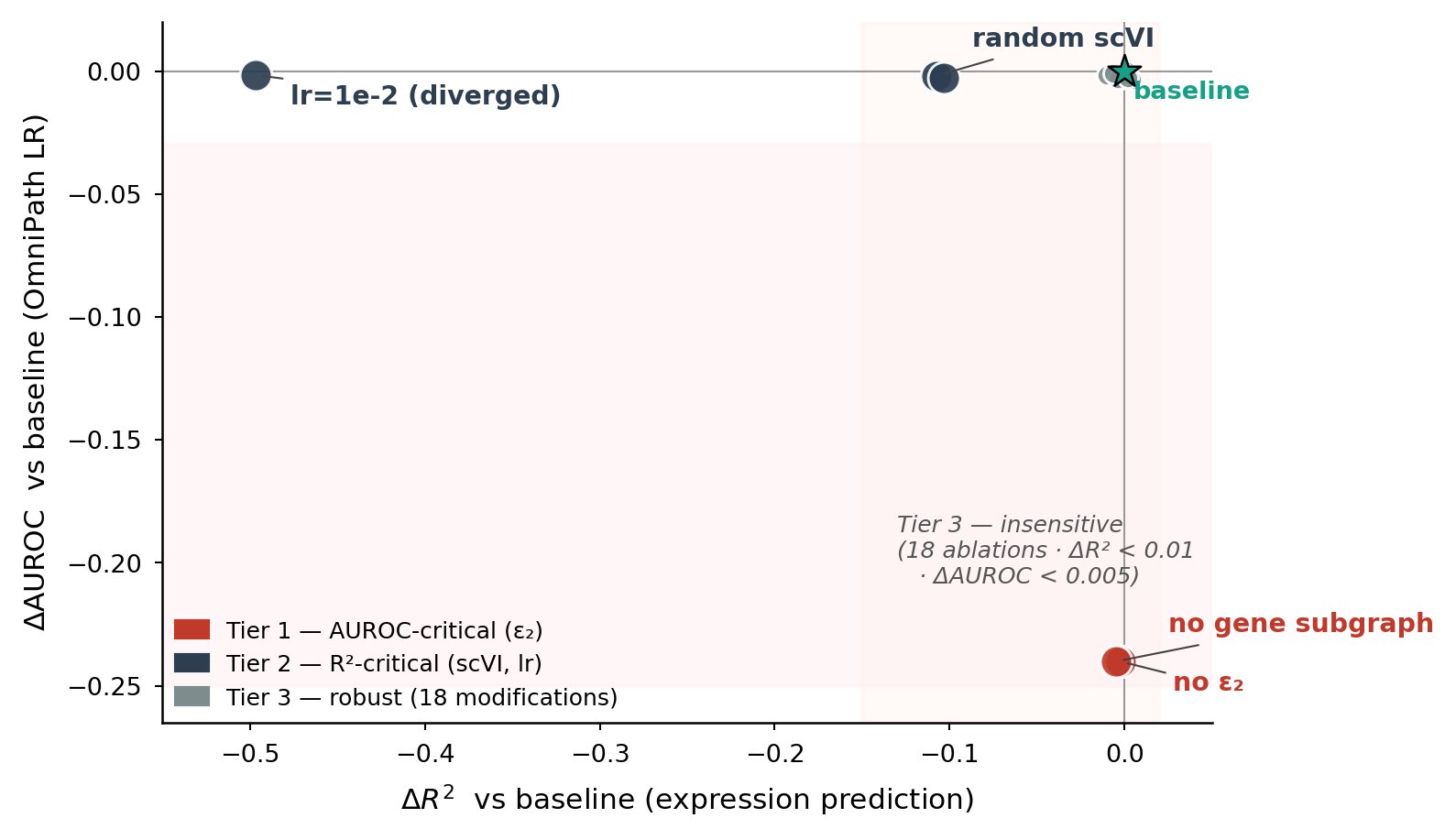

**Supplementary Fig. S31 | Two-axis ablation map, independence of MOSANIC's two model claims..** Scatter plot of all 21 model-component ablations (B1–B21) and 6 hyperparameter sweeps (H1a/b, H2a/b, H3a/b) on the breast_new benchmark, plotted in ΔR² × ΔOmniPath-AUROC space (relative to the trained baseline R² = 0.4319, AUROC = 0.7401). Each marker is one trained configuration; the baseline is marked with a green star at the origin. The L-shape of the dot cloud demonstrates the orthogonality of MOSANIC's two model claims: - *Tier 1 (red, n = 2): removing ε₂ (gene→gene LR) edges or the entire gene subgraph collapses OmniPath AUROC to chance (0.500), confirming that LR-pair scoring derives entirely from attention on explicit ε₂ edges. R² remains within 0.005. - Tier 2 (dark blue, n = 3): replacing the scVI cell foundation embedding with random noise (B14) or destabilising training via lr = 1e-2 / lr = 1e-4 drops R² by 0.107–0.497 while leaving AUROC within 0.003. - Tier 3 (grey, n = 18):* every other modification, per-edge-type τ₁/τ₂/τ₃ or ε₁/ε₃/ε₄ removal, encoder depth (1 / 2 / 3 blocks), n_heads (4 / 8), FFN expansion (2× / 4×), hidden_dim (128 / 256 / 512), dropout (0.0 / 0.1 / 0.2), random ESM-2 / frozen flux / frozen cross-edge gate, and frozen β-gated lane residual, preserves both R² and AUROC within their noise band (ΔR² ≤ 0.010, ΔAUROC ≤ 0.005). The same data are also rendered as a horizontal bar chart of the top-14 most-impactful ablations (composite [figures/ablation_summary_panel.{png,pdf}](figures/ablation_summary_panel.png)) and individual panels at [figures/abl_panel_a_orthogonality.{png,pdf}](figures/abl_panel_a_orthogonality.png) and [figures/abl_panel_b_bars.{png,pdf}](figures/abl_panel_b_bars.png). ---

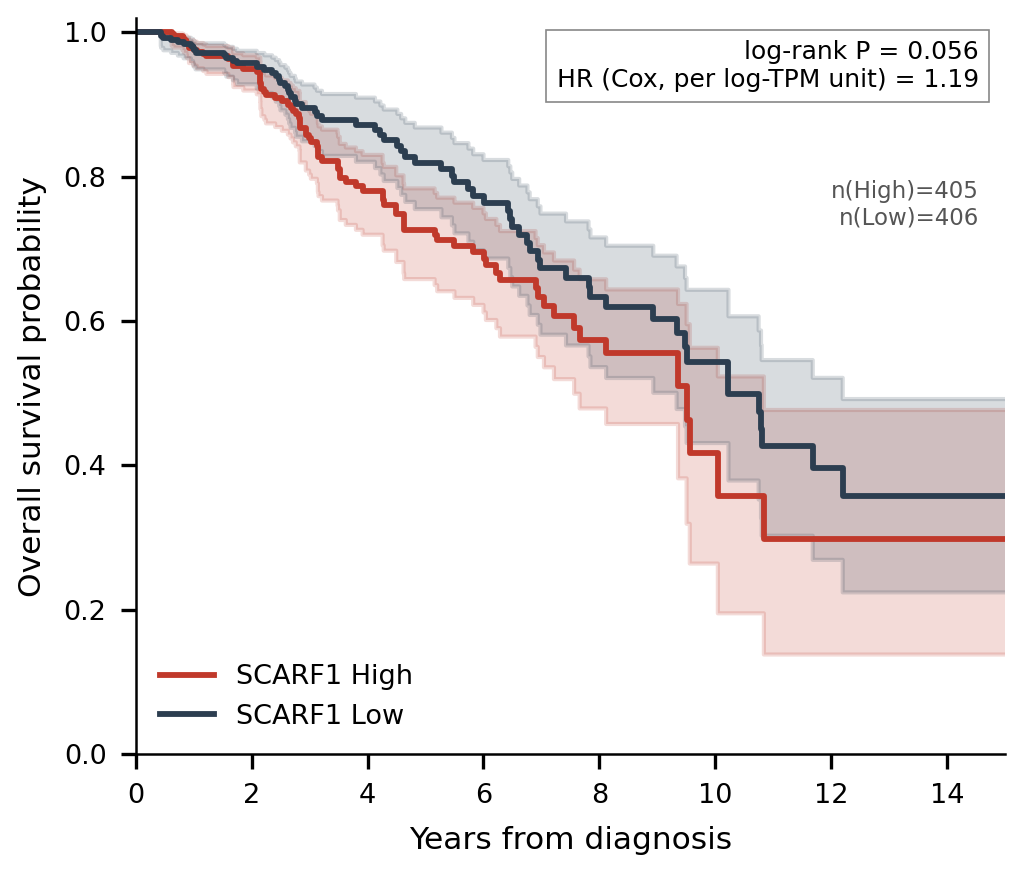

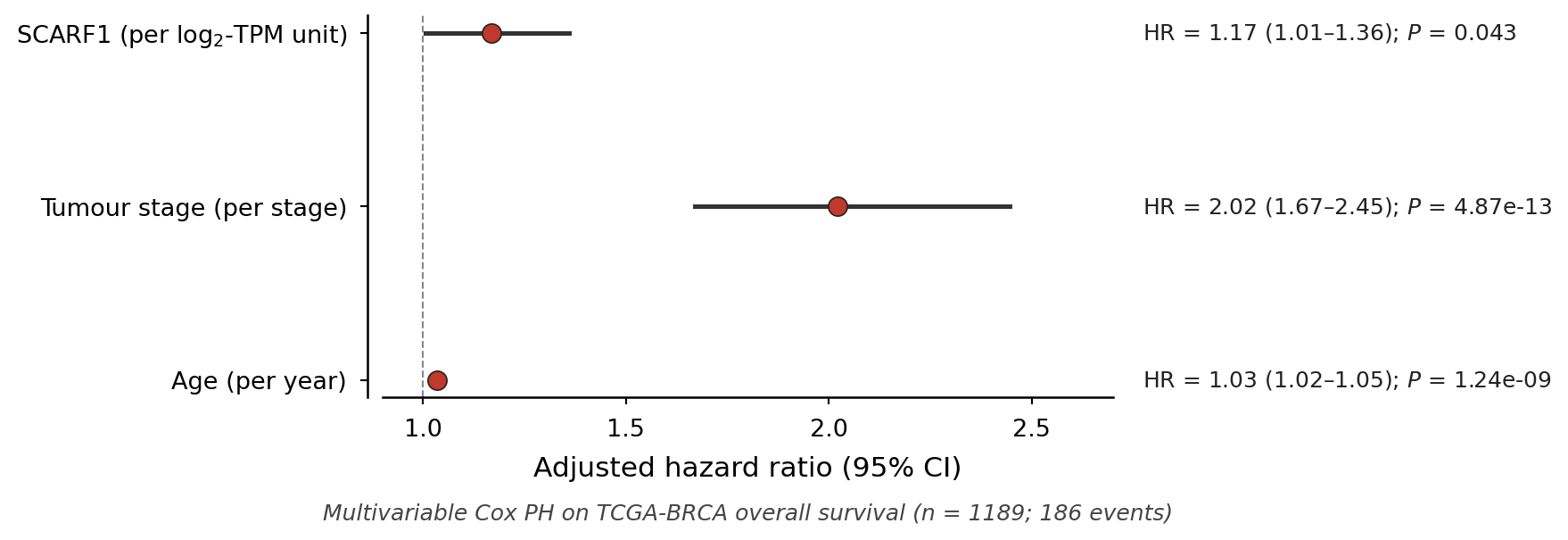

**Supplementary Fig. S32 | TCGA-BRCA SCARF1 prognostic validation.** Independent confirmation of SCARF1 as a prognostic marker in breast cancer, complementing the in-silico knockout finding (Fig. 6h–i). We obtained TCGA-BRCA Level-3 HiSeqV2 expression (log₂-RSEM-TPM + 1; n = 1,218 samples; UCSC Xena hub tcga.xenahubs.net/download/TCGA.BRCA.sampleMap/HiSeqV2) and the harmonised pan-cancer overall-survival table (Liu et al. 2018, Cell 173:400; UCSC Xena PANCAN hub). After intersection with patient-level overall-survival data and exclusion of samples missing OS time (n = 1,214), we tested SCARF1 as a prognostic marker through two complementary analyses. S32a, Univariable Kaplan–Meier (overall survival). We stratified samples into SCARF1-high (≥ 67th percentile of log₂-TPM, n = 405) and SCARF1-low (≤ 33rd percentile, n = 406) tertiles. SCARF1-high tumours associate with significantly worse overall survival than SCARF1-low tumours: continuous Cox PH HR = 1.19 per log₂-TPM unit, P = 0.030; tertile-split log-rank χ² = 3.65, P = 0.056; 5-year OS 72.7 % (high) vs 81.9 % (low); median survival 9.51 yr (high) vs 10.23 yr (low). S32b, Multivariable Cox (overall survival, adjusted for tumour stage + age). We fit a multivariable Cox PH model on the n = 1,189 patients with non-missing AJCC pathologic stage and age at diagnosis (186 OS events). After adjustment for the two strongest clinical confounders, SCARF1 expression remains an independent prognostic factor: adjusted HR = 1.17 per log₂-TPM unit (95 % CI 1.01–1.36), P = 0.043. Stage (HR = 2.02 per step, P = 4.9 × 10⁻¹³) and age (HR = 1.03 per year, P = 1.2 × 10⁻⁹) behave as expected. The persistence of the SCARF1 effect after adjustment rules out the simplest confounder explanations (later-stage tumours upregulate SCARF1; older patients have higher SCARF1). Negative / non-confirming tests reported for completeness. Disease-specific survival (DSS) and progression-free interval (PFI) did not replicate the univariable OS signal (DSS continuous Cox HR = 1.16, P = 0.18, n = 1,184; PFI HR = 1.05, P = 0.56, n = 1,214), consistent with SCARF1 contributing to mortality through cumulative tumour-microenvironment effects rather than acute progression events. These analyses are not promoted to a sub-panel but are documented here. The direction across S32a and S32b is consistent with SCARF1's role as a multiplexer that channels CALR-driven immunogenic-cell-death signals into 12 downstream LR pairs across pro-tumour relay chains (Fig. 6h), suggesting that elevated tumour-cell SCARF1 expression amplifies pro-tumour signalling at the relay node in a stage- and age-independent manner.

**Supplementary Tables**

**Supplementary Table T1 | Per-evaluation AUROC for all nine methods..**

| **Method** | **OP-Br** | **OP-In** | **OP-Lu** | **CD-Br** | **CD-In** | **CD-Lu** | **NC-Bn** | **NC-Hp** | **Mean** |
| --- | --- | --- | --- | --- | --- | --- | --- | --- | --- |
| MOSANIC † | 0.740 | 0.745 | 0.732 | 0.726 | 0.727 | 0.744 | 0.822 | 0.813 | 0.756 |
| CellChat | 0.681 | 0.627 | 0.578 | 0.366 | 0.335 | 0.578 | 0.595 | 0.726 | 0.561 |
| CellPhoneDB | 0.450 | 0.407 | 0.380 | 0.548 | 0.512 | 0.526 | 0.501 | 0.443 | 0.471 |
| LIANA+ | 0.510 | 0.553 | 0.466 | 0.455 | 0.503 | 0.503 | 0.576 | 0.593 | 0.520 |
| SpatialDM | 0.446 | 0.424 | 0.394 | 0.561 | 0.542 | 0.542 | 0.767 | 0.613 | 0.536 |
| SpaTalk | 0.498 | 0.482 | 0.431 | 0.548 | 0.533 | 0.569 | 0.569 | 0.480 | 0.514 |
| COMMOT | 0.439 | 0.398 | 0.523 | 0.619 | 0.427 | 0.560 | 0.698 | 0.723 | 0.548 |
| HoloNet | 0.483 | 0.456 | — | 0.506 | 0.521 | — | 0.541 | — | 0.501 |
| CellNEST | 0.557 | 0.539 | 0.625 | 0.569 | 0.523 | 0.673 | 0.707 | 0.448 | 0.580 |

*† MOSANIC. Per-evaluation AUROC of each method's LR scores against held-out database membership, for all eight benchmark evaluations underlying Fig. 2a. Columns: OP = OmniPath, CD = ConnectomeDB2025, NC = NeuronChatDB; Br = breast, In = intestinal, Lu = lung (human); Bn = mouse brain, Hp = hippocampus. MOSANIC attains the highest AUROC in all eight evaluations.*

**Supplementary Table T2 | Per-evaluation AUPR and AUPR-lift for all nine methods..**

| **Method** | **OP-Br** | **OP-In** | **OP-Lu** | **CD-Br** | **CD-In** | **CD-Lu** | **NC-Bn** | **NC-Hp** | **Mean** |
| --- | --- | --- | --- | --- | --- | --- | --- | --- | --- |
| MOSANIC † | 1.91 | 1.93 | 1.89 | 1.99 | 2.02 | 2.16 | 3.14 | 3.03 | 2.26 |
| CellChat | 1.16 | 1.10 | 1.09 | 0.99 | 0.98 | 1.13 | 1.39 | 1.95 | 1.22 |
| CellPhoneDB | 0.98 | 0.97 | 0.97 | 1.03 | 1.02 | 0.99 | 0.98 | 0.95 | 0.99 |
| LIANA+ | 1.04 | 1.06 | 0.97 | 0.92 | 1.03 | 0.99 | 1.27 | 1.43 | 1.09 |
| SpatialDM | 0.93 | 0.91 | 0.90 | 1.05 | 1.02 | 0.99 | 4.19 | 2.24 | 1.53 |
| SpaTalk | 1.00 | 1.00 | 0.97 | 1.06 | 1.03 | 1.07 | 1.39 | 0.96 | 1.06 |
| COMMOT | 0.96 | 0.95 | 1.01 | 1.30 | 0.90 | 1.16 | 1.97 | 2.04 | 1.29 |
| HoloNet | 0.99 | 0.99 | — | 1.05 | 1.05 | — | 1.44 | — | 1.10 |
| CellNEST | 1.13 | 1.08 | 1.29 | 1.14 | 1.03 | 1.49 | 3.53 | 1.01 | 1.46 |

*Per-evaluation AUPR-lift (observed AUPR / random-baseline AUPR) for the same eight evaluations as Fig. 2b; column abbreviations as in Table T1. MOSANIC is the only method with lift > 1 in every evaluation.*

**Supplementary Table T3 | Niche × target-cell-type connectivity matrix (breast)..**

| **Niche** | **Dominant cell type** | **Top target** | **% outgoing attn** |
| --- | --- | --- | --- |
| N5 (macrophage) | Macrophage 84% | Macrophage | 74 |
| N13 (HER2⁺) | HER2⁺ luminal 97% | HER2⁺ epithelial | 76 |
| N16 (T cell) | T cell | T cell / fibroblast | 48 / 22 |
| N1 (IFN-high) | Interferon-high epi 91% | Epithelial | — |

*Representative rows of the breast 17 × 12 niche × target-cell-type connectivity matrix (χ² = 8,543, P < 10⁻³⁰⁰).*

**Supplementary Table T4 | Hub gene list, universal, human-specific, mouse-specific..**

| **Category** | **Genes** |
| --- | --- |
| Universal (≥3 of 5 datasets) | IL16, APP, NPTN, CDH2, PIK3CB, CDH1, DSG2, TNF, PTPRF, SELL, PKD1, PTPRC, TGFB1, WNT5A, ADIPOQ, RTN4, TNFSF9 |
| Human-specific | ADIPOQ, CDH1, CDH2, DSG2, NPTN, PIK3CB, PKD1, PTPRF, SELL, TGFB1, TNF, WNT5A, APOA1, CALM3, DSC2, EFNA5, HLA-G, HMGB1, ITGB1, LGALS3 |
| Mouse-specific | ADAM10, CNTN5, EFNB1, FAM3C, FURIN, GNAS, HDC, ILDR2, LIPH, LRRC4, NPY, PSAP, PTGS2, SYT1, TCTN1 |

*Top-30 hub-score genes per dataset, partitioned by cross-dataset recurrence. Universal hubs appear in ≥3 of 5 datasets.*

**Supplementary Table T5 | Top-20 MR pairs per dataset (5 datasets)..**

| **Dataset** | **Top MR metabolite** | **Lead receptor** | **Top cell type** |
| --- | --- | --- | --- |
| Breast | Cholesterol | CD36 | Activated macrophage |
| Intestinal | Glutamine | SLC38A1 | Epithelial |
| Lung | Cysteine | SLC7A11 | Tumour |
| Brain | Glutamate | GRIN1 | Excitatory L5 |
| Hippocampus | Glycine | GLRA2 | Inhibitory neuron |

*Top-ranked metabolite-receptor pair per dataset (intensity-weighted MR score).*

**Supplementary Table T6 | Cell-type-resolved cholesterol receptor enrichment..**

| **Cell type** | **CD36** | **SCARB1** | **LDLR** | **Total cholesterol-MR** |
| --- | --- | --- | --- | --- |
| Activated macrophage | 3.5× | 1.0× | 0.8× | high |
| Stroma/macrophage | 1.4× | 1.1× | 1.0× | mid |
| Endothelial | 0.9× | 1.2× | 1.1× | mid |
| Luminal/tumour | 0.7× | 0.9× | 1.0× | low |

*Cell-type-resolved cholesterol-receptor enrichment (fold over mean; Kruskal–Wallis H = 163.3, P < 10⁻²⁹). CD36 dominates in activated macrophages.*

**Supplementary Table T7 | PPI + TF validation of breast two-hop relay patterns..**

| **Relay motif (LR1 → LR2)** | **Intracellular path** | **PPI score** | **Count** |
| --- | --- | --- | --- |
| IFNG→IFNGR1 → ARF1→PLD2 | IFNGR1 → STAT1 → [TF] → ARF1 | 0.155 | 1 |
| CD46→JAG1 → IGF1→TRPV2 | JAG1 → NOTCH1 → EP300 → JUN → [TF] → IGF1 | 0.199 | 1 |
| TNFSF12→CD163 → IFNG→IFNGR2 | CD163 → CSNK2B → BTRC → NFKB1 → [TF] → IFNG | 0.056 | 2 |
| SEMA4D→CD72 → LTF→LRP11 | CD72 → SIGLEC10 → ESR1 → [TF] → LTF | 0.213 | 2 |
| LRP11→ARF1 | LRP11 → CD1B → USP22 → MYC → [TF] → ARF1 | 0.043 | 7 |
| ARF1→PLD2 → LTF→LRP11 | PLD2 → RAC2 → PAK1 → ESR1 → [TF] → LTF | 0.089 | 5 |

*Representative validated receptor → PPI → TF → ligand paths for breast two-hop relay motifs (99/133 = 74 % validated; STRING + TRRUST/DoRothEA).*

**Supplementary Table T8 | Cross-channel relay validation (Receptor → enzyme → metabolite)..**

| **Receptor** | **Metabolite** | **Enzyme** | **PPI path (R → … → enzyme)** | **Count** |
| --- | --- | --- | --- | --- |
| CD72 | Deoxyadenosine | NT5E (CD73) | CD72 → PTPN6 → CXCR4 → NT5E | 2 |
| IL15RA | Methionine | CBS | IL15RA → IL2RB → HGS → CBS | 4 |
| CD163 | Serine | PSAT1 | CD163 → CSNK2B → RBM39 → PSAT1 | 3 |
| PLD2 | Deoxyadenosine | NT5E (CD73) | PLD2 → MTOR → NT5E | 1 |
| PLD2 | Lysine | SLC25A15 | PLD2 → RPTOR → PDCD1 → SLC25A15 | 6 |
| NFASC | Cholesterol | GMPS | NFASC → HNRNPL → GMPS | 5 |

*Representative validated receptor → enzyme → metabolite paths for cross-channel relay motifs (28/37 = 76 % validated; scFEA module–gene–metabolite mapping).*

**Supplementary Table T9 | SCARF1 relay subnetwork (breast)..**

| **Relay-transit gene** | **n chains** | **n distinct LR pairs** | **Lead downstream pair** |
| --- | --- | --- | --- |
| SCARF1 | 19 | 12 | TNFSF12 → CD163 |
| CALR | 11 | 3 | CALR → SCARF1 |
| TNFSF12 | 8 | 2 | TNFSF12 → CD163 |
| CD163 | 6 | 2 | CD163 → VSIG4 |
| ADAMTS4 | 4 | 2 | ADAMTS4 → VCAN |

*SCARF1 relay subnetwork summary (breast; SCARF1 as R1, 19 chains; 48 chains total across all relay-transit genes).*

**Supplementary Table T10 | Hub-gene enrichment terms (Enrichr, 50 breast hub genes)..**

| **Database** | **Top enriched term** | **Overlap** | **Adj. P** |
| --- | --- | --- | --- |
| GO BP | Cell-Cell Adhesion Via Plasma-Membrane Adhesion | 15/172 | 1.4×10⁻¹⁶ |
| KEGG | Cell adhesion molecules | 15/148 | 1.8×10⁻¹⁸ |
| Reactome | Cytokine Signaling In Immune System | 14/702 | 3.7×10⁻⁷ |

*Enrichr (Chen et al. 2013) enrichment of the 50 breast hub genes (top-decile hub-score; background = 1409 graph genes). Cell-adhesion is the single strongest enrichment; top term per library shown (Overlap = hub genes in term / term size).*

**Supplementary Table T11 | LR-network-hub KO summary (16 genes × 6 metrics).**

| **Gene** | **Tissue** | **n LR pairs** | **Attn removed** | **ΔLR loss %** | **ΔR²** | **% spots affected** |
| --- | --- | --- | --- | --- | --- | --- |
| LGALS3 | Breast | 31 | 13.84 | 3.36 | +0.0000 | 45.7 |
| NPTN | Breast | 10 | 9.06 | 0.62 | +0.0000 | 16.1 |
| APOA1 | Breast | 23 | 7.81 | 0.19 | +0.0000 | 0.4 |
| PIK3CB | Breast | 65 | 10.11 | 2.37 | +0.0000 | 32.1 |
| SLURP2 | Breast | 15 | 7.11 | 0.08 | +0.0000 | 0.7 |
| SLURP1 | Breast | 15 | 7.07 | 0.24 | +0.0000 | 4.7 |
| CDH2 | Breast | 29 | 8.75 | 0.90 | +0.0000 | 4.8 |
| PCDHGB1 | Breast | 11 | 8.00 | 0.59 | +0.0000 | 4.4 |
| GNAS | Brain | 18 | 10.70 | 10.34 | +0.0000 | 94.0 |
| ADAM10 | Brain | 21 | 8.72 | 1.99 | +0.0000 | 50.3 |
| IL16 | Brain | 10 | 7.13 | 1.11 | +0.0000 | 22.2 |
| EFNB1 | Brain | 18 | 6.55 | 0.05 | +0.0000 | 0.4 |
| APP | Brain | 36 | 9.76 | 7.97 | +0.0000 | 98.6 |

*Per-gene in-silico knockout effects (ε₂ LR-channel ablation). ΔLR loss % = Eq. (21) total-loss percentage; ΔR² = change in held-out expression R².*

**Supplementary Table T12 | SCARF1 fan-out × tier × literature evidence.**

| **LR pair** | **Tier** | **Lead compound** | **Trial / PMID** | **OT phase** | **Mechanism note** |
| --- | --- | --- | --- | --- | --- |
| TNFSF12 → CD163 | Clinical | enavatuzumab | NCT01383733 | 2 | anti-TWEAK trial; CD163 link inferential |
| AGT → ANPEP | Clinical | tosedostat | NCT01010373 | 2 | aminopeptidase inhibitor; AGT side not the trial target |
| C3 → CR2 | Clinical | pegcetacoplan | NCT04919629 | 2 | complement C3 inhibitor; CR2 link inferential |
| FBN1 → EBP | Clinical | tamoxifen | approved | 4 | tamoxifen/EBP indirect; FBN1 not the SERM target |
| APP → NFASC | Clinical | nirogacestat | NCT03785964 | 4 | γ-secretase clips APP; NFASC link inferential |
| ARF1 → PLD2 ★ | Preclinical | 36918511 | PMID 36918511 | 0 | PLD2 cancer growth; preclinical only |
| RARRES2 → CMKLR1 ★ | Preclinical | 35421316 | PMID 35421316 | 0 | chemerin–CMKLR1 axis; preclinical |
| LTF → LRP11 | Preclinical | 36195716 | PMID 36195716 | 0 | bovine LF CRC trial not via LRP11; gene-level only |
| PSAP → GPR37 ★ | Preclinical | 40032834 | PMID 40032834 | 0 | Sci Adv 2025 gastric PNI mechanism |
| CD99 → PILRA ★ | Preclinical | 40226493 | PMID 40226493 | 0 | Wang 2025 Nat Cancer; emerging checkpoint |
| CCL5 → GPR75 | Novel | — | — | 0 | GPR75 orphan; CCL5-GPR75 link contested |
| CCL19 → CCRL2 | Novel | — | — | 0 | CCRL2 atypical; chemerin > CCL19 |

*Translational evidence for the 12 SCARF1 downstream fan-out LR pairs (the SCARF1 multiplexer of Fig. 6h, spanning 19 relay chains; verified May 2026). Pairs are ordered by tier (clinical → preclinical → novel). OT phase = OpenTargets maximum clinical phase; ★ marks pairs whose lead compound directly antagonises that LR axis.*

**Supplementary Table T13 | Complete ablation and hyperparameter sensitivity grid (27 configurations)..**

| **Tier** | **ID** | **Configuration** | **Test R²** | **ΔR²** | **OP AUROC** | **ΔAUROC** | **CDB AUROC** | **ΔCDB** | **Params** |
| --- | --- | --- | --- | --- | --- | --- | --- | --- | --- |
| 1 | B2 | Remove ε₂ LR edges only | 0.4296 | −0.002 | 0.5000 | −0.240 | 0.5000 | −0.226 | 7,018,708 |
| 1 | B1 | Remove gene subgraph (gene nodes + ε₁ + ε₂) | 0.4270 | −0.005 | 0.5000 | −0.240 | 0.5000 | −0.226 | 5,108,178 |
| 2 | H2b | lr = 1e-2 (training divergence) | −0.0650 | −0.497 | 0.7385 | −0.002 | 0.7291 | +0.003 | 7,548,118 |
| 2 | H2a | lr = 1e-4 (under-fit) | 0.3283 | −0.104 | 0.7374 | −0.003 | 0.7298 | +0.004 | 7,548,118 |
| 2 | B14 | Replace scVI cell features with randn | 0.3246 | −0.107 | 0.7382 | −0.002 | 0.7306 | +0.004 | 7,548,118 |
| 3 | H1a | hidden_dim = 128 | 0.4219 | −0.010 | 0.7384 | −0.002 | 0.7290 | +0.003 | 2,134,418 |
| 3 | B16 | n_layers = 3 | 0.4264 | −0.006 | 0.7382 | −0.002 | 0.7310 | +0.005 | 11,005,149 |
| 3 | B12 | Remove ε₁ (cell → gene) only | 0.4267 | −0.005 | 0.7383 | −0.002 | 0.7295 | +0.003 | 7,018,708 |
| 3 | B6 | Zero scFEA flux edge attributes | 0.4271 | −0.005 | 0.7380 | −0.002 | 0.7311 | +0.005 | 7,548,118 |
| 3 | B4 | Remove ε₄ (metabolite → gene) only | 0.4278 | −0.004 | 0.7383 | −0.002 | 0.7311 | +0.005 | 7,019,732 |
| 3 | B18 | ffn_ratio = 2 (compact FFN) | 0.4279 | −0.004 | 0.7390 | −0.001 | 0.7320 | +0.006 | 5,972,182 |
| 3 | H1b | hidden_dim = 512 | 0.4281 | −0.004 | 0.7369 | −0.003 | 0.7289 | +0.003 | 23,127,818 |
| 3 | H3a | dropout = 0.0 | 0.4253 | −0.007 | 0.7393 | −0.001 | 0.7321 | +0.006 | 7,548,118 |
| 3 | B7 | Uniform cross-edge gate (frozen logits) | 0.4283 | −0.004 | 0.7375 | −0.003 | 0.7300 | +0.004 | 7,548,118 |
| 3 | B17 | n_heads = 8 (head_dim = 32) | 0.4301 | −0.002 | 0.7368 | −0.003 | 0.7302 | +0.004 | 7,548,118 |
| 3 | B10 | Remove τ₂ (cell-cell metabolite) only | 0.4310 | −0.001 | 0.7375 | −0.003 | 0.7304 | +0.004 | 7,018,708 |
| 3 | H3b | dropout = 0.2 | 0.4339 | +0.002 | 0.7374 | −0.003 | 0.7287 | +0.002 | 7,548,118 |
| 3 | B8 | n_layers = 1 | 0.4323 | +0.000 | 0.7387 | −0.001 | 0.7309 | +0.005 | 4,091,087 |
| 3 | B9 | Remove τ₁ (cell-cell secreted) only | 0.4325 | +0.001 | 0.7386 | −0.002 | 0.7318 | +0.006 | 7,019,220 |
| 3 | B3 | Remove metabolite subgraph (all metab edges + nodes) | 0.4331 | +0.001 | OOM† | — | OOM† | — | 5,282,770 |
| 3 | B5 | Replace ESM-2 with random gene features | 0.4332 | +0.001 | 0.7378 | −0.002 | 0.7303 | +0.004 | 7,548,118 |
| 3 | B21 | Remove β-gated lane residual (force β ≈ 0 in all lanes) | 0.4360 | +0.004 | 0.7365 | −0.004 | 0.7292 | +0.003 | 7,548,118 |

*27-configuration leave-one-out ablation + hyperparameter sweep on breast_new. Bold = catastrophic / R²-critical. Default: R² = 0.4319, OmniPath AUROC = 0.7401.*

**Supplementary Table T14 | Trainable parameter accounting..**

| **Block** | **Equation** | **Instances** | **Trainable parameters** |
| --- | --- | --- | --- |
| Per-node-type input projection MLP | Eq. (1) | 3 (cell, gene, metabolite) | 516,352 |
| HetGT attention bank (7 lanes × Q/K/V/skip/β-gate/edge-attr) | Eqs. (2)–(7) | 2 stacked blocks | 3,754,496 |
| Cross-edge softmax gate + Add+LN | Eqs. (8)–(9) | 2 stacked blocks | 3,086 |
| FFN + Add+LN per node type | Eqs. (10)–(11) | 2 stacked blocks | 3,156,480 |
| Expression decoder | Eq. (12) | 1 | 117,704 |
| Total trainable |  |  | 7,548,118 |

*Trainable-parameter accounting for the 7,548,118-parameter baseline encoder. Foundation models are frozen; read-out heads carry zero parameters.*

**Supplementary Notes**

**Supplementary Note N1. Bootstrap confidence-interval methodology.**

For every (method, evaluation) AUROC in the benchmark, we report a 95 % bootstrap confidence interval computed by resampling the positive and negative ranked-pair sets with replacement (2,000 resamples). The resampling preserves the true class balance per evaluation. CIs for the AUROC difference (Supplementary Fig. S3) are computed as the difference of the resampled AUROCs for the two methods on the same resample, paired, this accounts for dependence between method scores on shared pairs. The Wilcoxon signed-rank test on the eight per-evaluation AUROC differences yields P = 0.004 (n = 8) and Cohen's d = 2.35.

**Supplementary Note N2. Leiden / k-means clustering parameters and coherence metric.**

Clustering parameters. The breast and mouse-cortex niches in Fig. 3a, b are obtained by k-means (random_state = 42, n_init = 10; k = 17 and k = 26 respectively) on MOSANIC's learned cell embedding. The intestinal, lung and hippocampus niches in Supplementary Fig. S9 are obtained by Leiden clustering of the embedding k-nearest-neighbour graph (resolution 1.0–1.5, target k ≈ 15), giving 21, 25 and 16 niches respectively. Coherence metric. For the per-dataset spatial maps (Supplementary Fig. S9), spatial coherence C is the fraction of each cell's k = 15 nearest spatial neighbours that share its niche label: C = (1 / N) × Σᵢ (1 / k) × |{j ∈ NN₁₅(i): labelⱼ = labelᵢ}|. This spatial-neighbour metric is symmetric in labels and applies to both coarse (5 cell types, mouse cortex) and fine (17 niches, breast) partitions. For the niche partitions in Fig. 3a, b we additionally report cluster purity, the mean over niches of the largest single-cell-type fraction, which quantifies how cell-type-pure each learned niche is: breast niches reach purity 0.656 and cortex niches 0.949, versus 0.236 and 0.476 for random partitions of the same granularity (1,000 label permutations).

**Supplementary Note N3. Per-cell MR intensity definition.**

MR intensity per pair (M, R) in cell i is Iᵢ^MR(M, R) = α^ε₄_MR · |fᵢ_M| · max(xᵢ_R, 0) where α^ε₄_MR is the model's learned ε₄ attention weight for the (metabolite, receptor) edge, fᵢ_M is the scFEA-predicted flux of metabolite M in cell i (absolute value because both production and consumption indicate activity), and xᵢ_R is log-normalised expression of receptor gene R in cell i. Total per-cell MR intensity is the sum across all ε₄ edges.

**Supplementary Note N4. scFEA flux inference and MR database.**

Metabolic flux was inferred with scFEA (Alghamdi et al., Genome Res. 2021) using the Human_M168 model (168 reaction modules spanning 70 metabolite compounds; 100 epochs on log-normalised expression). Output: a per-cell metabolite-balance matrix [N × 70]. MR database: 794 human (790 mouse) curated metabolite–receptor pairs (MEBOCOST; Chen et al. 2023) supplemented with canonical neurotransmitter receptor assignments. The LIANA+ metabolite pipeline used for Fig. 4e is rank-aggregate scoring (CellPhoneDB + SingleCellSignalR, expression-proportion threshold 0.0).

**Supplementary Note N5. Relay-chain detection algorithm and intracellular validation.**

Detection. A two-hop relay chain A→B→C requires: (i) edges A→B and B→C both in the top 10 % of per-channel attention (ε₂ for secreted, ε₄ for metabolite); (ii) cell B a shared spatial neighbour of both A and C; (iii) the best-matching LR pair for A→B distinct from that for B→C, ensuring different molecular mechanisms at each hop; and (iv) relay score τ = attn(A→B) × attn(B→C) exceeding the dataset-specific threshold from 1,000 permutation nulls (edge weights shuffled, spatial structure preserved). Cross-channel chains additionally require hop 1 on ε₂ and hop 2 on ε₄. To prioritise high-confidence events we apply conservative enumeration caps, top-300 hub cells by out-degree, top-5 incoming edges per hub, best-ranked LR pair per edge from the model's top-100 LR scores, yielding 208 (breast) and 135 (brain) two-hop chains before ranking; exhaustive enumeration without caps recovers 646 and 618, with identical unique patterns and validation rates. Intracellular validation. For each unique motif the shortest weighted path Receptor₁ → [PPI intermediates] → TF → Ligand₂ is queried against STRING signalling PPIs (584,347 edges; experimental score ≥ 0.1) and TRRUST + DoRothEA TF-target tables (276,731 edges), with PPI thresholds swept 0.5→0.1 and the highest-confidence valid path retained. For cross-channel motifs the TF step is replaced by a metabolic-enzyme step using the scFEA module–gene–metabolite mapping. Validation rates: breast two-hop 99/133 (74.4 %), cross-channel 28/37 (75.7 %), brain two-hop 36/36 (100 %); the higher brain rate reflects densely-connected GNAS/ADCY intermediates. All unvalidated motifs fail only because Receptor₁ is absent from the STRING signalling network (Supplementary Tables T7, T8). Significance of the per-hop τ decay (Mann–Whitney on raw distributions): 1-hop→2-hop P = 3 × 10⁻⁶⁶, 2-hop→3-hop P = 2 × 10⁻¹². Comparison with CellNEST. CellNEST (Zohora et al. 2025) introduced relay detection but connects each cell to all neighbours within 4× spot spacing (mean degree 80.3, 202,139 breast edges), so at top-10 % attention it retains 8.6 edges per active cell across 93.6 % of cells in one component, designating 87 % of cells as potential hubs. MOSANIC uses k = 6 neighbours within 150 µm (15,060 edges, 13.4× fewer), retaining 1.7 edges across 34.5 % of cells and resolving 1,177 structured relay communities. Both methods share the PPI+TF validation infrastructure; MOSANIC extends the concept to multi-channel, cross-modal relays.

**Supplementary Note N6. Literature validation of the relay predictions.**

Post-hoc mining confirms that every major relay prediction is consistent with independent published experimental evidence, recovered from attention topology without pathway or receptor-function annotation. - TNFSF12→CD163. CD163 is a direct TWEAK receptor on monocytes (Bover et al. 2007, J. Immunol. 178:8183); TWEAK drives macrophage accumulation via MCP-1 (Dwyer et al. 2021, J. Hepatol. 74:860); the clinical anti-TWEAK-receptor antibody enavatuzumab confirms therapeutic relevance (Lam et al. 2018, Mol. Cancer Ther. 17:215). - CALR→SCARF1. Calreticulin is the dominant pro-phagocytic "eat-me" signal on human cancers (Chao et al. 2010, Sci. Transl. Med. 2:63ra94); SCARF1 mediates apoptotic-cell clearance and SCARF1-null mice develop lupus-like autoimmunity (Ramirez-Ortiz et al. 2013, Nat. Immunol. 14:917). - Cholesterol→CD36. CD36-mediated lipid uptake is required for TAM differentiation (Su et al. 2020, Cancer Res. 80:1438); membrane cholesterol efflux drives TAM reprogramming (Goossens et al. 2019, Cell Metab. 29:1376); CD36-oxidised-lipid uptake promotes T-cell dysfunction (Xu et al. 2021, Immunity 54:1561). - SEMA4D→CD72. SEMA4D is produced by TAMs and drives tumour angiogenesis (Sierra et al. 2008, J. Exp. Med. 205:1673); the anti-SEMA4D antibody pepinemab promotes immune infiltration (Evans et al. 2015, Cancer Immunol. Res. 3:689). The validated path CD72→PTPN6→CXCR4→NT5E/CD73 connects to the adenosine immunosuppressive axis (18 clinical mAbs). - Methionine→SLC43A1. The related transporter SLC43A2 mediates methionine competition between tumour and T cells, causing immune evasion (Bian et al. 2020, Nature 585:277). - GNAS→ADCY1 (brain). ADCY1 knockout disrupts cortical barrel maps (Abdel-Majid et al. 1998, Nat. Genet. 19:289); GNAS haploinsufficiency causes cognitive impairment (Turan & Bastepe 2015, Curr. Osteoporos. Rep. 13:146); the one-Gαs-to-many-AC fan-out is textbook G-protein signalling (Sadana & Dessauer 2009). - AGRN (brain). Agrin-deficient cortex loses 30 % of excitatory (not inhibitory) synapses (Ksiazek et al. 2007, J. Neurosci. 27:7183). These concordances establish that relay chains preferentially involve genes with established intercellular-communication functions, and that the cross-channel and tissue-specific relay biology the model surfaces is real signalling rather than a graph artefact.

**Supplementary Note N7. In-silico knockout by attention-edge ablation: method & assumptions**

**Supplementary Note N8. Tier classification of LR pairs (clinical / preclinical / novel), sources & rules**

**Supplementary Note N9. The small Tier-3 Δs are genuine, not a metric artefact: intensity-weighted discriminative metrics.**

The OmniPath ΔAUROC values across Tier 3 (≤ 0.005) and the ConnectomeDB ΔAUROC values (≤ 0.006) are conspicuously small. We tested whether this reflects a coarse metric that masks top-K ranking differences or a genuinely robust model. Saturation diagnostic. In the breast LR graph there are 9,571 ε₂ edges, of which 183 (≈ 2 %) are saturated at ᾱlr ≥ 0.999 because the corresponding receptor has in-degree 1 (per-destination softmax assigns probability 1 to the single incoming ligand). Any model that runs forward over these edges produces the same saturated set, so a top-K ranking on raw attention is identical across all 13 non-catastrophic configurations (Top-100 Jaccard ≡ 1.000 against the baseline). AUROC against OmniPath / ConnectomeDB therefore captures only the non-saturated (9,388) tail, which differs at most in the third-to-fourth decimal across ablations. The metric is correct, small AUROC ΔΔs are reproducible across random seeds, but it is intrinsically coarse for this graph. Intensity-weighted recomputation. The canonical user-facing score is the intensity-weighted variant Ī(l, r) = ᾱlr · xl+ · xr+ (Methods Eq. 15), which uses per-gene mean expression to break the degree-1 saturation. With this score the 183 previously-tied saturated pairs each resolve to a distinct value (4,651 unique scores across the vocabulary), making Top-K Jaccard a meaningful sensitivity probe. We recomputed Spearman ρ vs the baseline ranking, Top-K Jaccard at K = 20 / 50 / 100, Lift@100 for both reference databases, and AUROC under intensity-weighted scoring:

| **ID** | **Configuration** | **Spearman ρ** | **Top-20 J** | **Top-50 J** | **Top-100 J** | **Lift@100 OP** | **AUROC OP (intensity)** |
| --- | --- | --- | --- | --- | --- | --- | --- |
| baseline | (full model) | 1.0000 | 1.000 | 1.000 | 1.000 | 1.367 | 0.6902 |
| B2 | no ε₂ (catastrophe) | (n/a) | 0.000 | 0.000 | 0.000 | 0.557 | 0.5000 |
| B14 | random scVI | 0.9997 | 1.000 | 1.000 | 0.905 | 1.417 | 0.6902 |
| B7 | uniform gate | 0.9997 | 1.000 | 1.000 | 0.923 | 1.417 | 0.6900 |
| B5 | random ESM-2 | 0.9989 | 1.000 | 1.000 | 0.923 | 1.468 | 0.6899 |
| B10 | no τ₂ | 0.9997 | 1.000 | 1.000 | 0.942 | 1.265 | 0.6901 |
| B12 | no ε₁ | 0.9996 | 1.000 | 1.000 | 0.942 | 1.265 | 0.6898 |
| B18 | ffn_ratio = 2 | 0.9993 | 1.000 | 1.000 | 0.942 | 1.417 | 0.6906 |
| B21 | no β-gated residual | 0.9997 | 1.000 | 1.000 | 0.942 | 1.367 | 0.6896 |
| B16 | n_layers = 3 | 0.9997 | 1.000 | 1.000 | 0.961 | 1.367 | 0.6901 |
| B17 | n_heads = 8 | 0.9998 | 1.000 | 1.000 | 0.961 | 1.316 | 0.6903 |
| B8 | n_layers = 1 | 0.9997 | 1.000 | 1.000 | 0.961 | 1.367 | 0.6903 |
| B9 | no τ₁ | 0.9996 | 1.000 | 1.000 | 0.980 | 1.367 | 0.6904 |
| B6 | no flux | 0.9998 | 1.000 | 1.000 | 0.980 | 1.417 | 0.6904 |
| B4 | no ε₄ | 0.9998 | 1.000 | 1.000 | 0.980 | 1.367 | 0.6905 |

(13 non-catastrophic configurations shown.) Interpretation. Top-20 and Top-50 Jaccard ≡ 1.000 in every non-catastrophic configuration: the highest-confidence LR predictions (the 20–50 pairs reported in the main figures) are invariant across every Tier-3 modification. Top-100 Jaccard ranges 0.90–0.98; the most-perturbed are B14 (random scVI), B7 (uniform gate) and B5 (random ESM-2), modifications that disrupt features or attention shaping, confirming the rank perturbation that AUROC partially captures is real. Spearman ρ ≥ 0.999 confirms global rank similarity. The small AUROC ΔΔs in Table T13 are not a metric artefact; they reflect a model that produces a nearly-invariant ranking across encoder perturbations. The primary metrics in T13 (R², OmniPath AUROC, ConnectomeDB AUROC) correctly capture the three-tier conclusion.

**Supplementary Note N10. Literature concordance of the in-silico knockout predictions**
